# Engineered human cytokine/antibody fusion proteins expand regulatory T cells and confer autoimmune disease protection

**DOI:** 10.1101/2022.05.29.493918

**Authors:** Derek VanDyke, Marcos Iglesias, Jakub Tomala, Arabella Young, Jennifer Smith, Joseph A. Perry, Edward Gebara, Amy R. Cross, Laurene S. Cheung, Arbor G. Dykema, Brian T. Orcutt-Jahns, Tereza Henclová, Jaroslav Golias, Jared Balolong, Luke M. Tomasovic, David Funda, Aaron S. Meyer, Drew M. Pardoll, Joanna Hester, Fadi Issa, Christopher A. Hunter, Mark S. Anderson, Jeffrey A. Bluestone, Giorgio Raimondi, Jamie B Spangler

## Abstract

Low dose human interleukin-2 (hIL-2) treatment is used clinically to treat autoimmune disorders due to the cytokine’s preferential expansion of immunosuppressive regulatory T cells (T_Reg_s). However, high toxicity, short serum half-life, and off-target immune cell activation limit the clinical potential of IL-2 treatment. Recent work showed that complexes comprising hIL-2 and the anti-hIL-2 antibody F5111 overcome these limitations by preferentially stimulating T_Reg_s over immune effector cells. Although promising, therapeutic translation of this approach is complicated by the need to optimize dosing ratios and by the instability of the cytokine/antibody complex. We leveraged structural insights to engineer a single-chain hIL-2/F5111 antibody fusion protein, termed F5111 immunocytokine (IC), that potently and selectively activates and expands T_Reg_s. F5111 IC conferred protection in mouse models of colitis and checkpoint inhibitor-induced diabetes mellitus. These results provide a roadmap for IC design and establish a T_Reg_-biased immunotherapy that could be clinically translated for autoimmune disease treatment.

## Introduction

IL-2 is a pleiotropic cytokine that regulates key functions in maintaining immune homeostasis, including promotion of the proliferation, survival, and activation of both pro-inflammatory immune effector cells (e.g., CD4^+^/CD8^+^ effector T cells, natural killer [NK] cells) and anti-inflammatory regulatory T cells (T_Reg_s). IL-2 mediates its activities by forming complexes with membrane-embedded receptors that orchestrate signaling through the Janus kinase (JAK)-signal transducer and activator of transcription (STAT) pathway to influence immune-related gene expression and regulate functional outcomes (Malek, 2008; Stroud and Wells, 2004; Murray, 2007). IL-2 forms either an intermediate-affinity (equilibrium dissociation constant [K_D_]≈1 nM) heterodimeric receptor complex, consisting of the IL-2 receptor-β (IL-2Rβ, CD122) and common gamma (γ_c_, CD132) chains, or a high-affinity (K_D_≈10 pM) heterotrimeric receptor complex, consisting of the non-signaling IL-2 receptor-α subunit (IL-2Rα, CD25) in addition to the IL-2Rβ and γ_c_ chains (Boyman and Sprent, 2012; Liao et al., 2013; Malek, 2008; Wang et al., 2005). The IL-2Rα subunit is highly expressed on T_Reg_s but virtually absent from naïve immune effector cells, thus rendering T_Reg_s 100-fold more sensitive to IL-2 (Boyman and Sprent, 2012; Malek, 2008; Sakaguchi et al., 1995; Spangler et al., 2015a; Taniguchi and Minami, 1993). IL-2Rα expression is also induced in activated immune effector cells, and although expression levels are lower than those of T_Reg_s, activated immune effector cells may compete with T_Reg_s for extracellular IL-2 (Baecher-Allan et al., 2001; Höfer et al., 2012; Sakaguchi et al., 1995; Schmidt et al., 2012). Hence, IL-2 can promote both pro-and anti-inflammatory responses, which has made it an attractive, albeit complex, candidate for immunotherapy.

Exploiting the cytokine’s pro-inflammatory activities, high-dose IL-2 treatment has received FDA approval for treatment of renal cell carcinoma and metastatic melanoma (Alva et al., 2016; Rosenberg, 2014; Sim and Radvanyi, 2014). However, the essential role of IL-2 as a growth factor for T_Reg_s has complicated treatment efficacy, and high-dose IL-2 therapy can lead to toxic side effects such as vascular leak syndrome, further limiting its clinical usage (Alva et al., 2016; Sim and Radvanyi, 2014). In contrast, the anti-inflammatory activities of IL-2 have been exploited by administering low doses of the cytokine to treat autoimmune conditions such as diabetes, graft-versus-host disease, allograft rejection, and ulcerative colitis (Allegretti et al., 2021; Cheng et al., 2011; Grinberg-Bleyer et al., 2010; Klatzmann and Abbas, 2015; Koreth et al., 2011; Saadoun et al., 2011; Shevach, 2012; Tahvildari and Dana, 2019). However, this approach has encountered limitations including difficulties in dosing optimization due to the short *in vivo* half-life of IL-2 and dangerous off-target effects that result from activation of immune effector cells (Dong et al., 2021; Klatzmann and Abbas, 2015). Therefore, striking a balance between efficacy, safety, and reduced off-target activation remains a challenge in realizing the clinical potential of IL-2 treatment, and isolation of the cytokine’s immunosuppressive activities would greatly advance autoimmune disease therapy and transplantation medicine.

Several approaches have been explored to overcome the limitations of IL-2 therapy, including design of IL-2 muteins (Peterson et al., 2018; Carmenate et al., 2018; Khoryati et al., 2020; Glassman et al., 2021), PEGylated IL-2 variants (Charych et al., 2016; Dixit et al., 2021; Zhang et al., 2021), and IL-2/IL-2Rα fusion proteins (Hernandez et al., 2021; Lopes et al., 2020; Ward et al., 2018, 2020; Xie et al., 2021). Another approach, pioneered by Boyman et al., is the formation of complexes between IL-2 and anti-IL-2 antibodies, which has been shown to increase therapeutic efficacy and reduce toxicity of the cytokine by both extending its *in vivo* half-life and selectively targeting its functions towards particular immune cell subsets through structural and molecular mechanisms (Arenas-Ramirez et al., 2016; Boyman and Sprent, 2012; Boyman et al., 2006b, 2006a; De Paula et al., 2020; Karakus et al., 2020; Lee et al., 2020; Spangler et al., 2015b; Trotta et al., 2018). In a recent example, Trotta and colleagues identified a human monoclonal antibody (F5111) against hIL-2, that sterically blocks cytokine binding to the IL-2Rβ subunit and also allosterically reduces the cytokine’s affinity for IL-2Rα, which leads to preferential activation of T_Reg_s (Trotta et al., 2018). When bound to hIL-2, F5111 occludes the IL-2Rβ binding site and causes perturbations in the AB and BC loops of IL-2 that propagate towards the IL-2Rα binding site to disfavor IL-2Rα binding, resulting in an advantage for IL-2Rα^High^ T_Reg_s. Additionally, since the F5111 antibody must be released in order for IL-2 to activate the IL-2Rβ/γ_c_ heterodimeric receptor, it is speculated that interaction of the IL-2/F5111 complex with IL-2Rα may destabilize the cytokine/antibody complex and increase availability of the IL-2Rβ binding site on the cytokine, thus promoting selective activation of the IL-2 signaling on cells that are IL-2Rα^High^ (i.e., T_Reg_s). This paradigm resembles the exchange/release mechanism observed for the anti-mouse IL-2 (mIL-2) antibody, JES6-1, and the anti-hIL-2 antibody UFKA-20, both of which preferentially stimulate T_Reg_s when complexed with IL-2 (Boyman et al., 2006a; Karakus et al., 2020; Spangler et al., 2015b). The F5111 antibody was affinity matured towards hIL-2 to create the F5111.2 antibody, which was found to direct IL-2 towards preferential expansion of T_Reg_s and ameliorate autoimmune diseases in mice (Trotta et al., 2018). Although this approach is promising, clinical translation of cytokine/antibody complexes is complicated by the need for optimization of dosing ratios and by the instability of the complex, as dissociation leads to off-target effects and rapid clearance of the unbound IL-2.

To overcome the limitations in clinical development of IL-2/anti-IL-2 antibody complexes, the IL-2 cytokine has been genetically fused to anti-IL-2 antibodies (Sahin et al., 2020; Spangler et al., 2018; Tomala et al., 2013). In one example, a single-chain fusion protein (immunocytokine, IC) comprising mIL-2 and the JES6-1 antibody was engineered, and it was demonstrated that the IC led to superior disease control compared to the IL-2/JES6-1 complex in a mouse model of colitis (Spangler et al., 2018). Here, we used a structure-based engineering approach to design an IC that links hIL-2 to the F5111 antibody (termed F5111 IC) (Spangler and VanDyke, 2020). We showed that F5111 IC preferentially promoted both mouse and human T_Reg_ activation and expansion *ex vivo* and *in vivo*. By modulating the cytokine/antibody affinity, we developed a panel of IC variants that exhibited a range of potencies on different immune cell subsets. Interestingly, we found that amongst all variants, our original F5111 IC construct was poised at a unique affinity optimum and thus demonstrated an exquisitely selective T_Reg_ bias that exceeded the bias of the hIL-2/F5111.2 complex. Finally, we established the therapeutic promise of F5111 IC by showing efficacy in mouse models of colitis and immune checkpoint inhibitor-induced diabetes mellitus.

## Results

### Immunocytokine design and optimization

To advance development of F5111 IC, we first produced the F5111 human immunoglobulin G1 (IgG1) antibody (Trotta et al., 2018) from a mammalian expression system. Binding studies using the yeast surface display platform (Boder and Wittrup, 1997; Chao et al., 2006) revealed that our recombinantly expressed F5111 bound yeast-displayed hIL-2 with similar affinity to literature reports (Trotta et al., 2018). We also found that the antibody was not cross reactive with mIL-2 (**Figure S1A**).

F5111 IC was constructed by tethering hIL-2 to the N-terminus of the light chain (LC) of the full-length human F5111 antibody with a flexible linker (**Figure 1A**). Since the mechanism of action for biased immune activation by hIL-2/F5111.2 complexes requires dissociation of the cytokine from the antibody (Trotta et al., 2018), we hypothesized that tethering the cytokine to the antibody could hinder this dissociation through enhanced avidity effects (Spangler et al., 2018). Therefore, in order to reduce the intramolecular cytokine/antibody affinity and thereby ensure proper release of hIL-2 from the antibody, we chose to formulate the F5111 IC using F5111 rather than the affinity matured antibody variant F5111.2 (Trotta et al., 2018). Based on the published molecular structure of hIL-2/F5111 complex (Trotta et al., 2018), the N-terminus of the F5111 LC is a distance of 43 Å from the C-terminus of hIL-2 (**Figure 1B**); therefore our initial design consisted of a 15 amino acid (Gly_4_Ser)_3_ flexible linker to allow for intramolecular cytokine engagement. We refer to this initial design as F5111 IC LN15. To allow for additional flexibility, constructs were also prepared using a 25 amino acid (Gly_4_Ser)_5_ linker (F5111 IC LN25) and a 35 amino acid (Gly_4_Ser)_7_ linker (F5111 IC LN35). A control immunocytokine (Control IC) representing bivalent, IgG-fused IL-2 was also constructed by replacing the variable heavy (V_H_) and variable light (V_L_) chains of the F5111 IC LN35 construct with those of an irrelevant anti-fluorescein isothiocyanate (FITC) antibody (Honegger et al., 2005) (**Table S1**).

**Figure 1.**
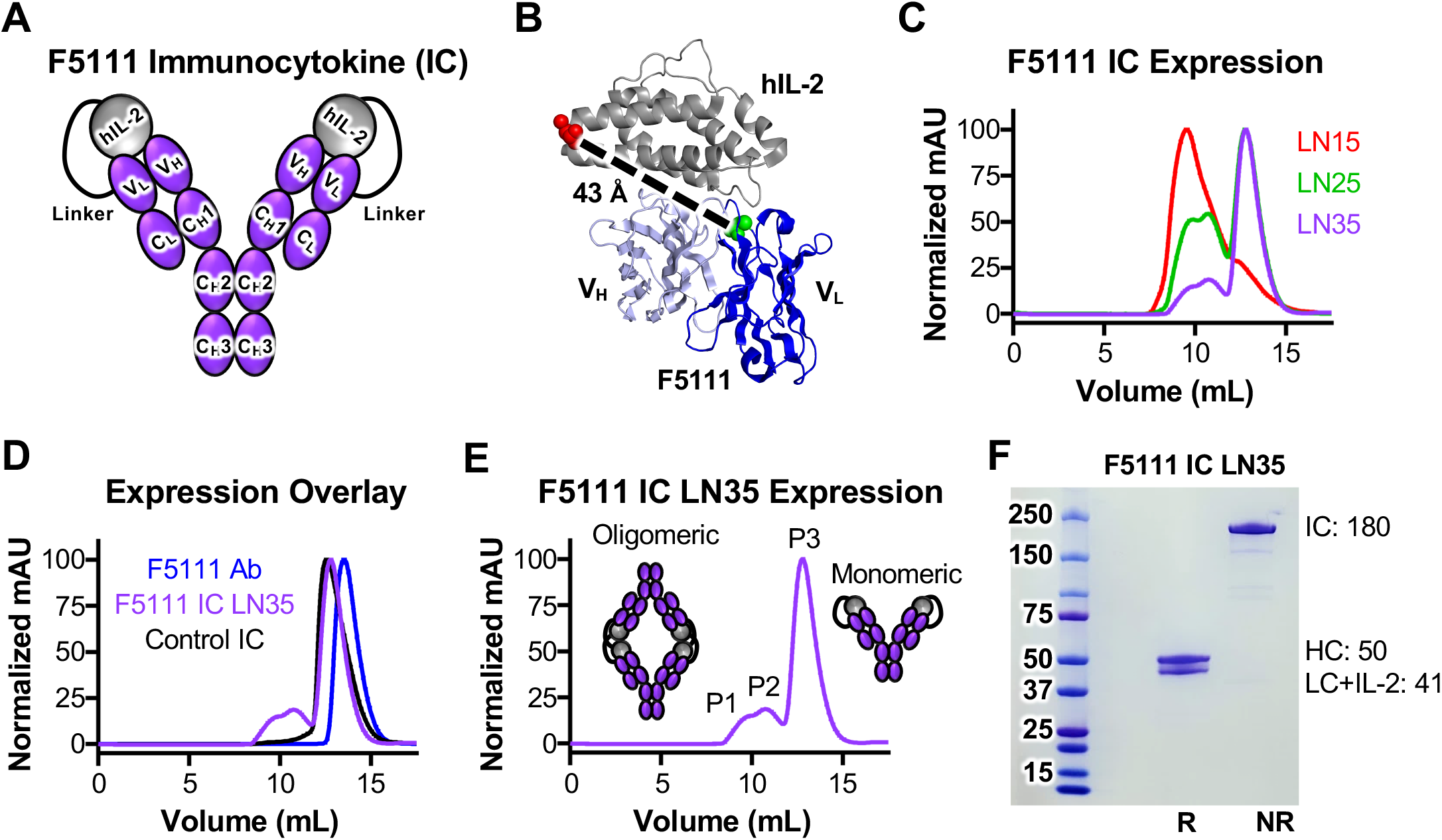
Design and production of F5111 IC. **(A)** Schematic of F5111 IC design. The C-terminus of hIL-2 is tethered to the N-terminus of the F5111 antibody light chain via a flexible amino acid linker consisting of Gly_4_Ser repeats. **(B)** Structure of the hIL-2/F5111 complex (PDB ID 5UTZ) (Trotta et al., 2018), illustrating the distance between the C-terminus of hIL-2 (red) and the N-terminus of the F5111 antibody light chain (green). **(C)** Overlay of the SEC traces of F5111 IC LN15, F5111 IC LN25, and F5111 IC LN35. **(D)** Overlay of the SEC traces of F5111 antibody (Ab), F5111 IC LN35, and Control IC. **(E)** SEC trace of F5111 IC LN35 with the oligomeric peaks (P1, P2) and the monomeric peak (P3) highlighted. **(F)** SDS-PAGE analysis of F5111 IC LN35 P3. NR = non-reducing. R = reducing. Molecular weights (in kDa) of IC, HC, and LC + IL-2 are indicated at right. See also **Figure S1; Table S1**.

ICs were expressed recombinantly through transient transfection of human embryonic kidney (HEK) 293F cells and purified via protein G affinity chromatography followed by size-exclusion chromatography (SEC). Compared to the F5111 antibody, which eluted as a single monodisperse peak, F5111 IC LN15 eluted as two broader peaks with the majority of the IC eluting earlier than the antibody (**Figures 1C and 1D**). This observation suggested possible oligomerization of F5111 IC LN15, prompting us to evaluate constructs with longer linkers. Analysis of F5111 IC LN25 and LN35 revealed three elution peaks (denoted P1, P2, and P3) (**Figures 1C-1E**). It was expected that due to their earlier elution volumes, P1 and P2 contained higher order oligomeric structures, whereas P3, contained the monomeric IC (**Figures 1D and 1E**). These higher order oligomers likely represent engagement of linked hIL-2 by F5111 on a neighboring IC (intermolecular assembly) rather than intramolecular assembly (**Figure 1E**). This presumably results from use of a shorter linker, which hinders intramolecular interaction between hIL-2 and F5111. This finding is supported by the fact that we saw less oligomerization as we increased the linker length between the antibody and cytokine. Additionally, Control IC (which does not assemble intramolecularly or intermolecularly) eluted as a single peak that coincided with P3 of F5111 IC LN25 and LN35, further suggesting that P3 indeed represents the monomeric IC (**Figure 1D**). Thus, henceforth, references to F5111 IC LN25 and F5111 IC LN35 indicate P3 unless otherwise specified. Also, references to F5111 IC LN15 indicate the majority peak (elution volume 10 mL, **Figure 1C**). SDS-PAGE verified the purity of ICs (**Figure 1F**).

### F5111 ICs are intramolecularly assembled and exhibit expected cytokine and receptor interactions

To confirm proper assembly of ICs and assess functionality, we characterized the binding of the F5111 ICs to immobilized hIL-2, human IL-2Rα (hIL-2Rα), and human IL-2Rβ (hIL-2Rβ) using bio-layer interferometry. For all binding experiments, hIL-2/F5111 was used at a 1:1 molar ratio to minimize presence of the free cytokine.

If properly assembled, F5111 would not be expected to engage immobilized hIL-2 since the hIL-2 within the IC is stably bound to the antibody. The F5111 antibody bound to hIL-2 with an affinity consistent with previous reports (Trotta et al., 2018), whereas the F5111 IC LN15, F5111 IC LN25, and F5111 IC LN35 all exhibited significantly impaired binding to the cytokine (**Figures 2A, left, S1C, left, and S1D, left**; **Table S2**), reflecting intramolecular assembly of the ICs. Notably, hIL-2 binding to ICs was detectable above background, suggesting transient exchange between intramolecular assembly of the IC and interaction of the component antibody with hIL-2. Binding to immobilized hIL-2 decreased as the linker length increased, reinforcing the observation that longer linker lengths stabilize proper assembly of the IC. The three peaks that eluted from F5111 IC LN25 showed impaired binding to hIL-2, and no major differences were observed between the peaks (**Figure S1E, left**). Binding was observed between hIL-2 and hIL-2/F5111 complex; however, this can be explained by the fact that a 1:1 molar ratio of hIL-2 to F5111 was used for binding studies, leaving unbound sites on the antibody available for interaction (**Figure 2A, left**). Taken together, binding studies against immobilized hIL-2 indicated that our lead IC construct (F5111 IC LN35) was stably assembled through intramolecular interactions.

**Figure 2.**
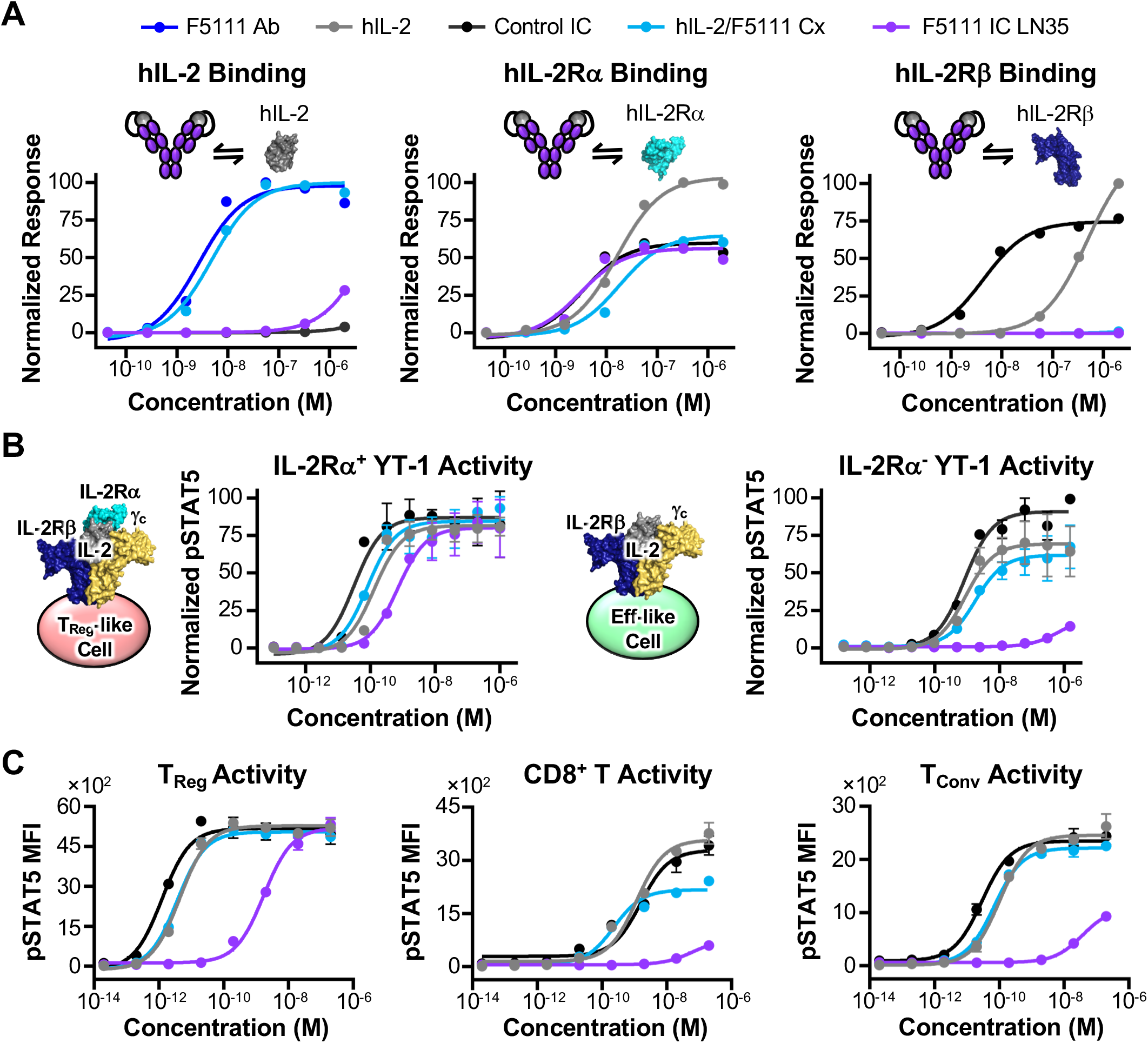
F5111 IC blocks IL-2 binding to IL-2Rβ and biases towards T_Reg_ activation. **(A)** Equilibrium biolayer interferometry-based titrations of F5111 antibody (Ab), hIL-2, Control IC, hIL-2/F5111 complex (Cx, 1:1 molar ratio), and F5111 IC LN35 binding to immobilized hIL-2 (left), immobilized hIL-2Rα (middle), and immobilized hIL-2Rβ (right). **(B)** STAT5 phosphorylation response of IL-2Rα^+^ (left) and IL-2Rα^-^ (right) YT-1 human NK cells stimulated with either hIL-2, Control IC, hIL-2/F5111 Cx (1:1 molar ratio), or F5111 IC LN35. **(C)** STAT5 phosphorylation response of human T_Reg_ (left), CD8^+^ T (middle), and T_Conv_ (right) cell populations within human PBMCs stimulated with either hIL-2, Control IC, hIL-2/F5111 Cx, or F5111 IC LN35. Data represent mean ± SD (n=3). See also Figures S1 and S2**; Tables S2-S4**.

To further interrogate the biophysical properties of F5111 ICs, we investigated their interactions with immobilized hIL-2Rα. We expected that hIL-2Rα would bind F5111 IC since the receptor-binding epitope on hIL-2 is not directly obstructed by the antibody (Trotta et al., 2018). F5111 ICs with varying linker lengths bound to hIL-2Rα with similar affinities, all of which were at least 2-fold higher affinity compared to free hIL-2 and hIL-2/F5111 complex, due to avidity-mediated enhancement resulting from bivalent cytokine presentation (**Figures 2A, middle, S1C, middle, and S1D, middle; Table S2**). Likewise, Control IC had a 5-fold higher affinity for hIL-2Rα compared to free IL-2 due to its bivalent construction (**Figure 2A, middle; Table S2**). A slightly reduced hIL-2Rα affinity (2-fold) was observed for F5111 IC LN15 as compared to F5111 IC LN25 and F5111 IC LN35, which was likely due to the observed oligomerization. Indeed, when we compared hIL-2Rα binding of the 3 peaks of F5111 IC LN25, we found that P3 had the highest affinity whereas P1 had the lowest (**Figure S1 E, middle**). These findings indicate that the monomeric F5111 IC functionally binds hIL-2Rα, as designed.

In contrast with hIL-2Rα, hIL-2Rβ should *not* bind F5111 IC since the antibody should fully block the receptor-binding epitope on hIL-2, as reported for hIL-2/F5111 complex (Trotta et al., 2018). Free hIL-2 bound to hIL-2Rβ with an expected affinity (Spangler et al., 2015b), and Control IC had a 100-fold affinity improvement due to its bivalent presentation of hIL-2 (**Figure 2A, right; Table S2**). Whereas binding to hIL-2Rβ was completely abolished for hIL-2/F5111 complex (**Figure 2A, right; Table S2**), F5111 IC LN15 bound to hIL-2Rβ with an affinity similar to that of free hIL-2 (**Figure S1C, right**), indicating deficient intramolecular assembly and/or presence of oligomeric IC for this construct. Lengthening the linker between the cytokine and antibody eliminated these issues, as F5111 IC LN25 and F5111 IC LN35 did not engage hIL-2Rβ (**Figures 2A, right, S1C, right, and S1D, right; Table S2**). Consistent with expression observations, F5111 IC LN25 P1 and F5111 IC LN25 P2 showed a small amount of residual binding to hIL-2Rβ, whereas F5111 LN25 P3 was found to ablate receptor interaction (**Figure S1E, right**). Overall, hIL-2Rβ binding studies confirmed the proper assembly and function of F5111 ICs, and support their molecular bias towards T_Reg_ engagement.

### F5111 ICs demonstrate T_Reg_ bias *in vitro*

F5111 ICs were assessed for induction of IL-2-dependent STAT5 phosphorylation on YT-1 human NK cells that either express or lack IL-2Rα, as a surrogate for stimulation of T_Reg_-like versus Eff-like cells, respectively (Kuziel et al., 1993; Yodoi et al., 1985). ICs were compared to hIL-2/F5111 complex at a 1:1 molar ratio, and it was observed that increasing the cytokine to antibody ratio did not affect signaling potency (**Figure S1B; Table S3**).

On IL-2Rα^+^ YT-1 (T_Reg_-like) cells, F5111 IC LN15, F5111 IC LN25, and F5111 IC LN35 induced robust activation, similar to that elicited by free hIL-2 and hIL-2/F5111 complex (**Figures 2B, left and S1F; Table S3**). Increased potency was observed for Control IC compared to free hIL-2 due to its bivalency (**Figure 2B, left; Table S3**), whereas potency of F5111 ICs was slightly reduced compared to that of free IL-2 due to incomplete dissociation of the antibody from IL-2. On IL-2Rα^-^ YT-1 (Eff-like) cells, F5111 IC LN15 had 15-fold weaker potency compared to free hIL-2 and Control IC, and F5111 IC LN25 and F5111 IC LN35 induced little to no activation (**Figures 2B, right and S1G; Table S3**), illustrating the significant IL-2Rα^+^ cell bias for F5111 ICs. hIL-2/F5111 complex was approximately 2-fold less potent than free hIL-2 on IL-2Rα^-^ cells (**Figure 2B, right; Table S3**), highlighting the advantage of the IL-2/F5111 single-chain fusion versus the complex with respect to IL-2Rα^+^ cell bias. The two peaks of F5111 IC LN15 elicited approximately 10-fold weaker activation of IL-2Rα^-^ cells relative to free hIL-2 (**Figure S1H, top; Table S3**). For F5111 IC LN25, P1 and P2 had half maximal effective concentration (EC_50_) values that were 50 and 100-fold weaker than that of hIL-2, respectively, whereas the monomeric P3 had activity that was at least 2,000-fold weaker (**Figure S1H, bottom; Table S3**). Additionally, all three peaks of F5111 IC LN25 showed reduced activation of IL-2Rα^-^ cells compared to F5111 IC LN15, further confirming that increasing the linker length improved IC function. The potencies of F5111 IC LN25 and of F5111 IC LN35 on IL-2Rα^+^ and IL-2Rα^-^ cells were similar; however there was less overall aggregation observed for F5111 IC LN35 (**Figure 1B**). We therefore decided to move forward with F5111 IC LN35, which is hereafter denoted F5111 IC.

To confirm T_Reg_ bias in a mixed cell population, IC activity was interrogated on peripheral blood mononuclear cells (PBMCs) isolated from human whole blood. We found that F5111 IC exhibited reduced potency compared to free hIL-2 and the IL-2/F5111 complex (∼500-fold) as well as Control IC (∼1,000-fold) on T_Reg_s due to incomplete dissociation of the antibody from IL-2 (**Figure 2C, left; Table S4**). However, an equivalent maximum response (E_Max_) value was achieved for all constructs. F5111 IC induced little to no activation on CD8^+^ T cells, as indicated by reduced potency and ∼80% reduction in E_Max_ value (**Figure 2C, middle; Table S4**) compared to free hIL-2 and Control IC. In contrast, hIL-2/F5111 complex actually showed improved potency on CD8^+^ T cells with a milder 35% reduction in E_Max_ relative to free IL-2 and Control IC. Similar to what was observed on CD8^+^ T cells, F5111 IC orchestrated little to no activation of conventional CD4^+^ T cells (T_Conv_), and 55% reduction in E_Max_ compared to free hIL-2 and Control IC (**Figure 2C, right; Table S4**). In contrast, hIL-2/F5111 complex activated T_Conv_ cells with similar potency and E_Max_ as free IL-2, presumably due to complex dissociation. Control IC was slightly more potent than free hIL-2 due to its bivalency. These results indicated that hIL-2/F5111 had little to no T_Reg_ bias as compared to free hIL-2. Taken together, our *ex vivo* studies with human PBMCs revealed that F5111 IC is strongly biased towards activation of T_Reg_s compared to CD8^+^ T and T_Conv_ cells due to its preferential engagement of IL-2Rα^High^ cell subsets.

To better understand the mechanistic basis for the more dramatic potency reduction for F5111 IC as compared to free hIL-2 on human T_Reg_s relative to what was observed for IL-2Rα^+^ YT-1 cells, we measured the IL-2Rα expression levels on primary human cell populations and YT-1 cell lines. We found that the IL-2Rα mean fluorescence intensity (MFI) on T_Reg_s was only 40-fold greater than the fluorescence minus one (FMO) control, whereas IL-2Rα MFI was 75-fold greater than FMO control for IL-2Rα^+^ YT-1 cells (**Figure S2B, left**), indicating that the IL-2Rα^+^ YT-1 express higher levels of IL-2Rα than do T_Reg_s. IL-2Rα MFI levels on CD8^+^ T, T_Conv_, and IL-2Rα^-^ YT-1 cells were similar to those of FMO (**Figure S2B, right**). As F5111 IC binding requires IL-2Rα-dependent disruption of the cytokine/antibody interaction, the lower expression levels of IL-2Rα on human primary cells potentially explain the weakened activity of F5111 IC on primary human T_Reg_s.

### Tuning IC intramolecular affinity modulates IL-2 receptor binding properties and T_Reg_ bias

Based on our understanding of the molecular mechanism for F5111 IC activity, we hypothesized that modulation of the cytokine/antibody affinity would impact relative engagement of IL-2Rα^High^ versus IL-2Rα^Low^ cells, allowing for optimization of IC bias on different cell types with various IL-2Rα expression levels. We conjectured, based on early work comparing complexes containing higher affinity F5111 variants (Trotta et al., 2018), that reducing the affinity of the IL-2/F5111 interaction would lead to enhanced IL-2 signaling, particularly on human T_Reg_s. Informed by the crystallographic structure of the IL-2/F5111 complex (Trotta et al., 2018), we rationally designed a panel of eight single-point alanine mutations of the F5111 antibody, encompassing 3 V_L_ (Y33, Y94, S96) and 5 V_H_ residues (Y35, Y52, Y54, Y60, and V103) residues at the cytokine/antibody interface (**Figure 3A**). Each F5111 variant and the affinity matured F5111.2 antibody (Rondon et al., 2015; Trotta et al., 2018) were formatted and expressed as ICs with 35 amino acid linkers (**Figure S3A**).

**Figure 3.**
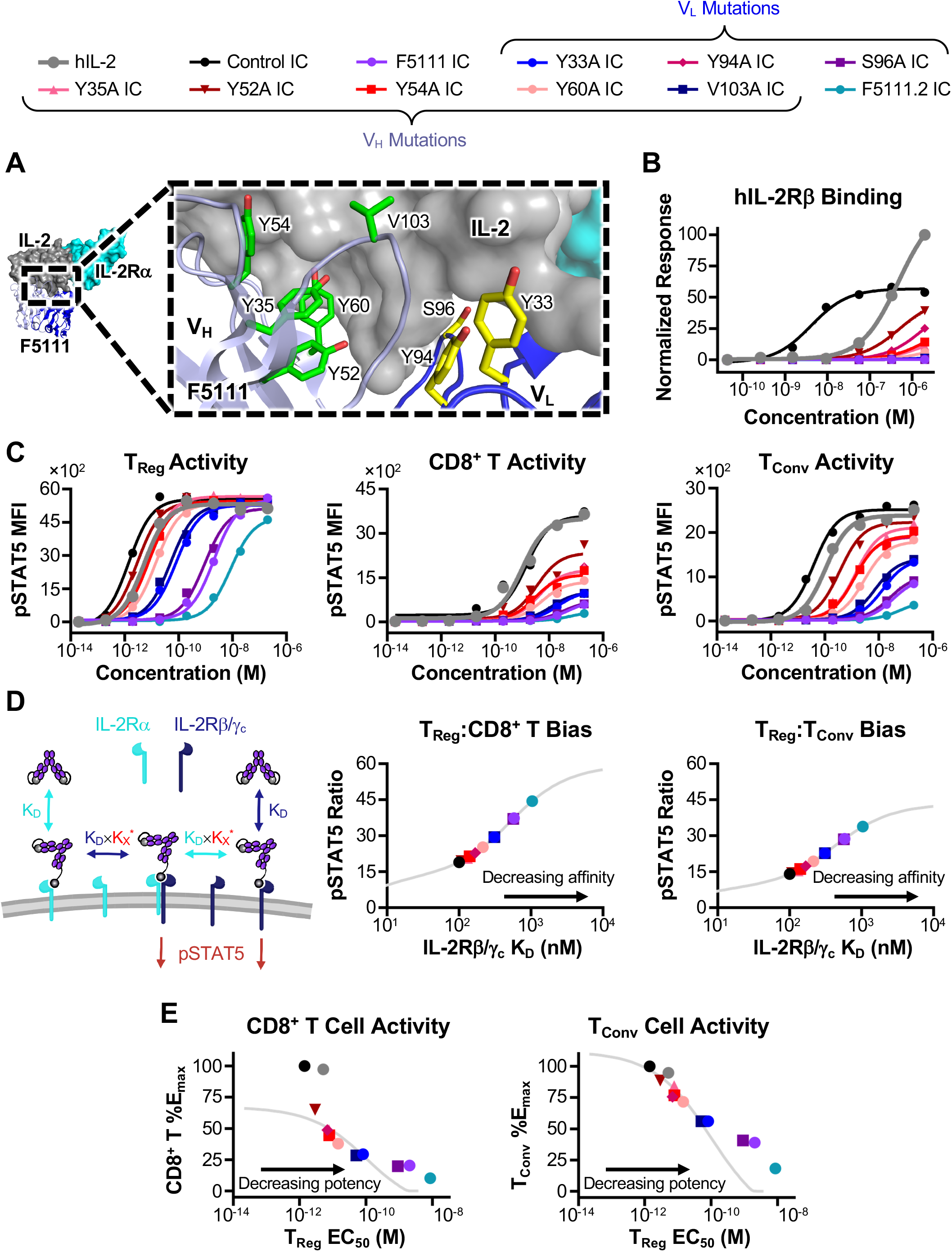
Tuning IC intramolecular affinity modulates IL-2 receptor binding properties and T_Reg_ bias. **(A)** Crystal structure of the hIL-2/F5111 interface (PDB ID 5UTZ) (Trotta et al., 2018) with residues mutated to alanine highlighted in yellow (light chain) or green (heavy chain). Human IL-2Rα is overlaid from the IL-2 cytokine/receptor quaternary complex structure for reference (PDB ID 2B5I) (Wang et al., 2005). **(B)** Equilibrium biolayer interferometry-based titrations of hIL-2, Control IC, and F5111 IC variants binding to immobilized hIL-2Rβ. **(C)** STAT5 phosphorylation response of human T_Reg_ (left), CD8^+^ T (middle), and T_Conv_ (right) cell populations within human PBMCs stimulated with either hIL-2, Control IC, or F5111 IC variants. **(D)** Schematic of the multivalent binding model. Initial association of ICs is assumed to proceed according to the monovalent cytokine/receptor affinity, and subsequent binding events proceed with the monovalent affinity scaled by a K^∗^ parameter (left). Predicted T_Reg_:CD8^+^ T (middle) and T_Reg_:T_Conv_ (right) pSTAT5 ratios for bivalent ICs at 10 pM IC concentration are plotted against predicted IL2Rβ/γ_c_ dissociation constants (K_D_, nM). **(E)** Predicted (lines) and experimental (points) percent of Control IC pSTAT5 E_Max_ for CD8^+^ T (left) or T_Conv_ (right) cells achieved by bivalent ICs is plotted against either the predicted (lines) or experimental (points) pSTAT5 EC_50_ on T_Reg_s. See also **Figure S3; Tables S2 and S4**.

Bio-layer interferometry studies revealed that all F5111 IC variants had minimal engagement of hIL-2, confirming their intramolecular assembly (**Figure S3B; Table S2**). Further, all IC variants bound hIL-2Rα with similar affinities, comparable to that observed for the parent F5111 IC (**Figure S3B; Table S2**). All F5111 IC variants showed significantly attenuated binding to IL-2Rβ compared to free hIL-2 and Control IC, indicating successful receptor blockade (**Figure 3B; Table S2**). However, whereas most of the IC variants completely ablated hIL-2Rβ binding, IC variants Y94A, Y35A, Y52A, Y54A, and Y60A exhibited observable binding at high concentrations, indicating that a potentially weaker cytokine/antibody interaction for these clones allowed for increased IL-2 engagement with hIL-2Rβ.

We sought to determine whether differences in hIL-2Rβ engagement by the IC variants would impact IL-2 signaling bias. In human PBMC studies, all F5111 IC variants were found to either maintain or enhance T_Reg_ potency compared to the parent F5111 antibody (**Figure 3C, left; Table S4**), indicating that the mutations had weakened cytokine/antibody binding, as intended. IC variants were grouped into 3 cohorts: the “high” potency group (Y94A, Y35A, Y52A, Y54A, and Y60A ICs), which had potencies similar to that of Control IC; the “intermediate” potency group (Y33A and V103A ICs), which had potencies ∼50-fold weaker than Control IC; and the “low” potency group (parent F5111 IC and the S96A IC), which had potencies ∼1,000-fold weaker than Control IC. F5111.2 IC impaired the T_Reg_ potency (>6,000-fold weaker than Control IC), confirming that tighter cytokine/affinity interaction led to reduced cytokine/antibody dissociation. As anticipated, weakened cytokine/antibody affinity also potentiated the activity of F5111 IC variants on both CD8^+^ T cells (**Figure 3C, middle; Table S4**) and T_Conv_ cells (**Figure 3C, right; Table S4**). However, all IC variants did show impaired immune effector cell activation as compared to free hIL-2 and Control IC; indicating that they still retained some level of T_Reg_ bias. As expected, IC variants with the highest T_Reg_ potencies led to the most potent activation of CD8^+^ T and T_Conv_ cells, and F5111.2 IC, which had the weakest potency on T_Reg_s, elicited the weakest immune effector cell activation. Moreover, signaling potency on both T_Reg_ and immune effector cells was directly correlated with extent of hIL-2Rβ engagement, as anticipated.

The panel of F5111 IC variants represents a collection of IL-2Rα “probes,” as illustrated by analysis of IL-2Rα MFI within the activated cell populations for various immune cell subsets. At saturating concentrations, the mean IL-2Rα MFIs of activated T_Reg_s were similar following treatment with all F5111 ICs except for the significantly less potent F5111.2 IC (**Figure S3C, left**). This finding suggests that T_Reg_ cell IL-2Rα expression is sufficient to induce cytokine/antibody dissociation for all F5111 ICs other than F5111.2 IC, which requires higher levels of IL-2Rα to induce cytokine/antibody dissociation. On CD8^+^ T and T_Conv_ cells, ICs that induced more potent immune effector cell activation required lower levels of IL-2Rα to stimulate IL-2 signaling (**Figure S3C, middle and right**).

To further explore the relationship between cytokine/antibody affinity and T_Reg_ bias of ICs we developed a multivalent binding model that predicts F5111 IC signaling properties in specific immune cell subsets based on receptor affinity (**Figure 3D, left**). This model was able to accurately predict activities of the IC variants (**Figures S3D-S3F**) and infer hIL-2Rβ/γ_c_ affinities (**Figures S3G and S3H**), something we were unable to do for the experimental data set due to the IC variants not reaching binding saturation. We plotted the predicted affinity of each IC against the predicted phosphorylated STAT5 (pSTAT5) MFI ratios of T_Reg_:CD8^+^ T and T_Reg_:T_Conv_ at a fixed concentration of 0.01 nM (**Figure 3D, middle and right**). As the affinity of the IC towards hIL-2Rβ/γ_c_ decreased, we observed a corresponding increase in T_Reg_ bias. This increased bias resulted from attenuated signaling on both CD8^+^ T and T_Conv_ cells but was also accompanied by a decrease in activation of T_Reg_s (**Figures 3C and 3E**). Thus, a tradeoff was observed between T_Reg_ activation and selectivity. Based on our experimental and computational insights, we selected 3 IC variants for further analysis that improved T_Reg_ potency while minimizing immune effector cell activation (Y60A, Y33A, and V103A ICs).

To improve the therapeutic potential of our engineered ICs, we introduced the N297A mutation into the fragment crystallizable (Fc) region of all cytokine/antibody fusion proteins. The N297A mutation prevents glycosylation at this site and has been reported to significantly impair Fc γ receptor binding and thus reduce antibody effector functions (Delidakis et al., 2022; Mimura et al., 2000, 2001; Saunders, 2019; Tao et al., 1993; Wang et al., 2018). Bio-layer interferometry studies showed no measurable differences between F5111 IC with and without the N297A mutation in terms of binding to hIL-2, hIL-2Rα, and hIL-2Rβ (**Figure S4A; Table S2**). Human PBMC signaling assays demonstrated that the N297A mutation did not affect the activity of Control IC, F5111 IC, or IC variants (Y60A, V103A, Y33A, and F5111.2 ICs) on T_Reg_, CD8^+^ T, and T_Conv_ cells (**Figures S4B and S4C; Table S4**). Thus, all future references to ICs will denote the IC with the N297A mutation unless otherwise noted.

### Parent F5111 IC maximizes T_Reg_ expansion bias *in vivo*

We sought to determine whether the *in vitro* IL-2 signaling bias of our engineered ICs towards activation of T_Reg_s would translate into selective expansion of T_Reg_s in mice. To compare potencies of our ICs between human and mouse primary cells, we evaluated their behavior on splenocytes isolated from non-obese diabetic (NOD) mice. Overall, IC activation on mouse primary cells showed similar trends to that observed on human primary cells (**Figures S4C and S4D; Tables S4 and S5**), albeit with weaker potency, presumably due to the weaker affinity of hIL-2 for mIL-2 receptors as compared to hIL-2 receptors (Spangler et al., 2015b).

To inform *in vivo* dosing strategies, we plotted *in vitro* T_Reg_:CD8^+^ T and T_Reg_:T_Conv_ pSTAT5 MFI ratios as a function of IC concentration (**Figure S4E**). Each IC variant exhibited a unique optimum concentration and maximum level of T_Reg_ bias, and IC variants that were more potent on T_Reg_s were found to have a lower optimum concentration and a higher magnitude bias.

NOD mice were administered 4 daily doses (1.5 μg hIL-2 equivalent) of each IC and immune cell subset expansion within harvested splenocytes was evaluated 24 hours after the last dose (**Figure S5A**). The Y33A IC, the V103A IC, and the Y60A IC showed significant T_Reg_ bias, as evidenced by the T_Reg_:CD8^+^ T, T_Reg_:T_Conv_, and T_Reg_:NK cell ratios (**Figure 4A**). Remarkably, the parent F5111 IC led to the most biased T_Reg_ expansion amongst the ICs tested. The Control IC and F5111.2 IC exhibited the least bias. These *in vivo* studies revealed that the optimum hIL-2/antibody affinity for T_Reg_ bias corresponded to that of the parent F5111 IC. This is visualized by plotting the T_Reg_:CD8^+^ T cell ratio against the T_Reg_ EC_50_ value (obtained from human PBMC STAT5 signaling, **Figure S4C, left; Table S4**) for each IC (**Figure 4B**). If T_Reg_ activity is too weak (as for F5111.2 IC), no activity is observed on any immune cell subset. On the other hand, if the T_Reg_ activity is too potent, all immune cell subsets are activated, confounding T_Reg_ bias (as for Control IC). The optimum T_Reg_ EC_50_ lies between these extrema, and F5111 IC appears to be close to this optimum. Non-negative matrix factorization was performed on this data using a 2-component analysis, and we found that F5111 IC had equivalent T_Reg_ specificity to the other IC variants while maintaining a clear inhibition of immune effector cell activation (**Figure 4C**), thus optimizing the balance between T_Reg_ activity and selectivity.

**Figure 4.**
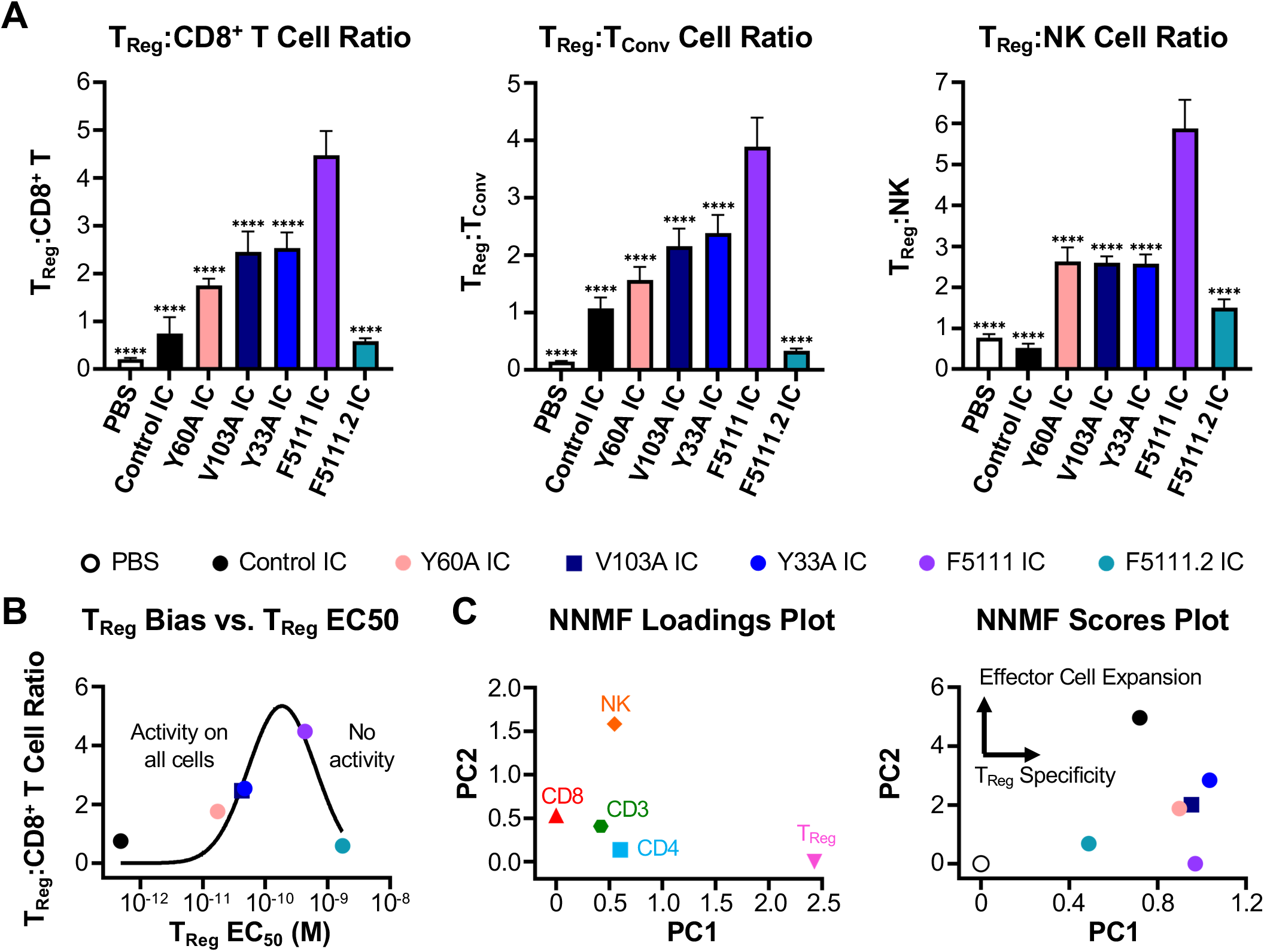
Parent F5111 IC exhibits maximum T_Reg_ expansion bias *in vivo*. **(A)** Ratios of T_Reg_ to CD8^+^ T cells (left), T_Reg_ to T_Conv_ cells (middle), and T_Reg_ to NK cells (right) in spleens harvested from NOD mice treated for four days (*i.p.*) with either PBS (n=4) or 8.2 μg (1.5 μg IL-2 equivalence) Control IC (n=4), Y60A IC (n=5), V103A IC (n=4), Y33A IC (n=4), F5111 IC (n=5), or F5111.2 IC (n=4). Data represent mean + SD. Statistical significance was determined by one-way ANOVA with a Tukey post hoc test. Only statistical significance compared to F5111 IC is shown on the plots (all statistical data are provided in **Table S6**). *P≤0.05, **P≤0.01, ***P≤0.001, ****P≤0.0001. **(B)** Plot showing the average ratio of T_Reg_ to CD8^+^ T cell expansion *in vivo* versus the EC_50_ of STAT5 phosphorylation on T_Reg_ cell populations within stimulated human PBMCs (**Figure S4C, left; Table S4**). Data is fit with a Gaussian distribution. **(C)** Loadings plot of non-negative matrix factorization (NNMF) of immune cell subset expansion studies in NOD mice (left) and scores plot of NNMF of immune cell subset expansion studies in NOD mice (right). Using two components, 84% of the variance of the PBS-normalized *in vivo* expansion data was explained. See also Figures S4 and S5.

We also tested immune cell subset expansion elicited by our panel of IC variants in a humanized mouse model. BALB/c Rag2^-/-^ γ_c_^-/-^ H2^d^ (BRG) mice engrafted with human PBMC were treated with a single dose of the various ICs. Immune cell expansion in the peritoneum was evaluated 4 days later, and in contrast to what we observed in NOD mice, in the humanized model we saw that the greatest T_Reg_ expansion bias was observed for the Y60A and Y33A IC variants (**Figure S4F**), which have weaker cytokine/antibody affinity than the parent F5111 IC. Thus, the optimal T_Reg_-promoting therapy may vary between mouse and human models.

To determine whether Fc effector function would benefit or hinder F5111 IC-induced T_Reg_ expansion, we compared immune activation by the IC with and without the N297A mutation. We found that F5111 IC with the N297A mutation (impaired Fc effector function) led to significantly more T_Reg_ bias and >3-fold more T_Reg_ expansion (**Figure S5B**). Based on our collective *in vivo* T_Reg_ expansion studies, we elected to employ F5111 IC (with impaired Fc effector function) in mouse models of disease.

### F5111 IC demonstrates greater T_Reg_ bias than hIL-2/F5111.2 complex

We hypothesized that the IC could have stability improvements over cytokine/antibody complexes; thus, we compared the previously reported lead IL-2/antibody complex (hIL-2/F5111.2 complex) (Trotta et al., 2018) to our lead construct (F5111 IC). The N297A mutation was installed in the F5111.2 antibody for consistency with F5111 IC. Bio-layer interferometry studies showed that the F5111.2 antibody and hIL-2/F5111.2 complex bound to hIL-2 (at a 1:1 molar ratio), whereas Control IC had no binding to hIL-2, and the F5111 IC had minimal binding above background (**Figure S5C, left; Table S2**). hIL-2/F5111.2 complex bound hIL-2Rα with ∼18-fold weaker affinity than F5111 IC and Control IC and ∼3-fold weaker affinity than free hIL-2 (**Figure S5C, middle; Table S2**). This is in agreement with previous reports that hIL-2/F5111.2 complex had weaker affinity to hIL-2Rα (Trotta et al., 2018) compared to free hIL-2. Both hIL-2/F5111.2 complex and F5111 IC fully blocked binding to hIL-2Rβ (**Figure S5C, right; Table S2**).

We next compared T_Reg_ bias induced by either F5111 IC or hIL-2/F5111.2 complex in human PBMCs. On T_Reg_s, F5111 IC was ∼1,700-fold weaker than Control IC, whereas hIL-2/F5111.2 complex was ∼10-fold weaker than Control IC (**Figure 5A, left; Table S4**). On CD8^+^ T cells, F5111 IC was ∼6-fold less potent than Control IC, whereas hIL-2/F5111.2 complex was in fact 15-fold more potent than Control IC (**Figure 5A, middle; Table S4**). F5111 IC and hIL-2/F5111.2 reduced the E_Max_ by 85% and 80%, respectively, relative to Control IC. Similar trends were observed on T_Conv_ cells; F5111 IC was ∼1,000-fold weaker with a 60% reduction in E_Max_ compared to Control IC, whereas hIL-2/F5111.2 complex was ∼7-fold weaker and reduced E_Max_ by 50% (**Figure 5A, middle; Table S4**).

**Figure 5.**
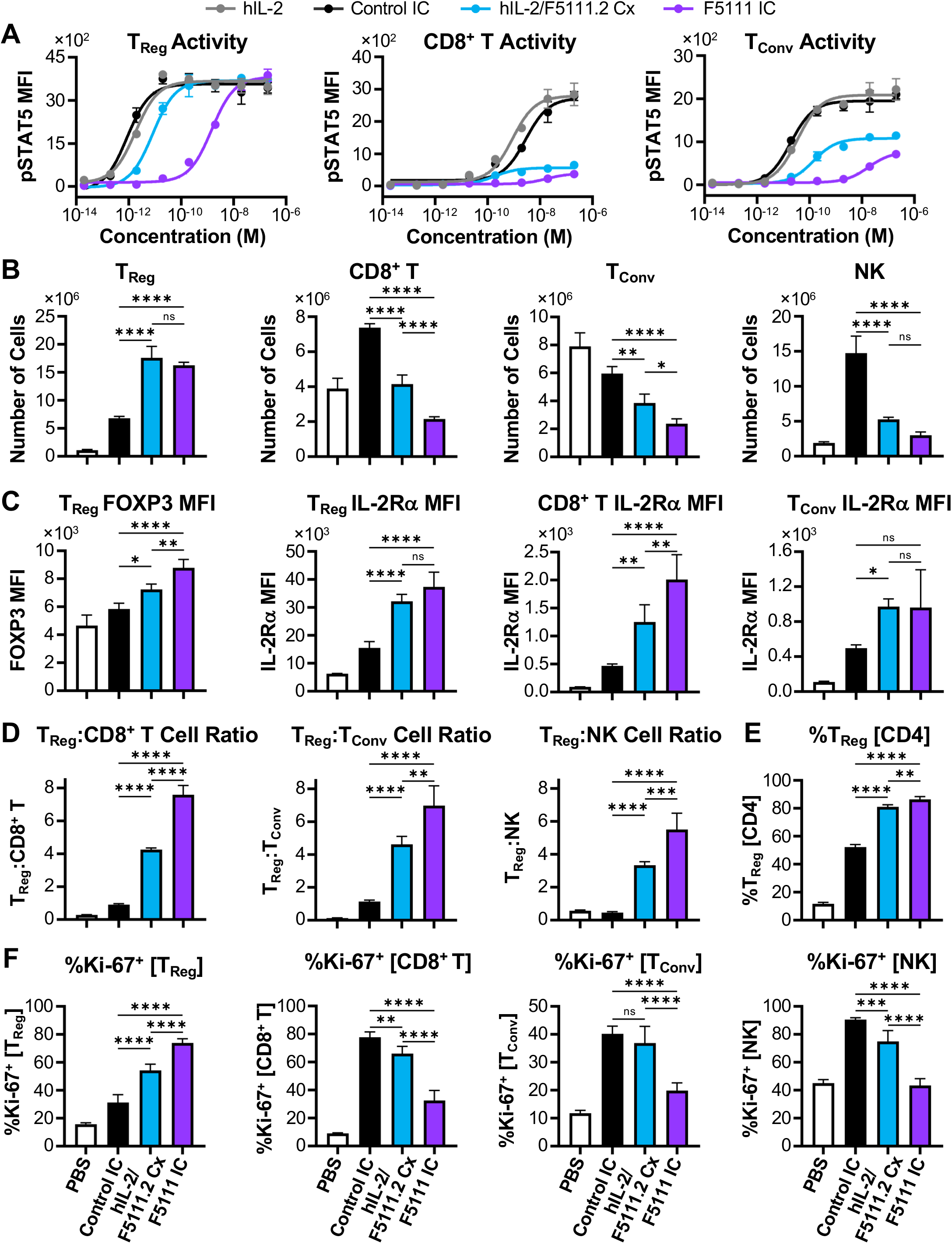
F5111 IC demonstrates greater T_Reg_ bias than hIL-2/F5111.2 complex. **(A)** STAT5 phosphorylation response of human T_Reg_ (left), CD8^+^ T (middle), and T_Conv_ (right) cell populations within human PBMCs stimulated with either hIL-2, Control IC, hIL-2/F5111.2 complex (Cx, 1:1 molar ratio), or F5111 IC (n=3). **(B-E)** NOD mice (n=4 per group) were treated daily for 4 days (*i.p.*) with either PBS, 1.5 μg hIL-2 complexed with 6.6 μg F5111.2 antibody (1:2 molar ratio, hIL-2/F5111.2 Cx), or 8.2 μg (1.5 μg IL-2 equivalence) Control IC or F5111 IC. 24 hours after the last dose, spleens were harvested for analysis. **(B)** Number of T_Reg_, CD8^+^ T, T_Conv_, and NK cells. **(C)** MFI of FOXP3 within T_Reg_s (left) and IL-2Rα MFI within T_Reg_, CD8^+^ T, and T_Conv_ populations. **(D)** Ratio of T_Reg_ to CD8^+^ T cells (left), T_Reg_ to T_Conv_ cells (middle), and T_Reg_ to NK cells (right). **(E)** Percent T_Reg_s within the CD4^+^ T cell population. **(F)** Percent Ki-67^+^ cells within T_Reg_, CD8^+^ T, T_Conv_, and NK cell populations from spleen harvested from NOD mice (n=5 per group) treated daily for 4 days (*i.p.*) with either PBS or 1.5 μg hIL-2 equivalence (as described above) and analyzed 24 hours after the final dose. Data represent mean ± SD for all panels. Statistical significance was determined by one-way ANOVA with a Tukey post hoc test. Significance is shown between Control IC, hIL-2/F5111.2 Cx, and F5111 IC. All statistical data are provided in **Table S6**. *P≤0.05, **P≤0.01, ***P≤0.001, ****P≤0.0001. See also **Figure S5; Table S4**.

We explored whether superior T_Reg_ bias in human PBMCs would correspond to enhanced *in vivo* T_Reg_ bias for F5111 IC relative to hIL-2/F5111.2 complex. To this end, we compared immune cell subset expansion in NOD mice following 4 daily injections of phosphate-buffered saline (PBS), hIL-2/F5111.2 complex (1.5 μg IL-2 equivalent; 2:1 molar ratio of cytokine to antibody), or Control IC or F5111 IC (1.5 μg IL-2 equivalent). Both F5111 IC and hIL-2/F5111.2 complex led to 15-fold increases in the total number of T_Reg_s compared to untreated mice, and induced 3-fold more T_Reg_ expansion than Control IC (**Figure 5B**). Additionally, F5111 IC and hIL-2/F111.2 complex treatment led to significantly less CD8^+^ T, T_Conv_, and NK cell expansion compared to Control IC, and F5111 IC-treated mice had fewer CD8^+^ T and T_Conv_ cells than hIL-2/F111.2 complex-treated animals (**Figure 5B**). Both F5111 IC and hIL-2/F5111.2 complex induced elevated expression of forkhead box P3 (FOXP3) and IL-2Rα within the T_Reg_ population, as compared to Control IC, and F5111 IC induced significantly more FOXP3 expression relative to the complex (**Figure 5C**). Additionally, F5111 IC and hIL-2/F5111.2 complex induced elevated IL-2Rα expression on CD8^+^ T and T_Conv_ cells as compared to Control IC (**Figure 5C**). Compared to Control IC, we found that both F5111 IC and hIL-2/F5111.2 complex increased the T_Reg_:CD8^+^ T, T_Reg_:T_Conv_ and T_Reg_:NK cell ratios, and F5111 IC induced significantly more T_Reg_ expansion bias than did the complex (**Figure 5D**). Notably, F5111 IC treatment resulted in >80% T_Reg_s within the CD4^+^ population, a massive expansion over the 10% T_Reg_ proportion observed in saline-treated mice (**Figure 5E**). We further found that F5111 IC treatment resulted in a significantly higher percentage of Ki-67^+^ cells within T_Reg_s and significantly lower percentage within CD8^+^ T, T_Conv_, and NK as compared to hIL-2/F5111.2 complex and Control IC **(Figure 5F**), indicating that F5111 IC induced more actively proliferating T_Reg_s. Taken together, our *in vivo* results showcase the T_Reg_-biasing capacity of F5111 IC, and demonstrate its superiority to cytokine/antibody complex.

### F5111 IC induces durable expansion of functional T_Reg_s

To further characterize T_Reg_ expansion induced by F5111 IC, we examined the dose dependence of this phenomenon. We found that noticeable decreases in the percent of T_Reg_s within the CD4^+^ T cell population were not observed until the dose was decreased by two-thirds (to 0.5 μg IL-2 equivalence) (**Figure S5D**). To characterize the durability of F5111 IC-induced T_Reg_ expansion bias, we analyzed mouse spleens harvested 1, 3, 5, and 7 days after 4 daily F5111 IC injections. The largest percentage of T_Reg_s within the CD4^+^ population (72%) was observed 1 day after the last dose, and the percentage declined monotonically for each subsequent timepoint but still remained elevated 1 week after the final dose (**Figures 6A and 6B**). Similarly, maximum T_Reg_ bias was observed 1 day after the last dose, and the bias was mostly gone after 5 days (**Figure 6C**). Characterization of F5111 IC serum half-life in C57BL/6 mice revealed that the IC follows a two-phase decay with a fast half-life of ∼5 minutes and a slow half-life of 35 hours (**Figure 6D**).

**Figure 6.**
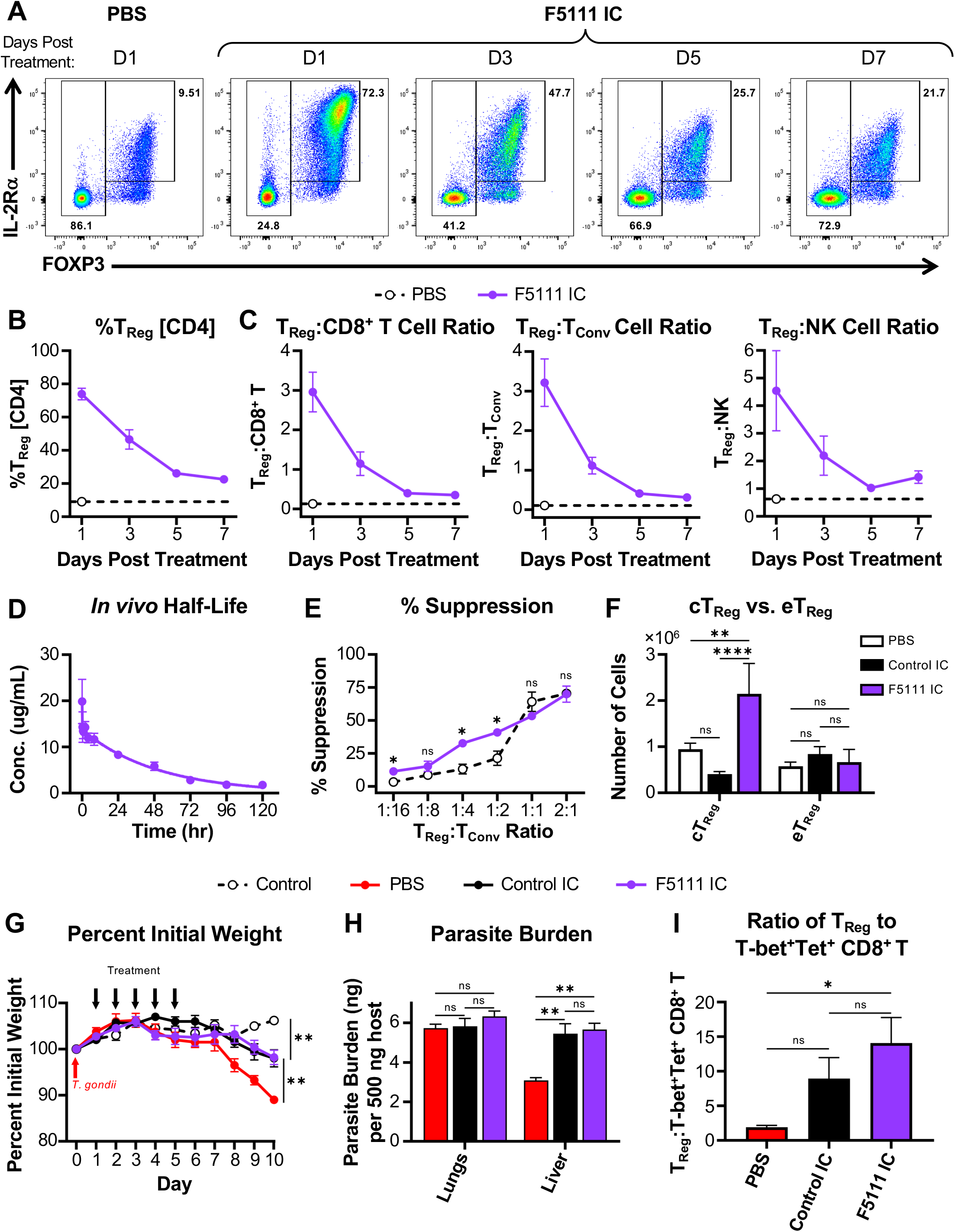
F5111 IC induces durable expansion of functional T_Reg_s without compromising infectious disease clearance. **(A-C)** C57BL/6 mice (n=3 per group) were treated daily for 4 days (*i.p.*) with either PBS or 8.2 μg F5111 IC (1.5 μg IL-2 equivalence) and spleens were harvested 1, 3, 5, and 7 days after the last dose. **(A)** Representative flow plots illustrating percent of T_Reg_s and levels of IL-2Rα and FOXP3 expression within CD4^+^ T cells at the indicated time points. **(B)** Percent T_Reg_s within the CD4^+^ T cell population at the indicated time points. **(C)** Ratios of T_Reg_ to CD8^+^ T cells (left), T_Reg_ to T_Conv_ cells (middle), and T_Reg_ to NK cells (right) at the indicated time points. Dashed lines represent the baseline values for the PBS-treated cohort harvested 1 day after the last dose. Data represent mean ± SD. **(D)** Serum half-life of F5111 IC in C57BL/6 mice (n=5) treated retro-orbitally with 2 mg/kg F5111 IC (∼0.4 mg/kg IL-2 equivalence). Data represent mean ± SD. **(E)** C57BL/6 CD45.1 RFP-FOXP3 mice were treated for 4 days with either PBS (n=3) or 6.2 μg F5111 IC (1.125 μg IL-2 equivalence, n=2). Plot represents the percent suppression of T_Conv_ cell proliferation with titrating ratios of T_Reg_s (isolated and pooled from spleens of treated mice) to T_Conv_ cells (isolated and pooled from untreated C57BL/6 CD45.2 mice, n=5). Data represent mean ± SD of technical replicates. Statistical significance at each ratio was determined by two-tailed unpaired Student *t* test. **(F)** C57BL/6 mice (n=5 per group) were dosed (*i.p.*) on days 0, 2, and 4 with either PBS or 8.2 μg (1.5 μg IL-2 equivalence) Control IC or F5111 IC, and spleens were harvested on day 6. Total number of conventional T_Reg_s (cT_Reg_s, CD4^+^FOXP3^+^IL-2Rα^High^BCL-2^High^) and effector T_Reg_s (eT_Reg_s, CD4^+^FOXP3^+^IL-2Rα^Low^BCL-2^Low^) are presented. Statistical significance was determined separately for cT_Reg_s and eT_Reg_s by one-way ANOVA with a Tukey post hoc test. Data represent mean + SD. **(G-I)** C57BL/6 mice were administered (*i.p.*) 25 cysts of the ME49 strain of *Toxoplasma gondii* (*T. gondii*) on day 0. Control group designates disease-free mice that were not given cysts. Starting on day 1, mice were treated daily for 5 days (*i.p.*) with either PBS (Control, n=5; PBS, n=4) or 8.2 μg (1.5 μg IL-2 equivalence) Control IC (n=5) or F5111 IC (n=5). Mice were sacrificed on day 10. **(G)** Mouse weight throughout the study. Data represent mean ± SEM. **(H)** Parasite burden evaluated in the lungs and liver. **(I)** Ratio of T_Reg_s to T-bet^+^Tetramer (Tet)^+^ CD8^+^ T cells within the spleen. Data represent mean + SEM for **(H)** and **(I)**. Statistical significance was determined by one-way ANOVA with a Tukey post hoc test. Significance in percent initial weight is shown on day 10 compared to F5111 IC. All statistical data are provided in **Table S6**. *P≤0.05, **P≤0.01, ***P≤0.001, ****P≤0.0001. See also **Figure S5**.

To characterize the functional properties of T_Reg_s expanded by F5111 IC, we conducted a suppression assay using T_Reg_s isolated from spleens of C57BL/6 CD45.1 RFP-FOXP3 mice treated with either PBS or F5111 IC. Spleens from C57BL/6 CD45.2 mice were taken for isolation of naïve T_Conv_ cells (CD45.2^+^CD4^+^CD8^-^IL-2Rα^-^CD62L^High^CD44^Low^). Compared to T_Reg_s isolated from saline-treated mice, we found that the T_Reg_s expanded by F5111 IC had equivalent suppressive capacity against naïve T_Conv_ cells isolated from C57BL/6 CD45.2 mice (**Figure 6E**). We also assessed the phenotype of F5111 IC-expanded T_Reg_s from C57BL/6 mice and found significantly increased numbers of conventional T_Reg_s (cT_Reg_s, CD4^+^FOXP3^+^IL-2Rα^High^BCL-2^High^) but not effector T_Reg_s (eT_Reg_s, CD4^+^FOXP3^+^IL-2Rα^Low^BCL-2^Low^) (**Figure 6F**). Interestingly, Control IC treatment resulted in a reduced number of cT_Reg_s as compared to saline-treated mice, whereas eT_Reg_ numbers were not affected. This aligns with the mechanism of F5111 IC, which biases the molecule towards immune cell subsets with high levels of IL-2Rα.

### Treatment with F5111 IC does not impair immune response to infection

A potential concern for therapeutic administration of a T_Reg_-promoting agent such as F5111 IC is that it may interfere with T cell-mediated clearance of infection (Belkaid and Tarbell, 2009; Belkaid et al., 2006; Oldenhove et al., 2009; Sacks and Anderson, 2004). We therefore interrogated how F5111 IC would act in a mouse model of toxoplasmosis. We found that treatment with either F5111 IC or Control IC following *Toxoplasma gondii* infection conferred protection against disease-associated weight loss compared to saline treatment (**Figure 6G**). Additionally, no significant differences were observed in parasite burden in the lung upon F5111 IC treatment (**Figure 6H**). Control IC treatment increased parasite burden in the liver relative to saline treatment, and F5111 IC treatment increased burden to the same extent. Histological comparisons of immune-mediated liver pathology in Control IC-treated mice demonstrated exacerbated leukophilia and necrosis when compared to PBS-treated mice (**Figure S5E**). Interestingly, despite similar parasite burden to Control IC-treated animals, mice treated with F5111 IC demonstrated an overall reduction in immunopathological changes compared to saline-treated mice (**Figure S5E)**. Importantly, F5111 IC had the same effects on mouse weight and parasite burden as Control IC despite inducing a significant increase in T_Reg_ numbers (**Figure S5F**) and significantly skewing ratios of T_Reg_s to both T-bet^+^Tetramer^+^ CD8^+^ T cells (**Figure 6I**) and T-bet^+^Tetramer^+^ CD4^+^ T_Conv_ cells (**Figure S5G**). Additionally, the expanded T_Reg_s showed evidence of active proliferation (**Figure S5H**) and upregulation of IL-2Rα (**Figure S5I**). This suggests that although F5111 IC promotes T_Reg_ activation and expansion, it does not lead to impaired infection clearance.

### F5111 IC efficacy in a mouse model of colitis

To evaluate the potential for F5111 IC to prevent autoimmune disease development, we assessed its performance as a prophylactic treatment in a mouse model of dextran sulfate sodium (DSS)-induced colitis (Chassaing et al., 2014; Cooper et al., 1993; Okayasu et al., 1990; Spangler et al., 2015b, 2018) (**Figure 7A**). Both F5111 IC and hIL-2/F5111.2 complex treatment significantly reduced the severity of weight loss as compared to Control IC and saline treatment (**Figures 7B and S6A**). Additionally, F5111 IC and hIL-2/F5111.2 complex treatment led to reduced disease activity scores on Day 15 as compared to the control treatments (**Figures 7C and S6B**). Relative to control mice without DSS exposure, mice treated with F5111 IC and hIL-2/F5111.2 complex exhibited a less significant reduction in colon length as compared to mice treated with Control IC and saline (**Figure 7D**). To further evaluate protection against disease-related intestinal damage we performed histology on colon cross sections. Both F5111 IC and hIL-2/F5111.2 complex treatment led to reduced histopathological scores compared to the control treatments (**Figures 7E and S6C**). Overall, this *in vivo* model of colitis illustrated the capacity of F5111 IC to confer protection against autoimmune disease pathogenesis.

**Figure 7.**
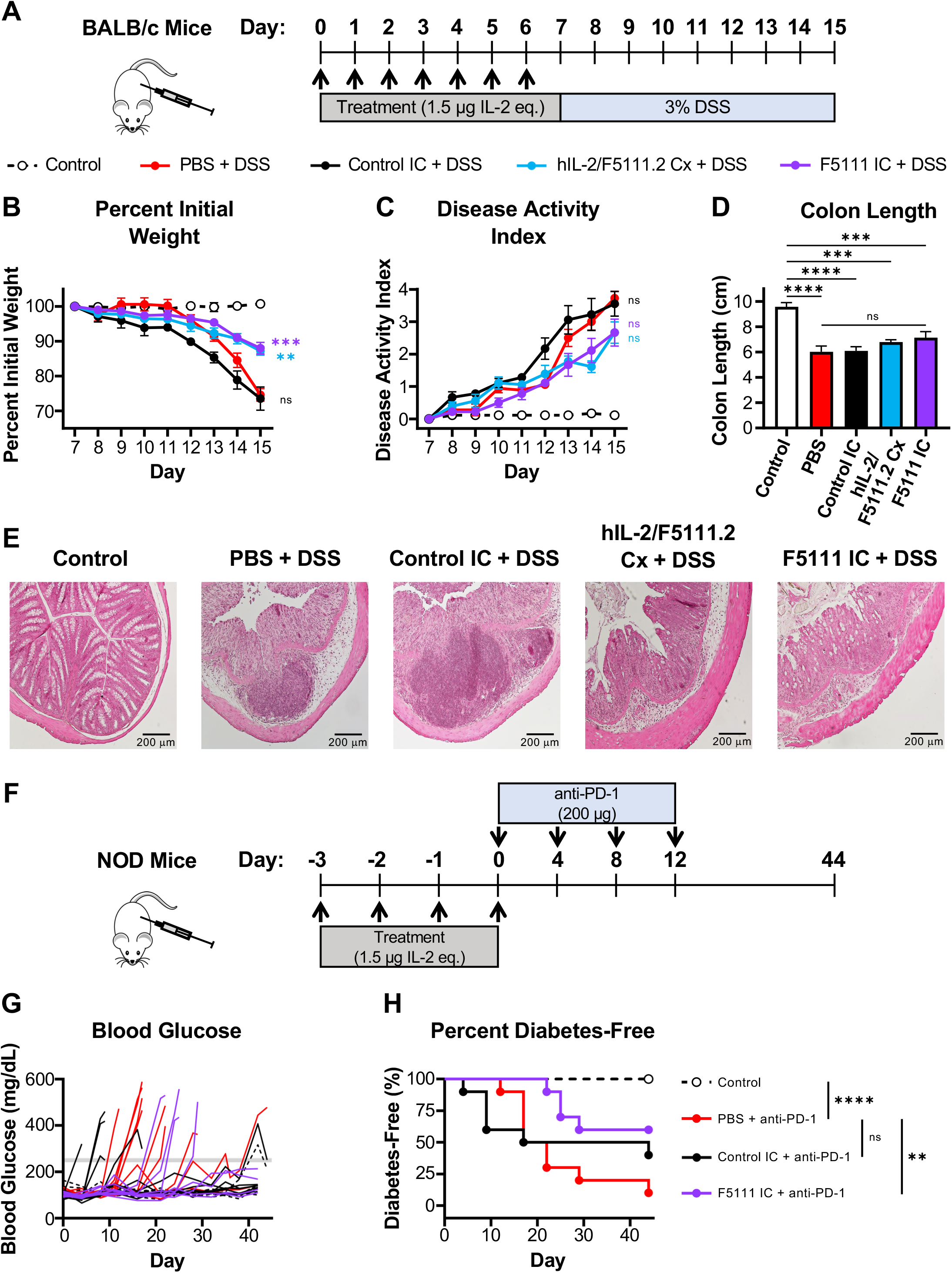
F5111 IC confers protection in mouse models of autoimmune disease. **(A)** BALB/c mice (n=6 per group) were treated daily for 7 days (*i.p.*) with either PBS (Control and PBS), 1.5 μg hIL-2 complexed with 6.6 μg F5111.2 antibody (1:2 molar ratio, hIL-2/F5111.2 Cx), or 8.2 μg (1.5 μg IL-2 equivalence) Control IC or F5111 IC. Beginning on day 7, all groups except for the disease-free cohort (Control) were administered 3% DSS in their drinking water. Mice were sacrificed on day 15. **(B)** Weight change throughout the study. **(C)** Disease activity index (DAI) over the course of the study. **(D)** Colon lengths on day 15 (n=6 PBS, Cx, F5111 IC, Control; n=5 Control IC). **(E)** H&E stained colons harvested from treated mice (n=5 Control, Control IC; n=6 PBS, Cx, F5111 IC). Data represent mean ± SEM. Statistical significance was determined by one-way ANOVA with a Tukey post hoc test. The plots of weight change and DAI show significance of Control IC, Cx, and F5111 IC treated mice versus PBS treated mice on day 15. For colon length, significance is shown compared to Control group and between PBS and F5111 IC treated mice. **(F-H)** 8-week-old NOD mice (n=10 per group) were treated daily for 4 days (*i.p.*, days -3, -2, -1, 0) with either PBS (Control and PBS) or 8.2 μg (1.5 μg IL-2 equivalence) Control IC or F5111 IC. Starting on day 0 (4 hours after the last IC dose), mice were administered anti-PD-1 antibody (200 μg) every 4 days until day 12. Control group designates mice that did not receive anti-PD-1 antibody. **(G)** Blood glucose concentrations over the course of the study. The threshold 250 mg/dL value is indicated by the gray line. **(H)** Percent diabetes-free mice. Statistical significance was determined by pairwise comparisons using the Log-rank (Mantel-Cox) test. Statistical significance compared to mice treated with PBS + anti-PD-1 is shown. All statistical data are provided in **Table S6**. *P≤0.05, **P≤0.01, ***P≤0.001, ****P≤0.0001. See also **Figure S6**.

### F5111 IC is protective in a mouse model of immune checkpoint inhibitor-induced diabetes mellitus

To further evaluate the therapeutic potential for F5111 IC in autoimmune disease treatment, we examined its performance in a murine model of immune checkpoint inhibitor-induced diabetes mellitus (Ansari et al., 2003; Fife et al., 2006; Hu et al., 2020). This mouse model mimics the etiology of a current clinical concern, in which cancer patients treated with immunotherapies are at risk of developing a broad spectrum of immune-related adverse events (irAEs), including diabetes mellitus (Quandt et al., 2021; Stamatouli et al., 2018; Young et al., 2018). NOD mice (8 weeks old) were treated for 4 days with either PBS, Control IC, or F5111 IC followed by 4 daily doses of anti-mouse PD-1 antibody to induce disease (**Figure 7F**). Mice treated with PBS started developing diabetes as early as day 12, whereas a proportion of mice treated with Control IC exhibited accelerated onset of diabetes starting on day 4 (**Figures 7G and 7H**). In contrast, all mice treated with F5111 IC remained diabetes free until day 22. On day 44 (prior to the onset of spontaneous disease in the control group that did not receive anti-PD-1), 60% of mice treated with F5111 IC were diabetes-free, whereas only 10% of mice in the saline-treated group were diabetes-free. Overall, disease-free survival was found to be significantly improved for F5111 IC-versus saline-treated mice that received immune checkpoint inhibitor, whereas a statistically significant improvement was not observed for Control IC-versus saline-treated mice that received immune checkpoint inhibitor. To further assess the durability of F5111 IC therapy in this model, we performed a similar study in which anti-mouse PD-1 antibody treatment was extended through day 16 (**Figure S6D**). The experiment was continued through day 175, demonstrating that F5111 IC led to significant long-term protection compared to saline in immune checkpoint inhibitor-treated mice, as no additional animals developed diabetes after day 31 (**Figures S6E and S6F**). Control mice that did not receive anti-PD-1 treatment began developing spontaneous diabetes beginning on day 50, and by day 157, all of the mice in this cohort had developed disease. Collectively, these studies highlight that the biased T_Reg_ expansion induced by F5111 IC is protective in preventing the onset of immune checkpoint inhibitor-induced diabetes mellitus in mice and that this therapy may also have benefit in preventing spontaneous diabetes onset in NOD mice.

## Discussion

Biased cytokine therapies are of great interest in the therapeutic discovery and development landscapes due to their potential to harness the powerful immunomodulatory properties of these ligands to precisely regulate their biological effects. Ongoing efforts to enable cytokine targeting include direct cytokine engineering (Peterson et al., 2018; Carmenate et al., 2018; Khoryati et al., 2020; Glassman et al., 2021), selective PEG-ylation (Charych et al., 2016; Dixit et al., 2021; Zhang et al., 2021), and complexation of cytokines with anti-cytokine antibodies (Arenas-Ramirez et al., 2016; Boyman and Sprent, 2012; Boyman et al., 2006b, 2006a; Karakus et al., 2020; Lee et al., 2020; Spangler et al., 2015b; Trotta et al., 2018). Here, we build upon previous work that developed an anti-hIL-2 antibody (F5111, later affinity matured to F5111.2) which selectively directs the activity of the cytokine towards IL-2Rα^High^ T_Reg_s (Trotta et al., 2018). To facilitate clinical realization of this technology, we engineered a single-chain fusion protein that links hIL-2 to the F5111 antibody. Compared to hIL-2/antibody complexes, our molecule benefits from extended *in vivo* half-life (due to neonatal Fc receptor-mediated recycling (Finkelman et al., 1993; Roopenian and Akilesh, 2007)), enhanced stability, reduced counterproductive activation of immune effector cells, and a stoichiometrically balanced cytokine to antibody ratio that does not require optimization. We found that F5111 IC induced remarkably potent *in vivo* expansion of T_Reg_s while reducing abundance of CD8^+^ T and T_Conv_ cells, leading to dramatic bias towards immunosuppressive effects that was significantly more pronounced than that of the hIL-2/F5111.2 complex.

It has previously been found that tethering IL-2 to an anti-IL-2 antibody enhances the apparent affinity of the cytokine/antibody interaction (Spangler et al., 2018). However, the mechanism of action for biased immune activation by hIL-2/F5111.2 complex relies on the disruption of cytokine/antibody interactions, specifically in the presence of IL-2Rα (Trotta et al., 2018). We speculated that the strengthened hIL-2/F5111.2 interaction in the context of the IC would be too strong, thereby reducing cytokine dissociation. Thus we employed the weaker affinity F5111 antibody in the single-chain fusion construct. In fact, when the F5111.2 antibody was formatted as an IC, we observed severely impaired activation of all human immune cell subsets as well as impaired T_Reg_ expansion and bias *in vivo*. Thus, although hIL-2/F5111.2 complex led to enhanced T_Reg_ bias compared to the hIL-2/F5111 complex, F5111 outperformed F5111.2 when formatted as ICs. This observation sets a limit to the desired stability for the cytokine/antibody interaction within an intramolecularly bound IC to enable exchange in the presence of IL-2Rα.

Based on this finding, coupled with the differential expression patterns for IL-2Rα between mouse and human immune cells, we designed a panel of F5111 IC variants with varying affinities between the cytokine and antibody and, consequently, varying degrees of IL-2 interaction with IL-2 receptors. Interestingly, although cellular studies indicated that our engineered ICs potentiated T_Reg_ activation, we found a relationship that this improved T_Reg_ potency was accompanied by increased potency on immune effector cell subsets. Although the affinities of all variants for IL-2Rα were identical, the discrepancies in IL-2Rβ affinities for the various ICs presumably drove differential engagement of immune effector cell subsets. Indeed, we found that within activated CD8^+^ T and T_Conv_ cell populations, the IC variants that more potently activated T_Reg_s trended toward lower levels of IL-2Rα expression, further indicating that they experience a lower threshold for exchange in the presence of IL-2Rα. These critical insights from our panel of F5111 IC affinity variants provide a roadmap for designing and optimizing biased therapies for the IL-2 system.

We deployed our panel of F5111 IC variants *in vivo*, and a clear cytokine-antibody affinity optimum for maximal T_Reg_ expansion bias in mice was identified, balancing activation of T_Reg_s with concurrent stimulation of immune effector cell subsets. Amongst all the IC variants, the parent F5111 IC induced the most T_Reg_-biased expansion *in vivo*. We expect this to be because the IC variants that impaired IL-2/antibody affinity led to increased potency on human immune effector cell populations *in vitro,* resulting in a reduced T_Reg_ bias *in vivo* as compared to F5111 IC. We also found that the T_Reg_ EC_50_ for F5111 IC was poised at an optimum: potentiating activity on T_Reg_s (Control IC, Y60A IC, V103A IC, Y33A IC) confounded bias by activating immune effector cells as well, whereas weakening activity on T_Reg_s (F5111.2 IC) resulted in poor stimulation of all cell types. It is important to note that there are differences in IL-2 receptor subunit expression levels between human and mouse immune cells, and the affinities of those subunits towards hIL-2 across species are different (Spangler et al., 2015b). These discrepancies highlight the importance of isolating a panel of IC variants with unique molecular properties, and indeed we observed that the optimal IC for biased T_Reg_ expansion in humanized mice differed from that in immune-competent mice. This further suggests the possibility for developing patient-specific treatments to address the heterogeneity of IL-2 receptor expression levels, carefully optimizing dosing schedules for each construct.

F5111 design efforts emphasized the importance of linker length in optimizing IC function. Our findings suggest that shorter linker lengths hindered intramolecular assembly of the cytokine and antibody, leading to the emergence of higher order oligomeric structures that represent intermolecular interactions between ICs. Moreover, IC variants with higher potency on T_Reg_s eluted as broad single peaks by SEC, whereas lower potency variants had more distinct peaks that eluted earlier, and F5111.2 IC eluted as a single peak at the expected molecular weight for the monomeric IC. This suggests that stronger cytokine/antibody interactions reduce the prevalence of IC oligomerization. Direct *in vivo* comparison of ICs with intact versus impaired effector function capabilities illustrated that curtailing antibody effector function is critical for IC function in order to avoid T_Reg_ depletion via antibody-dependent cellular cytotoxicity (ADCC) (Saunders, 2019; Wang et al., 2018). Collectively, these findings will help guide future design of IC constructs, especially within the growing research area of IL-2/antibody complexes and ICs (Karakus et al., 2020; Lee et al., 2020; Sahin et al., 2020). Furthermore, use of the modular and clinically employed hIgG1 scaffold allows for extension of the single-chain fusion approach to other cytokine systems.

Our *in vivo* therapeutic studies showed that F5111 IC selectivity activates and expand T_Reg_s to improve autoimmune disease outcomes in mouse models, without compromising pathogen clearance. This represents an important translational achievement for the application of human IL-2-based therapies in autoimmune disease treatment, which are hindered by the cytokine’s short half-life, dosing complications, off-target activation, and toxicity (Alva et al., 2016; Dong et al., 2021; Klatzmann and Abbas, 2015). Our findings in an experimental mouse model of immune checkpoint inhibitor-induced diabetes mellitus (Ansari et al., 2003) represent the first deployment of an IL-2-based therapy in this system, and we report that F5111 IC confers long-term protection against drug-induced disease and continues to protect mice from disease even when control mice that did not receive anti-PD-1 develop spontaneous disease. Remarkably, by delaying disease onset, F5111 IC has the reverse effect of Control IC, which accelerates disease development, likely due to stimulation of immune effector cell subsets. This acceleration of diabetes onset resonates with previous reports that IL-2 and untethered IL-2/antibody complexes can exacerbate disease pathogenesis (Dong et al., 2021; Tang et al., 2008; Wesley et al., 2010), highlighting the safety, selectivity, and efficacy advantages for our stable single-chain cytokine/antibody fusion protein. The generality of our T_Reg_-potentiating approach allows for our novel therapeutic to be extended to additional autoimmune conditions including multiple sclerosis, systemic lupus erythematosus, and graft-versus-host disease (Klatzmann and Abbas, 2015; Koreth et al., 2011; Webster et al., 2009). Additionally, selective T_Reg_ expansion mediated by F5111 IC could be leveraged to suppress anti-drug immune responses or to prevent irAEs in patients undergoing cancer treatment (June et al., 2017; Kang et al., 2021; Quandt et al., 2021; Stamatouli et al., 2018). Additionally, F5111 could be incorporated with antigen-specific immune-activating technologies to enable targeted activation of disease-protective T_Reg_s. Overall, this work presents a stable, off-the-shelf IL-2 therapy that potently expands T_Reg_s, introducing a new category of cytokine/antibody fusion proteins that promises to have extensive applications both as a research tool and for therapeutic design.

## Acknowledgments

The authors acknowledge funding from the National Institutes of Health (K99CA246061 to A.Y., U01AI148119 to A.S.M., R01AI125563 and R01AI41158 to C.A.H., and R01EB029455 to J.B.S.), Department of Defense (W81XWH-18-1-0735 and W81WH-21-1-0892 to G.R., and W81XWH-21-1-0891 to J.B.S.), Juvenile Diabetes Research Foundation (1-INO-2020-923-A-N to J.B.S.), Czech Science Foundation (20-13029S to J.T.), Institute of Biotechnology of the Czech Academy of Sciences (RVO 86652036 to J.T.), EU Horizon project ReSHAPE to F.I., and the Mark Foundation for Cancer Research to D.M.P.. We also acknowledge the Czech Centre for Phenogenomics at the Institute of Molecular Genetics (supported by RVO 68378050 and MEYS CR LM2018126). D.V. is the recipient of an ARCS^®^ Foundation Metro-Washington Chapter Scholar award and a National Science Foundation Graduate Research Fellowship Program award. A.R.C. is the recipient of an Oxford-Bristol Myers Squibb Fellowship. F.I. is a Wellcome Trust CRCD Fellow (211122/Z/8). The content is solely the responsibility of the authors and does not necessarily represent the official views of the National Institutes of Health. **Figure 3 C** was created using Biorender.com.

## Author Contributions

D.V. and J.B.S. conceived the project and oversaw the design of all experiments. D.V., M.I., J.T., T.H., J.G., A.Y., J.S, J.B., J.A.P., E.G., A.R.C., L.S.C., A.G.D., B.T.O.-J., and L.M.T. designed, executed, and analyzed experiments. D.F., A.S.M., D.M.P., J.H., F.I., C.A.H., M.S.A., J.A.B., G.R., and J.B.S oversaw experiment design and analysis. D.V. and J.B.S. wrote the manuscript, and all other authors provided critical comments on the manuscript. All authors read and approved the final manuscript.

## STAR Methods

### Protein purification and expression

The published V_H_ and V_L_ sequences of F5111 (Trotta et al., 2018) were used to formulate the recombinant F5111 antibody on the human IgG1 lambda isotype platform. The heavy chain (HC) and LC of the F5111 antibody were separately cloned into the gWiz vector (Genlantis). Antibodies were expressed recombinantly in HEK 293F cells via transient co-transfection of plasmids encoding the HC and LC. HC and LC plasmids were titrated in small-scale co-transfection tests to determine optimal ratios for large-scale expression. HEK 293F cells were grown to 1.2×10^6^ cells/mL and diluted to 1.0×10^6^ cells/mL. Midiprepped DNA (1 mg total of HC and LC plasmids per liter of cells) and 2 mg per liter of cells of polyethyleneimine (PEI, Polysciences) were independently diluted to 0.05 and 0.1 mg/mL in OptiPro medium (ThermoFisher Scientific), respectively, and incubated at room temperature for 15 minutes. Equal volumes of DNA and PEI were mixed and incubated at room temperature for an additional 15 minutes. Subsequently, the diluted HEK 293F cells and 40 mL/L of DNA/PEI mixture were added to a shaking flask and incubated at 37°C and 5% CO_2_ with rotation at 125 rpm for 5 days. Secreted antibodies were purified from cell supernatants 5 days post-transfection via protein G agarose (ThermoFisher Scientific) affinity chromatography followed by SEC on a Superdex 200 Increase 10/300 GL column (GE Healthcare) on a fast protein liquid chromatography (FPLC) instrument, equilibrated in HEPES-buffered saline (HBS, 150 mM NaCl in 10 mM HEPES pH 7.3). Purity (>99%) was verified by SDS-PAGE analysis.

For F5111 IC production, the hIL-2 cytokine (residues 1-133) was fused at the N-terminus of the F5111 antibody LC, connected by either a flexible 15-amino acid (Gly_4_Ser)_3_ linker (F5111 IC LN15), a 25-amino acid (Gly_4_Ser)_5_ linker (F5111 IC LN25), or a 35-amino acid (Gly_4_Ser)_7_ linker (F5111 IC LN35). We prepared a plasmid encoding the hIL-2-fused F5111 LC into the gWiz vector (Genlantis). ICs were expressed and purified via transient co-transfection of HEK 293F cells with the F5111 HC and the hIL-2-fused F5111 LC plasmids, as described for the F5111 antibody. F5111 IC variants (all with a 35-amino acid linker) were generated in the same manner, with the indicated single-point mutation in either the HC or the hIL-2 fused LC construct. The following single-point mutations were made to the V_H_ sequence: Y35A, Y52A, Y54A, Y60A, and V103A. The following single-point mutations were made to the V_L_ sequence: Y33A, Y94A, and S96A. The sequence for the F5111.2 antibody and F5111.2 IC constructs was obtained from the following patent: (Rondon et al., 2015).

The Control IC was generated in the same manner as described for F5111 IC. Published V_H_ and V_L_ sequences of the FITC-E2 antibody (Honegger et al., 2005) were used to formulate the recombinant Control IC on the human IgG1 lambda isotype platform. hIL-2 (residues 1-133) was fused at the N-terminus of the LC, connected by a 35-amino acid (Gly_4_Ser)_7_ linker. Separate plasmids were constructed in the gWiz vector (Genlantis) encoding the Control HC and the hIL-2-fused Control LC. The Control IC was expressed via transient co-transfection of HEK 293F cells with the HC and hIL-2-fused LC plasmids. Purification proceeded as described for the F5111 antibody.

Antibody or IC constructs with Fc effector function knocked out were generated in the same manner as above, using a HC plasmid with the N297A mutation (Mimura et al., 2000, 2001; Saunders, 2019; Tao et al., 1993; Wang et al., 2018). In all studies, the F5111.2 antibody includes the N297A mutation. All *in vivo* studies were performed with constructs with Fc effector function knocked out unless otherwise noted.

The hIL-2 cytokine (residues 1-133) was cloned into the gWiz vector (Genlantis) with a C-terminal hexahistidine tag. Protein was expressed via transient transfection of HEK 293F cells, as detailed for antibody constructs, and purified via Ni-NTA affinity chromatography followed by SEC using a Superdex 200 Increase 10/300 GL column (GE Healthcare) on an FPLC instrument, equilibrated in HBS. Purity (>99%) was verified by SDS-PAGE analysis.

For expression of biotinylated hIL-2, and the extracellular domains of the hIL-2Rα (residues 1-214) and hIL-2Rβ (residues 1-214) receptor subunits sequences were cloned into the gWiz vector (Genlantis) with a C-terminal biotin acceptor peptide (BAP)-GLNDIFEAQKIEWHE followed by a hexahistidine tag. Proteins were expressed and purified via Ni-NTA affinity chromatography and then biotinylated with the soluble BirA ligase enzyme in 0.5 mM Bicine pH 8.3, 100 mM ATP, 100 mM magnesium acetate, and 500 mM biotin (MilliporeSigma). Excess biotin was removed by SEC on a Superdex 200 Increase 10/300 column (GE Healthcare) on an FPLC instrument, equilibrated in HBS. Complete biotinylation was verified via SDS-PAGE streptavidin shift assay.

### Cell lines

HEK 293F cells (ThermoFisher Scientific) were cultivated in Freestyle 293 Expression Medium (ThermoFisher Scientific) supplemented with 2 U/mL penicillin-streptomycin (Gibco). Unmodified YT-1 (Yodoi et al., 1985) and IL-2Rα^+^ (Kuziel et al., 1993) YT-1 human NK cells were cultured in RPMI complete medium (RPMI 1640 medium supplemented with 10% FBS, 2 mM L-glutamine, 1× minimum non-essential amino acids, 1 mM sodium pyruvate, 25 mM HEPES, and 100 U/mL penicillin-streptomycin [Gibco]). All cell lines were maintained at 37°C in a humidified atmosphere with 5% CO_2_.

### Yeast surface binding studies

For binding studies on yeast, hIL-2 (residues 1-133) or mIL-2 (residues 1-149) were cloned into the pCT3CBN yeast display vector (a variant of pCT302 (Boder and Wittrup, 1997) with an N-terminal yeast agglutinin protein (Aga2) fusion followed by a 3C protease site, a C-terminal myc epitope tag, and BamHI/NotI gene-flanking restriction sites). After induction for 48 hours, 1×10^5^ cells of IL-2-displaying yeast per well were transferred to a 96-well plate and incubated in PBE (PBS with 0.1% BSA and 1 mM EDTA) containing serial dilutions of recombinant F5111 antibody for 2 hours at room temperature. Cells were then washed and stained with anti-human IgG Fc APC (HP6017, BioLegend 409306, 1:50) in PBE for 15 minutes at 4°C. After a final wash, cells were analyzed for antibody binding using a CytoFLEX flow cytometer (Beckman Coulter). Background-subtracted and normalized binding curves were fitted to a first-order binding model, and K_D_ values were determined using GraphPad Prism. Studies were performed three times with similar results.

### Bio-layer interferometry binding measurements

Binding studies were performed using bio-layer interferometry on an OctetRED96® bio-layer interferometry instrument (Molecular Devices). Biotinylated hIL-2, hIL-2Rα, and hIL-2Rβ were immobilized to streptavidin-coated biosensors (Molecular Devices) in 0.45 μm filtered PBSA (PBS pH 7.2 containing 0.1% BSA). hIL-2 and hIL-2Rβ were immobilized at a concentration of 50 nM for 120 seconds and hIL-2Rα was immobilized at a concentration of 100 nM for 120 seconds. Once baseline measurements were collected in PBSA, binding kinetics were measured by submerging the biosensors in wells containing serial dilutions of the appropriate analyte for 300 seconds (association) followed by submerging the biosensor in wells containing only PBSA for 600 seconds (dissociation). hIL-2/F5111 or hIL-2/F5111.2 complexes were formed by incubating a 1:1 molar ratio of the F5111 antibody to hIL-2 for 60 minutes at 37°C and then diluting to the appropriate concentration. An irrelevant protein (the monoclonal antibody trastuzumab) was immobilized to a reference streptavidin biosensor for subtraction of non-specific binding. Tips were regenerated in 0.1 M glycine pH 2.7. Data was visualized and processed using the Octet® Data Analysis software version 7.1 (Molecular Devices). Equilibrium titration curve fitting and K_D_ value determination was implemented using GraphPad Prism, assuming all binding interactions to be first order. Experiments were reproduced two times with similar results.

### YT-1 human NK cell activation studies

Approximately 2×10^5^ IL-2Rα^+^ YT-1 or IL-2Rα^-^ YT-1 cells were plated in each well of a 96-well plate and resuspended in 20 μL of RPMI complete medium containing serial dilutions of either hIL-2, hIL-2/F5111 complexes, or IC. hIL-2/F5111 complexes were formed by incubating a 1:1 molar ratio of the F5111 antibody to hIL-2 for 60 minutes at 37°C and then diluting to the appropriate concentration. Cells were stimulated for 20 minutes at 37°C and immediately fixed by addition of paraformaldehyde (Electron Microscopy Sciences) to a final concentration of 1.5% and incubated for 10 minutes at room temperature. Permeabilization of cells was achieved by resuspension in 200 μL of ice-cold 100% methanol (MilliporeSigma) for 30 minutes at 4°C. Fixed and permeabilized cells were washed twice with PBSA and incubated with anti-pSTAT5 AlexaFluor 647 (pY694, BD Biosciences 562076, 1:50) diluted in 20 μL of PBSA for 2 hours at room temperature. Cells were then washed twice in PBSA and analyzed on a CytoFLEX flow cytometer (BeckmanCoulter). Dose-response curves were fitted to a logistic model and EC_50_ values were calculated using GraphPad Prism data analysis software after subtraction of the MFI of unstimulated cells and normalization to the maximum signal intensity. Experiments were conducted in triplicate and performed at least twice with similar results.

### Human PBMC and mouse splenocyte activation studies

For IL-2 induced pSTAT5 assays in human PBMCs, leukopaks containing de-identified whole blood were obtained from Anne Arundel Medical Blood Donor Center (Anne Arundel, Maryland, USA). Human PBMCs were isolated from whole blood by density gradient centrifugation using a standard Ficoll gradient (Ficoll Paque, MilliporeSigma) according to the manufacturer’s protocol. PBMCs were subjected to ACK red blood cell lysis (Quality Biological) and resuspended in PBS. Approximately 2×10^6^ cells/well were then plated into 96-well plate, pelleted, and resuspended in 40 μL of RPMI complete medium containing serial dilutions of the appropriate treatment. hIL-2/antibody complexes were formed by incubating a 1:1 molar ratio of hIL-2 to either the F5111 or F5111.2 antibody for 60 minutes at 37°C and then diluting to the appropriate concentration. Cells were stimulated for 20 minutes at 37°C and immediately fixed by addition of 160 μL of 1× TFP Fix/Perm buffer (Transcription Factor Phospho Buffer Set, BD Biosciences) and incubated at 4°C for 50 minutes. 40 μL 1× TFP Perm/Wash buffer (Transcription Factor Phospho Buffer Set, BD Biosciences) was then added to each well, and cells were then pelleted and washed again with 200 μL of 1× TFP Perm/Wash buffer. Permeabilization was achieved by resuspending the cells in 150 μL of Perm Buffer III (BD Biosciences) and incubating for 30 minutes at 4°C. Cells were then washed with 200 μL of 1× TFP Perm/Wash buffer and then resuspended in 50 μL of 1× TFP Perm/Wash buffer containing the following antibodies: anti-human CD3 APC-eFlour780 (UCHT1, ThermoFisher Scientific 47-0038-42, 1:50), anti-human CD4 PerCp-Cy5.5 (SK3, BD Biosciences 341654, 1:20), anti-human CD8 BV605 (SK1, BioLegend 344742, 1:50), anti-human IL-2Rα BV421 (M-A251, BD Biosciences 562442, 1:100), anti-human FOXP3 PE (236A/E7, BD Biosciences 560852, 1:50), anti-pSTAT5 AlexaFluor 647 (pY694, BD Biosciences 562076, 1:50), and anti-human CD127 Alexa Fluor 488 (eBioRDR5, ThermoFisher Scientific 53-1278-42, 1:50). Cells were incubated for 2 hours at room temperature and then washed twice with PBSA. Data were collected on a BD Biosciences LSRII flow cytometer (Becton Dickinson) and analyzed using FlowJo software (FlowJo, LLC). T_Reg_s were gated as CD3^+^CD4^+^IL-2Rα^High^FOXP3^High^ cells, CD8^+^ T cells were gated as CD3^+^CD8^+^ cells, and T_Conv_ cells were gated as CD3^+^CD4^+^FOXP3^-^ cells. pSTAT5 dose-response curves were fitted to a logistic model and EC_50_ values were calculated using GraphPad Prism data analysis software after subtraction of the MFI of unstimulated cells and normalization to the maximum signal intensity. Unless otherwise specified, PBMC activation experiments were conducted in triplicate and performed at least twice using PBMCs from independent donors.

For IL-2-induced pSTAT5 assays in murine lymphocytes, spleens from female NOD/ShiLtJ mice (The Jackson Laboratory) were collected and processed into a single-cell suspension followed by ACK red blood cell lysis (Quality Biological). Cells were seeded at 2×10^6^ cells/well into a 96-well plate, and the same protocol as for PBMC studies was followed using a different panel of antibodies: anti-mouse CD3 BV510 (145-2C11, BioLegend 100353, 1:100), anti-mouse CD4 APC-eF780 (RM4-5, ThermoFisher Scientific 47-0042-82, 1:100), anti-mouse CD8a AlexaFluor 488 (53-6.7, BD Biosciences 557668, 1:50), anti-mouse IL-2Rα PE (PC61.5, ThermoFisher Scientific 12-0251-83, 1:100), anti-mouse FOXP3 eFlour450 (FJK-16s, ThermoFisher Scientific 48-5773-82, 1:50), and anti-pSTAT5 Alexa Fluor 647 (pY694, BD Biosciences 562076, 1:50). Mouse splenocyte activation studies were conducted in triplicate.

### Quantification of IL-2Rα expression levels

Human PBMCs were isolated as described in the section “Human PBMC and mouse splenocyte activation studies.” Approximately 2×10^6^ human PBMCs/well or 0.2×10^6^ YT-1 cells/well were resuspended in 50 μL of PBS containing LIVE/DEAD™ Fixable Blue Dead Cell Stain Kit (ThermoFisher Scientific L34961, 1:1000) and stained for 15 minutes at 4°C. Cells were washed with PBSA, resuspended in 40 μL of RPMI, and then fixed by addition of 160 μL of 1× TFP Fix/Perm buffer (Transcription Factor Phospho Buffer Set, BD Biosciences) and incubated at 4°C for 50 minutes. 40 μL 1× TFP Perm/Wash buffer (Transcription Factor Phospho Buffer Set, BD Biosciences) was then added to each well, and cells were then pelleted and washed again with 200 μL of 1× TFP Perm/Wash buffer. Permeabilization was achieved by resuspending the cells in 150 μL of Perm Buffer III (BD Biosciences) and incubating for 30 minutes at 4°C. Cells were then washed with 200 μL of 1× TFP Perm/Wash buffer and resuspended in 50 μL of 1× TFP Perm/Wash buffer containing the following antibodies: anti-human CD3 APC-eFlour780 (UCHT1, ThermoFisher Scientific 47-0038-42, 1:50), anti-human CD4 PerCp-Cy5.5 (SK3, BD Biosciences 566316, 1:100), anti-human CD8 BV605 (SK1, BioLegend 344742, 1:50), anti-human IL-2Rα BV421 (M-A251, BD Biosciences 562442, 1:100), and anti-human FOXP3 PE (236A/E7, BD Biosciences 560852, 1:50). For the IL-2Rα FMO control, the same panel was used minus the addition of anti-human IL-2Rα BV421. Cells were incubated for 2 hours at room temperature and then washed twice with PBSA. Data were collected on a BD Biosciences LSRII flow cytometer (Becton Dickinson) and analyzed using FlowJo software (FlowJo, LLC). Staining was conducted in triplicate and performed twice using PBMCs from independent donors.

### Development of a multivalent binding model for IC signaling

The multivalent binding model used to predict cell type-specific signaling response to ICs was formulated as described in Tan et al. (Tan and Meyer, 2021). Each IL-2 molecule within the IC was assumed to bind to one free IL-2Rα and one IL-2Rβ/γ_c_ receptor; therefore, bivalent ICs were allowed to bind up to two IL-2Rα and IL-2Rβ/γ_c_ receptors each. Initial IL-2-IL-2Rα association was modeled as proceeding with the experimentally determined kinetics of monomeric ligand-receptor interaction. The affinities with which each IC initially interacted with IL-2Rβ/γ_c_ receptor dimer were inferred by fitting the binding model to our experimental *in vitro* pSTAT5 signaling data as described below, using least-squares fitting. Subsequent ligand-receptor binding interactions were modeled with an association constant proportional to the free receptor abundance and the monomeric affinity of receptor-ligand interaction multiplied by the scaling constant, 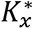. A single 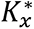 value was fit for all experiments and cell types when we fit our model to our *in vitro* pSTAT5 signaling data. To predict pSTAT5 response to IL-2 stimulation, we assumed that pSTAT5 is proportional to the amount of IL-2-bound IL-2Rβ/γ_c_, as complexes which contain these species actively signal through the JAK/STAT pathway. Scaling factors converting from predicted active signaling species to pSTAT5 abundance were fit to experimental data on a per-experiment and cell type basis. The abundance of each IL-2 receptor subunit on each cell type was assumed to be equal to previously published experimental human PBMC receptor quantitation data (Farhat et al., 2021).

### Immune cell subset expansion studies in NOD mice

For immune cell expansion studies in NOD mice (Tomala and Spangler, 2020), 8-week-old female NOD/ShiLtJ mice (4-5 mice per group, The Jackson Laboratory) were injected intraperitoneally (*i.p.*) for 4 consecutive days (Days 0, 1, 2, 3) with either 200 μL of PBS or the indicated treatment diluted in 200 μL of PBS. ICs were dosed at 8.2 μg per day (1.5 μg hIL-2 equivalence) and hIL-2/F5111.2 antibody complexes were formed by preincubating 1.5 μg hIL-2 with 6.6 μg F5111.2 antibody (1:2 antibody to cytokine molar ratio) in PBS for 60 min at 37°C. Mice were sacrificed 24 hours after the last dose (Day 4) by cervical dislocation, and spleens were harvested. Single-cell suspensions were prepared by mechanical homogenization and then subjected ACK red blood cell lysis (Quality Biological) and resuspended in PBS. Absolute count of splenocytes was assessed for each spleen. Approximately 2×10^6^ cells were used per sample. To assess viability, cells were resuspended in 50 μL of PBS containing eBioscience Fixable Viability Dye eFluor780 (ThermoFisher Scientific 65-0865-18, 1:2000) and stained for 15 minutes at 4°C. Cells were then washed with PBSA and resuspended in 50 μL of PBSA containing the following antibodies: anti-mouse CD3 BV510 (145-2C11, BioLegend 100353, 1:50), anti-mouse CD4 eFluor450 (RM4-5, ThermoFisher Scientific 48-0042-82, 1:100), anti-mouse CD8a AlexaFluor 488 (53-6.7, BD Biosciences 557668, 1:50), anti-mouse IL-2Rα PE (PC61.5, ThermoFisher Scientific 12-0251-83, 1:100), anti-mouse CD49b PE-Cy7 (DX5, BioLegend 108992, 1:100), and anti-mouse CD16/CD32 (2.4G2, BD Biosciences 553142, 1:100). Cells were stained for 30 minutes at 4°C. Cells were then washed with PBSA and resuspended in 200 μL of 1× eBioscience Fixation/Permeabilization buffer (ThermoFisher Scientific) and incubated for 45 minutes at 4 °C. 800 μL of 1× eBioscience Permeabilization buffer (ThermoFisher Scientific) was then added to each tube. Cells were subsequently resuspended in 50 μL of 1× eBioscience Permeabilization buffer containing anti-mouse FOXP3 APC (FJK-16s, ThermoFisher Scientific 17-5733-82, 1:80) and incubated for 45 minutes at 4°C. After a final wash and resuspension in PBSA, data were collected on a BD Biosciences LSRII flow cytometer (Becton Dickinson) and analyzed using FlowJo software (FlowJo, LLC). T_Reg_s were gated as CD3^+^CD4^+^IL-2Rα^High^FOXP3^High^ cells, CD8^+^ T cells were gated as CD3^+^CD8^+^ cells, T_Conv_ cells were gated as CD3^+^CD4^+^FOXP3^-^ cells, and NK cells were gated as CD3^-^CD49b^+^ cells. Statistical significance was determined by one-way ANOVA with Tukey post hoc test. Experiments were performed at least twice with similar results.

To quantify cell proliferation the same procedure as above was followed using the following panel: eBioscience Fixable Viability Dye eFluor780 (ThermoFisher Scientific 65-0865-18, 1:2000), CD3 BV510 (145-2C11, BioLegend 100353, 1:50), anti-mouse CD4 eFluor450 (RM4-5, ThermoFisher Scientific 48-0042-82, 1:100), anti-mouse CD8a BV570 (53-6.7, BioLegend 100739, 1:100), anti-mouse IL-2Rα PE (PC61.5, ThermoFisher Scientific 12-0251-83, 1:100), anti-mouse CD49b PE-Cy7 (DX5, BioLegend 108992, 1:100), anti-mouse CD16/CD32 (2.4G2, BD Biosciences 553142, 1:100), anti-mouse FOXP3 APC (FJK-16s, ThermoFisher Scientific 17-5733-82, 1:80), anti-mouse Ki-67 AlexaFluor 488 (16A8, BioLegend 652417, 1:100), anti-mouse CD44 PerCP-Cy5.5 (IM7, ThermoFisher Scientific 45-0441-82, 1:100), and anti-mouse Helios PE/Dazzle 594 (22F6, BioLegend 137231, 1:100). Statistical significance was determined by one-way ANOVA with Tukey post hoc test. Experiments were performed at least twice with similar results.

### Non-negative matrix factorization of immune cell subset expansion studies in NOD mice

The *in vivo* proliferative responses induced by the ICs were visualized using non-negative matrix factorization as implemented in scikit-learn (Pedregosa et al., 2011). The replicate average number of cells measured in NOD mice after treatment with each complex or IC was assembled. The log of the experimentally determined cell number was then taken, and number of each cell type measured during control (PBS) trials was subtracted from all other experimental measurements to obtain the log-fold changes in cell counts induced by each ligand in relation to the control trial. Non-negative matrix factorization with two components was subsequently performed on the processed matrix of cellular expansion data.

### Immune cell subset expansion studies in humanized mice

Human PBMCs (NHS Blood and Transport) were dyed with Violet Proliferation Dye 450 (BD Biosciences) before *i.p.* injection into male BRG mice (Charles River Laboratories). Mice received 5×10^6^ PBMCs on day 0 and on day 1 received a single *i.p.* dose of either PBS (n=6) or 8.2 μg (1.5 μg IL-2 equivalence) Y60A IC (n=5), Y33A IC (n=4), F5111 IC (n=4), or F5111.2 IC (n=5). Mice were sacrificed on day 4 and the PBMCs were retrieved by peritoneal lavage for flow cytometry analysis.

Cell were stained with Zombie NIR Fixable Dye (BioLegend 423105, 1:8000), anti-mouse CD45 APC-Cy7 (QA17A26, BioLegend 157617, 1:400), anti-mouse TER-119 APC-Cy7 (TER-119, BioLegend 116223, 1:400), anti-human CD3 BV605 (OKT3, BioLegend 317321, 1:200), anti-human CD56 PE (TULY56, ThermoFisher Scientific 12-0566-41, 1:200), anti-human CD4 PE-eFluor610 (RPA-T4, ThermoFisher Scientific 61-0049-42, 1:200), anti-human CD8 BV711 (SK1, BioLegend 344733, 1:200), anti-human IL-2Rα PE-Cy7 (M-A251, BD Biosciences 557741, 1:200), anti-human FOXP3 (259D, BioLegend 320214, 1:200), and anti-human Ki-67 FITC (20Rag1, ThermoFisher Scientific, 1:200). The eBioscience Foxp3/Transcription Factor Staining Buffer Set (ThermoFisher Scientific) was used for intracellular staining following the manufacturers protocol. Samples were collected on an Attune NxT flow cytometer and analyzed using FlowJo software (FlowJo, LLC). T_Reg_s were gated as CD3^+^CD4^+^IL-2Rα^High^FOXP3^High^ cells, CD8^+^ T cells were gated as CD3^+^CD8^+^ cells, and T_Conv_ cells were gated as CD3^+^CD4^+^FOXP3^-^cells. Murine cell populations (mCD45^+^mTER-119^+^) were excluded from the gating. Statistical significance was determined by one-way ANOVA with Tukey post hoc test. The experiment was performed twice with similar results.

### Immune cell subset expansion study dose titrations

Female C57BL/6 mice (2 mice per group, The Jackson Laboratory) were administered either 200 μL of PBS or increasing doses of F5111 diluted in 200 μL of PBS via *i.p.* injection daily for 4 days. The doses of F5111 IC used were 0.91 μg (0.167 μg IL-2 equivalence), 2.7 μg (0.5 μg IL-2 equivalence), 4.1 μg (0.75 μg IL-2 equivalence), 6.2 μg (1.125 μg IL-2 equivalence), and 8.2 μg (1.5 μg IL-2 equivalence). Mice were euthanized 24 hours after the last dose and spleens were harvested. Splenocytes were isolated by mechanical dissociation through 100 μm filters and subjected to ACK red blood cell lysis buffer. Cells were stained with anti-mouse CD45 PerCP-Cy5.5 (30-F11, BioLegend 103132, 1:1200), anti-mouse CD3 BV605 (17A2, BioLegend 100237, 1:80), anti-mouse CD4 APC (GK1.5, BioLegend 100411, 1:3200), anti-mouse CD8a BV786 (53-6.7, BD Biosciences 563332, 1:400), anti-mouse IL-2Rα BV421 (7D4, BD 564571, 1:200). Transcription Buffer Set (BD) was used for intracellular staining following the manufacturers protocol and then stained with anti-mouse FOXP3 FITC (FJK-16s, ThermoFisher Scientific 11-5773-82, 1:200). Samples were acquired on a BD Celesta and analyzed using FlowJo (FlowJo, LLC). T_Reg_s were gated as CD45^+^CD3^+^CD4^+^IL-2Rα^High^FOXP3^High^ cells. Statistical significance was determined by one-way ANOVA with Tukey post hoc test.

### Immune cell subset expansion study kinetics

8-week-old male C57BL/6J mice (3 mice per group, The Jackson Laboratory) were administered *i.p.* injections for 4 consecutive days (Days -3, -2, -1, 0) containing either 200 μL of PBS or 8.2 μg F5111 IC (1.5 μg hIL-2 equivalence) diluted in 200 μL of PBS. Spleens from the PBS-treated group were harvested one day post-treatment (Day 1). Spleens from the F5111 IC treated mice were harvested 1, 3, 5, and 7 days post-treatment (3 mice per harvest). The same protocol was implemented as described for the “Immune cell subset expansion studies in NOD mice” with the substitution of the following antibody panel: anti-mouse CD3 BV510 (145-2C11, BioLegend 100353, 1:100), anti-mouse NK-1.1 PerCp-Cy5.5 (PK136, BioLegend 108727, 1:100), anti-mouse CD4 eFluor450 (RM4-5, ThermoFisher Scientific 48-0042-82, 1:100), anti-mouse CD8a AlexaFluor 488 (53-6.7, BD Biosciences 557668, 1:100), anti-mouse IL-2Rα PE (PC61.5, ThermoFisher Scientific 12-0251-83, 1:100), anti-mouse FOXP3 APC (FJK-16s, ThermoFisher Scientific 17-5733-82, 1:80), and anti-mouse CD16/CD32 (2.4G2, BD Biosciences 553142, 1:100). T_Reg_s were gated as CD3^+^CD4^+^IL-2Rα^High^FOXP3^High^ cells, CD8^+^ T cells were gated as CD3^+^CD8^+^ cells, T_Conv_ cells were gated as CD3^+^CD4^+^FOXP3^-^ cells, and NK cells were gated as CD3^-^NK-1.1^+^ cells.

### Pharmacokinetic study

F5111 IC was labeled using 10-fold molar excess of N-hydroxysuccinimide (NHS)-Rhodamine (ThermoFisher Scientific), following the manufacturer’s protocol. Non-reacted NHS-rhodamine was removed by SEC on a Superdex 200 Increase 10/300 GL column (GE Healthcare), equilibrated in HBS. Protein concentration and degree of labeling were calculated according to the manufacturer’s protocol.

Prior to treatment, a small volume of blood was collected from the tail vein of 8-week-old male C57BL/6J mice (5 mice, The Jackson Laboratory). Mice were then administered retro-orbital injections of 2 mg/kg (∼0.36 mg/kg hIL-2 equivalence, ∼10 μg hIL-2 total/mouse) F5111 IC diluted in 200 μL of PBS. Blood was collected from the tail vein of each mouse at 5 minutes, 20 minutes, 40 minutes, 1 hour, 2 hours, 4 hours, 8 hours, 24 hours, 48 hours, 72 hours, 96 hours, and 120 hours. At each time point, blood was collected in EDTA-coated tubes and centrifuged at 500×g for 5 minutes. The plasma was collected and stored at 4°C for later analysis. After all samples were collected, the plasma was diluted in PBS (1:10 dilution), and 100 μL of diluted sample was added to a 96 well black clear-bottom plate. Fluorescence (Excitation/Emission: 540/590 nm) was measured on a BioTek Synergy 2 instrument, and blood samples collected before treatment were used for background subtraction. Standard curves were generated from fluorescent measurements of rhodamine-labeled F5111 IC at concentrations ranging from 3 μg/mL to 0.023 μg/mL (2-fold dilutions) and this standard curve was used to determine protein concentration of the collected samples. Serum half-life was calculated using a two-phase decay model in GraphPad Prism.

### *In vitro* T_Reg_ suppression assay

Female C57BL/6 CD45.1; RFP-FOXP3 mice (The Jackson Laboratory) were treated with either 200 μL of PBS (n=3) or 6.2 μg F5111 IC in 200 μL of PBS (n=2) *i.p.* every day for 4 days. 24 hours after the last dose, T_Reg_s were isolated from spleens from each group for the T_Reg_ suppression assay. Spleens from female C57BL/6 CD45.2 mice (n=5) were taken for isolation of T_Conv_ cells.

Pooled mouse spleens were digested and CD4 cells were positively selected from all mouse groups (Miltenyi, 130-117-043). CD4^+^ cell suspensions were stained with either a T_Reg_ isolation panel: anti-mouse CD45.1 APC-Cy7 (A20, BioLegend 110716, 1:400), anti-mouse CD45.2 PerCp-Cy5.5 (104, BioLegend 109828, 1:400), anti-mouse CD4 APC (GK1.5, BioLegend 100411, 1:3200), anti-mouse CD8a BV786 (53-6.7, BD 563332, 1:400), anti-mouse IL-2Rα BV421 (7D4, BD 564571, 1:200), or a T_Conv_ isolation panel: anti-mouse CD45.1 APC-Cy7 (A20, BioLegend 110716, 1:400), anti-mouse CD45.2 PerCp-Cy5.5 (104, BioLegend 109828, 1:400), anti-mouse CD4 APC (GK1.5, BioLegend 100411, 1:3200), anti-mouse CD8a BV786 (53-6.7, BD 563332, 1:400), anti-mouse IL-2Rα BV421 (7D4, BD 564571, 1:200), anti-mouse CD62L PE (MEL-14, BioLegend 104408, 1:200), and anti-mouse CD44 AlexaFluor 700 (IM7, BioLegend 103026, 1:400). CD45.1^+^CD4^+^CD8^-^IL-2Rα^High^RFP-FOXP3^+^ T_Reg_s were sorted from F5111 treated and PBS treated mice. CD45.2^+^CD4^+^CD8^-^IL-2Rα^-^CD62L^High^CD44^Low^ naïve T_Conv_ cells were sorted for effector cells. Cell sorting occurred on a BD FACSAria^TM^.

For division tracking, T_Conv_ cells were labeled with CellTrace^TM^ Violet (CTV, ThermoFisher Scientific) by incubating cells for 16 minutes at 37°C at a cell density of 10^6^/mL in PBSA. Cells were vortexed every 4 minutes of staining. Reaction was quenched in ice cold RPMI 1640 medium (Invitrogen) with 10% FBS (MilliporeSigma).

Cells were cultured in RPMI 1640 medium (Invitrogen) supplemented with non-essential amino acids (Gibco), 1 mM sodium pyruvate (MilliporeSigma), 10 mM HEPES (Gibco), 100 U/mL Penicillin, 100 μg/ml streptomycin (Quality Biologic), 50 μM 2-mercaptoethanol (Gibco), 2 mM L-glutamine (Gibco) and 10% FBS (MilliporeSigma). Cells were incubated at 37 °C in 5% CO_2_. The T_Reg_ suppression assay was performed as follows: 50,000 T_Conv_ cells were co-cultured with a varying ratio of T_Reg_s in 96-well U-bottom plate. Anti-CD3/anti-CD28 coated microbeads (Dynabeads Mouse T Activator, ThermoFisher Scientific, 11452D) were added at a 1:2 (bead: cell) ratio with final number of cells in each well. Proliferation was analyzed by flow cytometry done on a BD FACSCelesta^TM^ after 4 days. Cells were washed in PBS and stained with anti-mouse CD45.1 APC-Cy7 (A20, BioLegend 110716, 1:400) and anti-mouse CD45.2 PerCp-Cy5.5 (104, BioLegend 109828, 1:400) to differentiate T_Conv_ cells from T_Reg_s. A minimum number of 20,000 CD45.2^+^CTV^+^ cells were collected and analyzed with FlowJo software (FlowJo, LLC). The percent suppression is calculated using the following formula: ((%CTV^-^ of T_Conv_ cells alone - %CTV^-^ of T_Conv_ cells treated with T_Reg_)/%CTV^-^ of T_Conv_ cells alone)*100 (Collison and Vignali, 2011). CTV is gated on unstimulated T_Conv_ with all divisions beyond unstimulated condition considered CTV^-^. Experiment was conducted with technical replicates and repeated twice with similar results. Statistical significance at each ratio was determined by two-tailed unpaired Student *t* test.

### Comparison of cT_Reg_ versus eT_Reg_ expansion

8-week-old male C57BL/6 mice (5 mice per group, Taconic Biosciences) were injected *i.p.* every 48 hours (days 0, 2, and 4) with either 200 μL of 1× Dulbecco’s PBS (DPBS, Corning) or 8.2 μg (1.5 μg IL-2 equivalence) Control IC or F5111 IC diluted in 200 μL of 1× DPBS. The mice were sacrificed on day 6 and their splenocytes were harvested for analysis via flow cytometry. Splenocyte single cell suspensions were isolated by mechanically processing harvested spleens through 70 µm nylon filters. The suspensions were spun down at 300×g for 5 minutes, and then resuspended in 1 mL of lysis buffer (0.846% solution of NH_4_Cl) for 5 minutes to lyse red blood cells. The cells were then washed in 10 mL of complete RPMI and then resuspended to 30e^6^ cells/ml on ice for staining.

5e^6^ cells per sample were placed in a 96 well round bottom plate and washed with ice cold 1× DPBS. The cells were resuspended in 50 µl of Ghost Dye^TM^ Violet 510 viability dye (Tonbo Biosciences 13-0870-T100, 1:200) reconstituted in 1× DPBS for 20 minutes on ice and then washed in 0.2% FACS buffer (1× DPBS, 2 g/L BSA, 0.5 M EDTA). The cells were then resuspended in 50 µl of volume of 0.2% FACS buffer containing anti-mouse CD16/CD32 (2.4G2, Bio X Cell BP0307, 1 µg/mL) and rat IgG Isotype Control (ThermoFisher Scientific 10700, 1:200) for 30 minutes on ice. The cells were washed in 0.2% FACS buffer, and then incubated for 30 minutes on ice in 50 µl volume of antibody cocktail composed of: anti-mouse CD4 BUV563 (GK1.5, BD Biosciences 612923, 1:200), anti-mouse CD8a BUV615 (53-6.7, BD Biosciences 613004, 1:200), anti-mouse CD11a BUV805 (2D7, BD Biosciences 741919, 1:200), anti-mouse IL-2Rα BV785 (PC61, BioLegend 102051, 1:200), anti-mouse CD27 BV650 (LG.3A10, BioLegend 124233, 1:200), anti-mouse CD44 BV570 (1M7, BioLegend 103037, 1:100), anti-mouse CD69 BUV737 (H1.2F3, BD Biosciences 612793, 1:200), anti-mouse IL-2Rβ BUV661 (TM-β1, BD Biosciences 741493, 1:200), anti-mouse KLRG1 BUV395 (2F1, BD Biosciences 740279, 1:300), anti-mouse ICOS APC/Fire 750 (C398.4A, BioLegend 313536, 1:200), and anti-mouse PD-1 BV421 (29F.1A12, BioLegend 135218, 1:250) in 0.2% FACS buffer supplemented with Brilliant Stain Buffer (BD Biosciences, 1:10). The cells were washed in 0.2% FACS buffer and re-suspended in 100 µl 1× eBioscience Fixation/Permeabilization buffer (ThermoFisher Scientific) for 4 hours at 4⁰C. The cells were then washed twice in 1× eBioscience Permeabilization buffer (ThermoFisher Scientific), and then resuspended in 50 µl of 1× eBioscience Permeabilization buffer containing: anti-mouse BCL-2 AlexaFluor 647 (BCL/10C4, BioLegend 633510, 1:200), anti-mouse CD3 BV750 (17A2, BioLegend 100249, 1:200), anti-mouse CTLA-4 APC-R700 (UC10-4F10-11, BD Biosciences 565778, 1:300), anti-mouse FOXP3 PE-Cy5.5 (FJK-16s, ThermoFisher Scientific 35-5773-82, 1:200), anti-mouse Helios PE/Dazzle 594 (22F6, BioLegend 137231, 1:200), anti-mouse Ki-67 AlexaFluor 488 (B56, BD Biosciences 558616, 1:400), and anti-mouse T-bet PE-Cy5 (4B10, ThermoFisher Scientific 15-5825-82, 1:200) for 2 hours at 4⁰C. The cells were then washed with 1× eBioscience Permeabilization buffer twice, and then resuspended in 500 µl 0.2% FACS buffer for flow cytometric analysis. Fluorescence minus one (FMOs) controls and individual single color controls and were created using pooled remaining splenocytes and were prepared simultaneously with experimental samples. The prepared and stained cells were analyzed using a FACS Symphony A5 (BD Biosciences) running BD FACSDiva v9.0 (BD Biosciences). Raw flow cytometry data was compensated and analyzed using FlowJo (FlowJo, LLC). cT_Reg_s were gated as CD4^+^FOXP3^+^IL-2Rα^High^BCL-2^High^ and eT_Reg_s were gated as CD4^+^FOXP3^+^IL-2Rα^Low^BCL-2^Low^. Statistical significance was determined separately for cT_Reg_s and eT_Reg_s by one-way ANOVA with a Tukey post hoc test.

### *Toxoplasma gondii* infection mouse model

ME49 strain *Toxoplasma gondii* cysts were obtained from neural tissue harvested from chronically infected CBA/ca mice and resuspended to 125 cysts/mL in 1× DPBS (Corning). Cohorts of male C57BL/6 mice (5 mice per group, Taconic Biosciences) were infected via *i.p.* injection at 8-9 weeks of age using 200 μL of diluted ME49 cysts for an infection dose of 25 cysts/mouse. The disease-free control group did not receive ME49 cysts and instead received an injection of 1× DPBS. Mice were then injected *i.p.* with either 200 μL of 1× DPBS (control group and PBS) or 8.2 μg (1.5 μg IL-2 equivalence) Control IC or F5111 IC diluted in 200 μL of 1× DPBS every 24 hours after infection on days 1, 2, 3, 4, and 5. The mice were sacrificed on day 10 of infection, and splenocytes were harvested for analysis via flow cytometry, the right lateral lobe of the lungs and liver were frozen at −80°C for DNA isolation, and the left lateral lobe of the liver was stored in 10% buffered formalin (Jansen Pharmaceuticals) for histological analysis.

The staining and flow cytometry analysis was performed as described in the “Comparison of cT_Reg_ versus eT_Reg_ expansion” methods except anti-mouse CD27 was excluded and the following antibodies were added to the panel: anti-mouse CXCR3 BV650 (CXCR3-173, BioLegend 126531, 1:150), anti-mouse NK1.1 BV711 (PK136, BioLegend 108245, 1:300), and anti-mouse NKp46 BV605 (29A1.4, BioLegend 137619, 1:300). Additionally, toxoplasma-specific tetramers for CD4^+^ (Tetramer I-A(b) AS15 PE, NIH, AVEIHRPVPGTAPPS, 1:400) and CD8^+^ (Tetramer H2k(b) Tgd057 PE, NIH, SVLAFRRL, 1:400) T cells were included. T_Reg_s were gated as CD3^+^CD4^+^IL-2Rα^High^FOXP3^High^ cells, CD8^+^ T cells were gated as CD3^+^CD8^+^ cells, and T_Conv_ cells were gated as CD3^+^CD4^+^FOXP3^-^ cells.

Harvested left lateral liver lobes were kept submerged in 10% buffered formalin for 48 hours then mounted in paraffin, cut, and hematoxylin and eosin (H&E) stained by the University of Pennsylvania Comparative Pathology Core.

Total DNA was isolated from frozen sections of the liver, heart, and lungs using a DNeasy Isolation Kit (Qiagen) according to manufacturer protocol. Isolated DNA concentration was quantified via NanoDrop spectrophotometry (ThermoFisher Scientific) and then individual samples were diluted in DNase/RNase-free distilled water (ThermoFisher Scientific) to equivalent concentrations for qPCR analysis (lungs 40 ng/μL, livers 50 ng/μL). Parasite DNA quantity was assessed in triplicate from the individual prepared DNA samples via qPCR using toxoplasma specific primers: (forward) 5’-TCCCCTCTGCTGGCGAAAAGT-3’ and (reverse) 5’- AGCGTTCGTGGTCAACTATCGATT G-3’ with Power SYBR Green master mix (Applied Biosystems). The reaction conditions for the qPCR were: holding phase of 2 minutes at 50°C, and then 10 minutes at 95°C (occurs only once); followed by 50 cycles of PCR phases of 15 seconds at 95°C, and 1 minute at 60°C. The qPCR reaction was performed on a ViiA 7 Real-Time PCR system operating ViiA 7^TM^ Software. Analysis of parasite qPCR data was performed in Microsoft Excel version 2202 (Build 14931.20120).

Statistical significance was determined by one-way ANOVA with Tukey post hoc test. The experiment was performed twice with similar results.

### DSS–induced colitis mouse model

Female, 8-week-old, BALB/c mice (6 mice per group, Czech Centre for Phenogenomics) were injected *i.p.* daily for 7 days (days 0-6) with either 200 μL of PBS (control group and PBS) or 8.2 μg (1.5 μg hIL-2 equivalence) Control IC or F5111 IC diluted in 200 μL of PBS. For the complex, 1.5 μg hIL-2 was complexed with 6.6 μg F5111.2 antibody (1:2 antibody to cytokine molar ratio) in 200 μL of PBS for 60 min at 37°C. Beginning on day 7, all groups except for the disease-free control group were administered 3% DSS (MW = 36,000 – 50,000; MP Biomedicals) in their drinking water to induce colitis. Mice weight, stool consistency, and rectal bleeding were measured daily and scores were assigned for each category (Cooper et al., 1993). The weight loss score was calculated based on initial weight as follows: 0 (<1% weight loss); 1 (1-5% weight loss); 2 (5-10% weight loss); 3 (10-20% weight loss); 4 (>20% weight loss). Stool consistency was scored as follows: 0 (normal stool); 2 (loose stool); 4 (diarrhea/liquid stool). Rectal bleeding was scored as follows: 0 (no presence of blood); 4 (blood observed). The disease activity index was calculated as the average of the weight loss, stool consistency, and rectal bleeding. On day 15, mice were sacrificed and entire colons were removed (from cecum to anus). Colon length was measured, and shortening was used as an indirect marker of pathological inflammation. Distal colon sections were fixed in Carnoy’s solution and embedded in paraffin. Histological scoring of paraffin-embedded and hematoxylin and eosin-stained transversal colon sections was implemented in a blinded manner using a weighted score, ranging from 0 (no signs of inflammation) to 4 (severe inflammation). Statistical significance was determined by one-way ANOVA with Tukey post hoc test. The experiment was performed two times with similar results.

### Immune checkpoint inhibitor-induced diabetes mellitus mouse model

8-week-old female NOD/ShiLtJ mice (the Jackson Laboratory) were treated *i.p.* with either 200 μL of PBS (control group and PBS) or 8.2 μg (1.5 μg IL-2 equivalence) Control IC or F5111 IC diluted in 200 μL of PBS from day −3 to 0 prior to initiating anti-PD-1 treatment. Starting on day 0 (4 hours after the last IC treatment), mice were treated *i.p.* every 4 days with either PBS (control group) or 200 μg anti-PD-1 antibody (clone RMP1-14, Bio X Cell) for a total of four or five doses. Non-fasting blood glucose was monitored by using a OneTouch® Ultra® 2 glucometer. Diabetes onset was considered to have occurred when non-fasting blood glucose concentration exceeded 250 mg/dL for two consecutive measurements. Statistical significance was determined by pairwise comparisons using the Log-rank (Mantel-Cox) test.

### Mice

NOD/ShiLt and C57BL/6 mice were purchased from The Jackson Laboratory unless otherwise specified. Animals were housed in specific pathogen-free conditions and experiments conducted in accordance with National Institutes of Health guidelines, and approval by the Johns Hopkins University Animal Care and Use Committee.

For the immune cell subset expansion studies in humanized mice, BRG mice were obtained from Charles River Laboratories and were maintained under specific pathogen-free conditions in the Biomedical Services Unit of the University of Oxford (Oxford, United Kingdom). All experiments in this study were performed using protocols approved by the Committee on Animal Care and Ethical Review at the University of Oxford and in accordance with the UK Animals (Scientific Procedures) Act 1986. Human tissue samples taken by NHS Blood and Transport with informed consent and ethical approval from the Oxfordshire Research Ethics Committee, study number 07/H0605/130.

For the comparison of cT_Reg_ versus eT_Reg_ expansion and the *Toxoplasma gondii* infection mouse model, 6-week-old C57BL/6 mice were purchased from Taconic Biosciences (Rensselaer, NY, USA) and kept in the University of Pennsylvania Department of Pathobiology vivarium until they reached 8-9 weeks of age for experimental use. Mice housed in the University of Pennsylvania Department of Pathobiology vivarium were maintained under institutional guidelines of 12-hour light/dark cycles, temperature ranges of 68-77°F and humidity ranging from 35-55%. Ethical oversight of all animal use in this study was approved by the University of Pennsylvania Institutional Animal Care and Use Committee.

For the DSS-induced colitis mouse model, BALB/c mice were acquired from the colony kept at the Czech Centre for Phenogenomics, Prague, Czech Republic. They were housed and handled according to the institutional committee guidelines with free access to food and water. Animal experiments were approved by the Animal Care and Use Committee of the Institute of Molecular Genetics and were in agreement with local legal requirements and ethical guidelines.

For the immune checkpoint inhibitor-induced diabetes mellitus mouse model, NOD/ShiLtJ mice were purchased from the Jackson Laboratory and maintained in the UCSF specific pathogen-free animal facility in accordance with guidelines established by the Institutional Animal Care and Use Committee and Laboratory Animal Resource Center.

### Statistical analysis

All statistical analysis was performed using GraphPad Prism. The number of replicates, number of mice, and type of analysis performed is described in each of the above sections where applicable. Significance between all groups is not always shown in figures, for full analysis see **Table S6**.

### Flow cytometry antibodies and fluorescent dyes

A full list of antibodies and fluorescent dyes used for the above experiments can be seen in **Table S7**.

## Supplemental Material

### Supplemental Figure Legends

**Figure S1.**
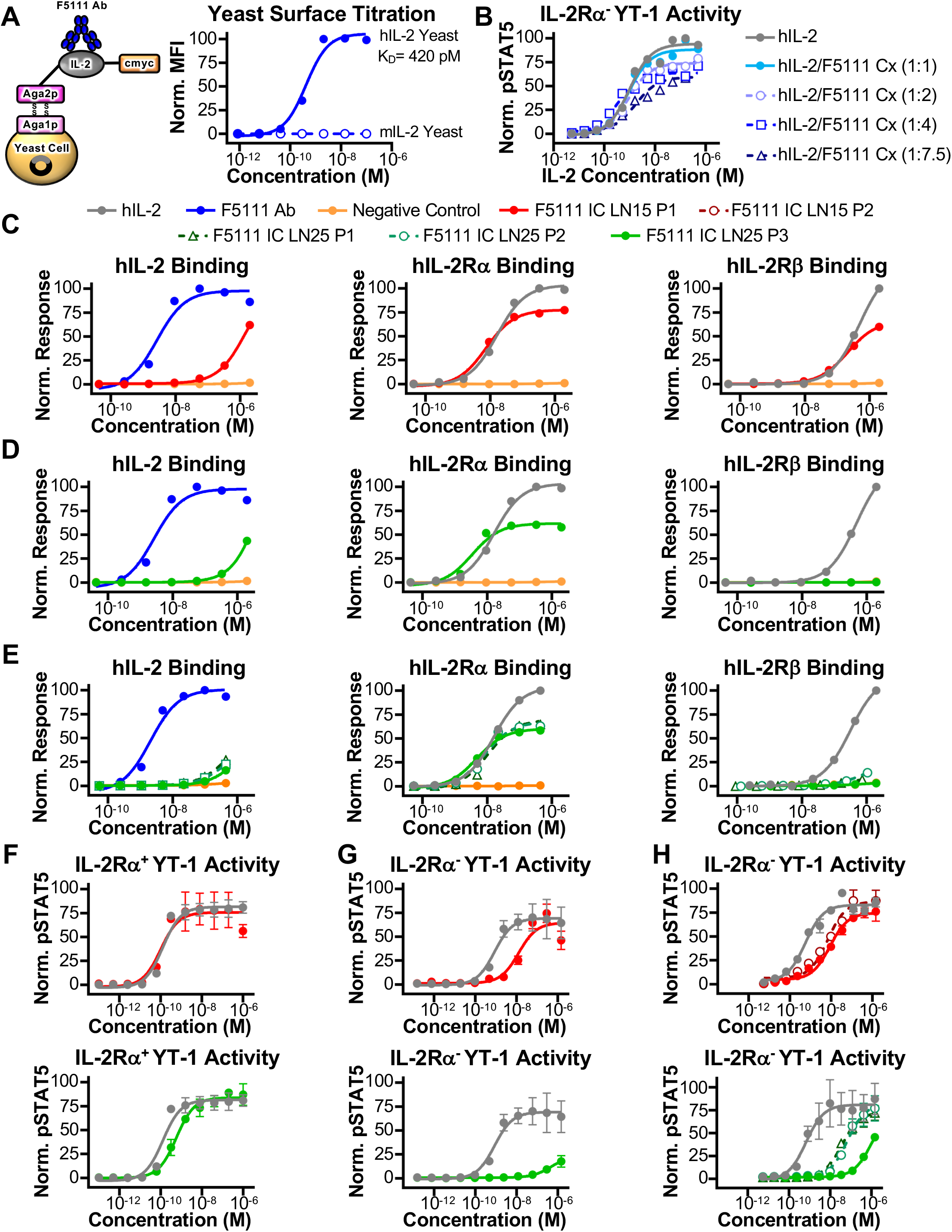
Increasing IC linker length improves T_Reg_ bias. **(A)** Binding of the F5111 antibody (Ab) to yeast-displayed hIL-2 or mIL-2. Fitted equilibrium dissociation constant (K_D_) is shown. **(B)** STAT5 phosphorylation response of IL-2Rα^-^ YT-1 human NK cells stimulated with either hIL-2 or hIL-2/F5111 complex (Cx) at varying molar ratios of cytokine to antibody. **(C-E)** Equilibrium biolayer interferometry-based titrations of hIL-2, F5111 antibody (Ab), Negative Control (trastuzumab), F5111 IC LN15 P1, and F5111 IC LN25 P1, P2, and P3 binding to immobilized hIL-2 (left), immobilized hIL-2Rα (middle), and immobilized hIL-2Rβ (right). **(F)** STAT5 phosphorylation response of IL-2Rα^+^ YT-1 human NK cells stimulated with either hIL-2, F5111 IC LN15 P1 (top), or F5111 IC LN25 P3 (bottom). **(G)** STAT5 phosphorylation response of IL-2Rα^-^ YT-1 human NK cells stimulated with either hIL-2, F5111 IC LN15 P1 (top), or F5111 IC LN25 P3 (bottom). **(H)** STAT5 phosphorylation response of IL-2Rα^-^ YT-1 human NK cells stimulated with either hIL-2, F5111 IC LN15 P1 and P2 (top), or F5111 IC LN25 P1, P2, and P3 (bottom). Data in **(F-H)** represent mean ± SD (n=3). See also **Tables S2 and S3**.

**Figure S2.**
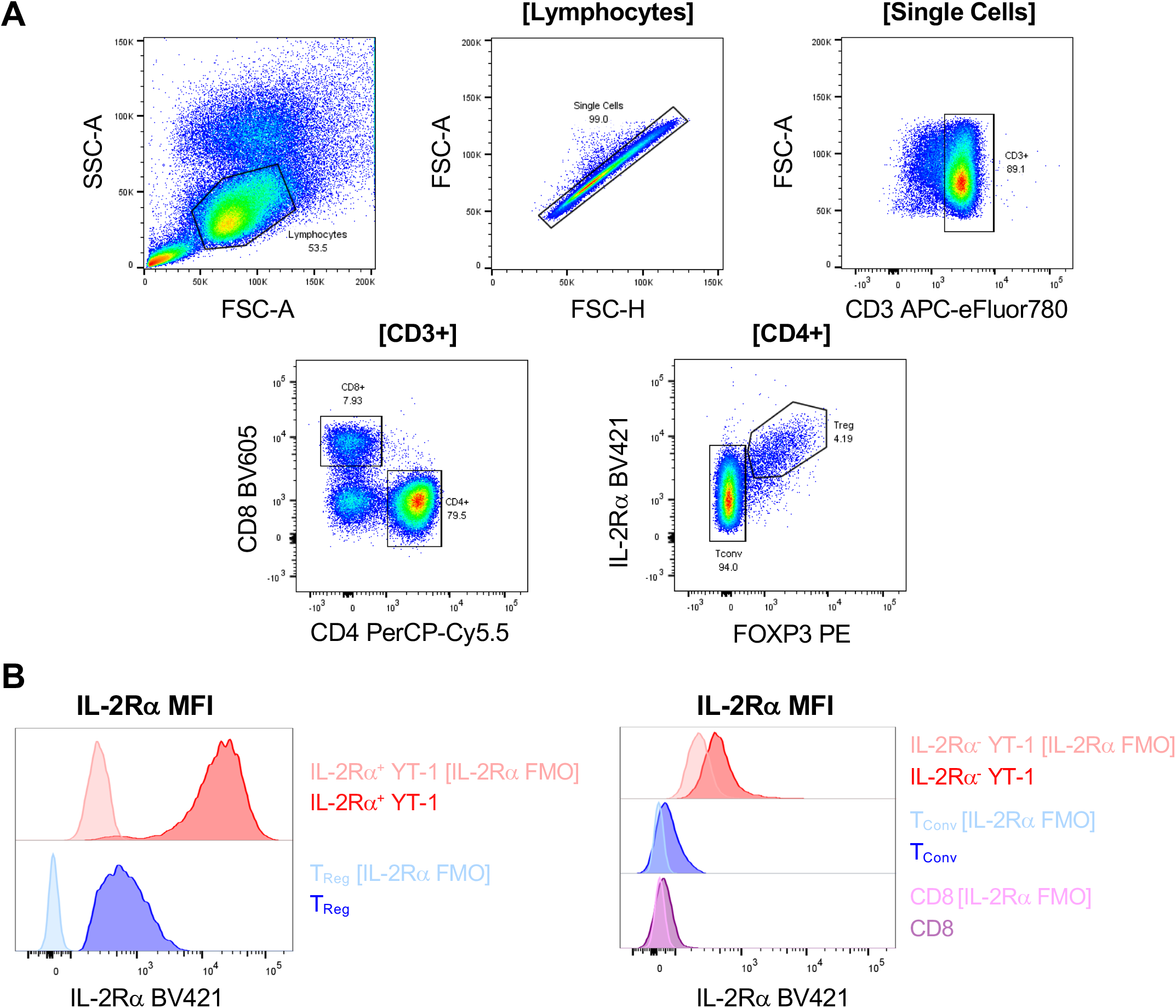
IL-2Rα^+^ YT-1 cells express higher levels of IL-2Rα than human T_Reg_s. **(A)** Representative flow cytometry plots illustrating the gating strategy used for human PBMCs. **(B)** Representative histograms illustrating IL-2Rα MFI of IL-2Rα^+^ YT-1 cells and human T_Reg_s (left) and IL-2Rα^-^ YT-1 cells, human CD8^+^ T cells, and human T_Conv_ cells (right) as compared to the IL-2Rα fluorescence minus one (FMO) controls.

**Figure S3.**
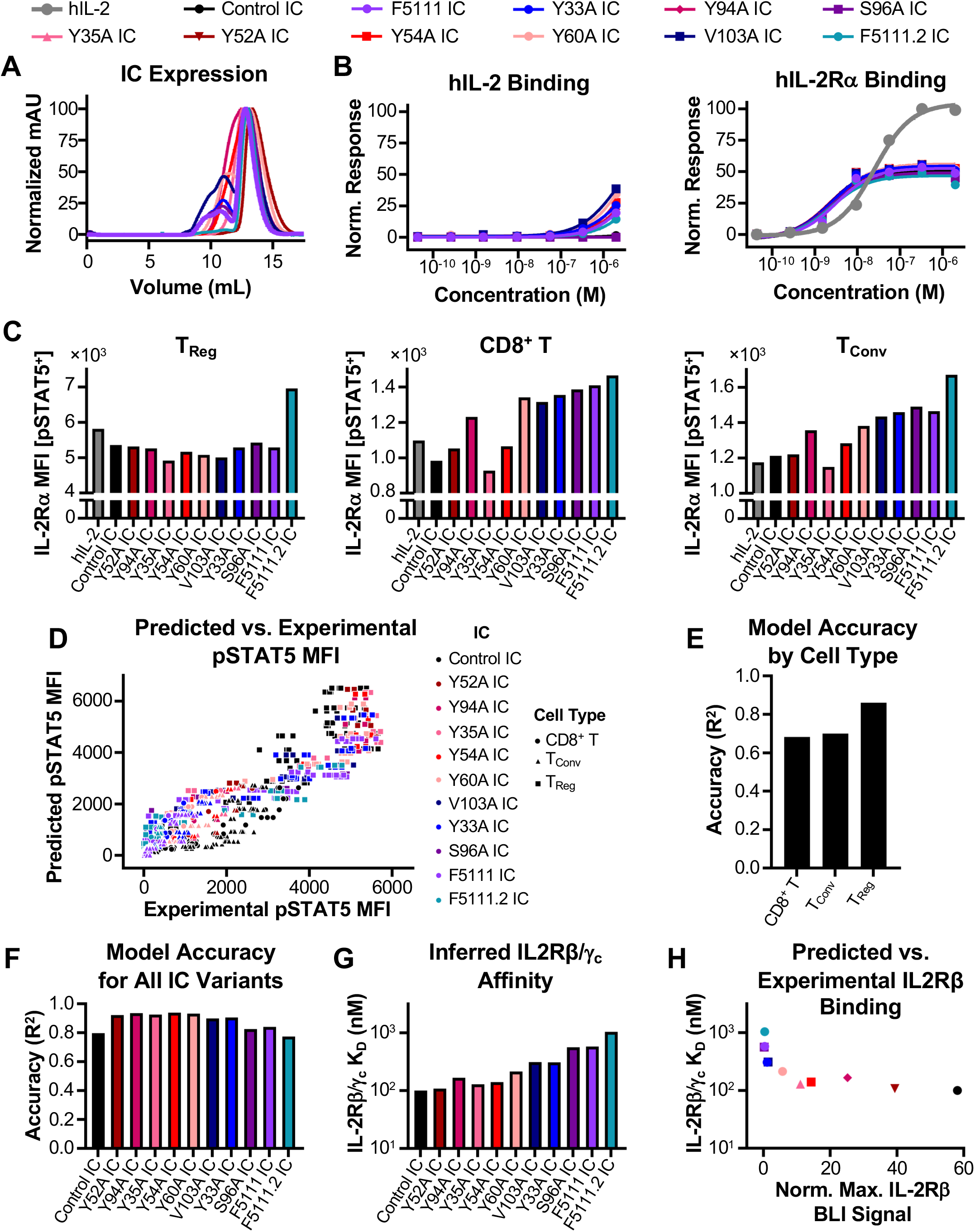
Characterization of F5111 IC variants. **(A)** Overlay of SEC traces for F5111 IC variants. **(B)** Equilibrium biolayer interferometry-based titrations of hIL-2, Control IC, and F5111 IC variants binding to immobilized hIL-2 (left) and immobilized hIL-2Rα (right). Binding to immobilized hIL-2 was normalized based on the binding of the F5111.2 antibody (**Figure S5C, left**). **(C)** IL-2Rα MFI within the pSTAT5^+^ population of T_Reg_ (left), CD8^+^ T (middle), and T_Conv_ (right) cell populations within human PBMCs stimulated with either hIL-2, Control IC, or F5111 IC variants at treatment concentrations of 2 nM (left) and 200 nM (middle and right). **(D)** Predicted vs. experimentally measured pSTAT5 MFI for all cell types, ICs, and concentrations modeled. Each point represents a single experimental measurement (n=1). **(E)** Model accuracy delineated by cell type for all ICs. **(F)** Model accuracy delineated by treatment for all ICs. Accuracies are calculated as a Pearson’s correlation R^2^. **(G)** Inferred IL2Rβ/γ_c_ equilibrium dissociation constants (K_D_, nM) for each IC. **(H)** Inferred IL2Rβ/γ_c_ equilibrium dissociation constants (K_D_, nM) compared to the maximum normalized IL2Rβ biolayer interferometry (BLI) signal experimentally measured for each IC. See also **Table S2**.

**Figure S4.**
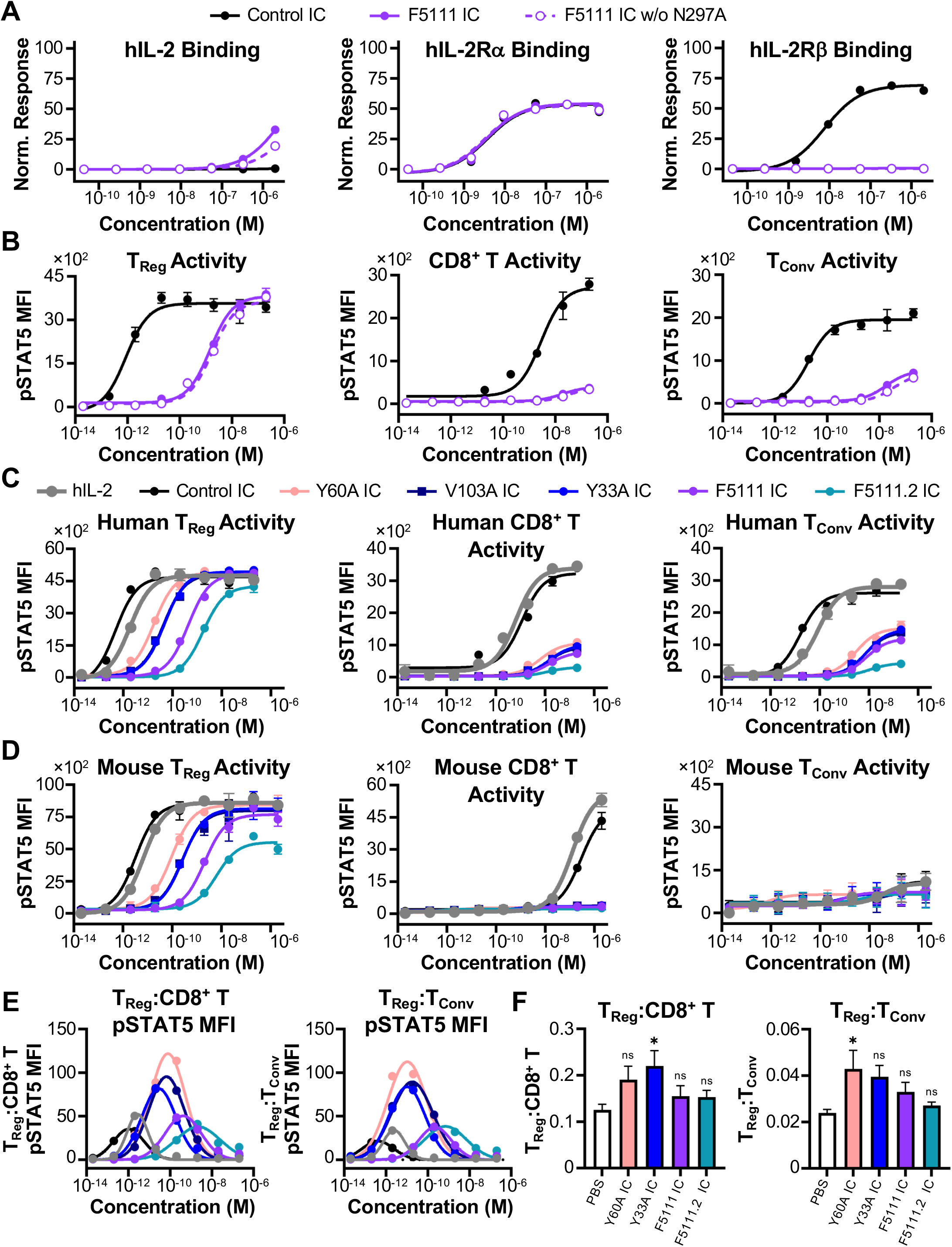
The N297A mutation does not impact *in vitro* function and IC variants modulate T_Reg_ bias *in vitro* and *in vivo*. **(A)** Equilibrium biolayer interferometry-based titrations of Control IC, F5111 IC, and F5111 IC without the N297A mutation (w/o N297A, intact effector function) binding to immobilized hIL-2 (left), immobilized hIL-2Rα (middle), and immobilized hIL-2Rβ (right). Binding to immobilized hIL-2 was normalized based on the binding of the F5111.2 antibody (**Figure S5C, left**). **(B)** STAT5 phosphorylation response of T_Reg_ (left), CD8^+^ T (middle), and T_Conv_ (right) cell populations from human PBMCs stimulated with either Control IC, F5111 IC, or F5111 IC w/o N297A. Data represent mean ± SD (n=3). **(C)** STAT5 phosphorylation response of T_Reg_ (left), CD8^+^ T (middle), and T_Conv_ (right) cell populations within human PBMCs stimulated with either hIL-2, Control IC, Y60A IC, V103A IC, Y33A IC, F5111 IC, or F5111.2 IC (all with N297A mutation). Data represent mean ± SD (n=3). **(D)** STAT5 phosphorylation response of T_Reg_ (left), CD8^+^ T (middle), and T_Conv_ (right) cells isolated from spleens of NOD mice and stimulated with either hIL-2, Control IC, Y60A IC, V103A IC, Y33A IC, F5111 IC, or F5111.2 IC (all with N297A mutation). Data represent mean ± SD (n=3). **(E)** Average pSTAT5 MFI ratio of human T_Reg_:CD8^+^ T (left) and human T_Reg_:T_Conv_ (right) for each IC determined at each stimulation concentration from the experiment shown in **(C)**. **(F)** BRG mice were administered 5×10^6^ human PBMCs (*i.p.*) on day 0 and then on day 1 were treated (*i.p.*) with either PBS (n=6) or 8.2 μg (1.5 μg IL-2 equivalence) Y60A IC (n=5), Y33A IC (n=4), F5111 IC (n=4), or F5111.2 IC (n=5). Cells were collected by lavage from the peritoneum on day 4 and ratios of human T_Reg_:CD8^+^ T (left) and T_Reg_:T_Conv_ (right) were evaluated. Data represent mean + SEM. Statistical significance was determined by one-way ANOVA with a Tukey post hoc test. Only statistical significance compared to PBS is shown on the plots. All statistical data are provided in **Table S6**. *P≤0.05, **P≤0.01, ***P≤0.001, ****P≤0.0001. See also **Tables S2, S4, and S5**.

**Figure S5.**
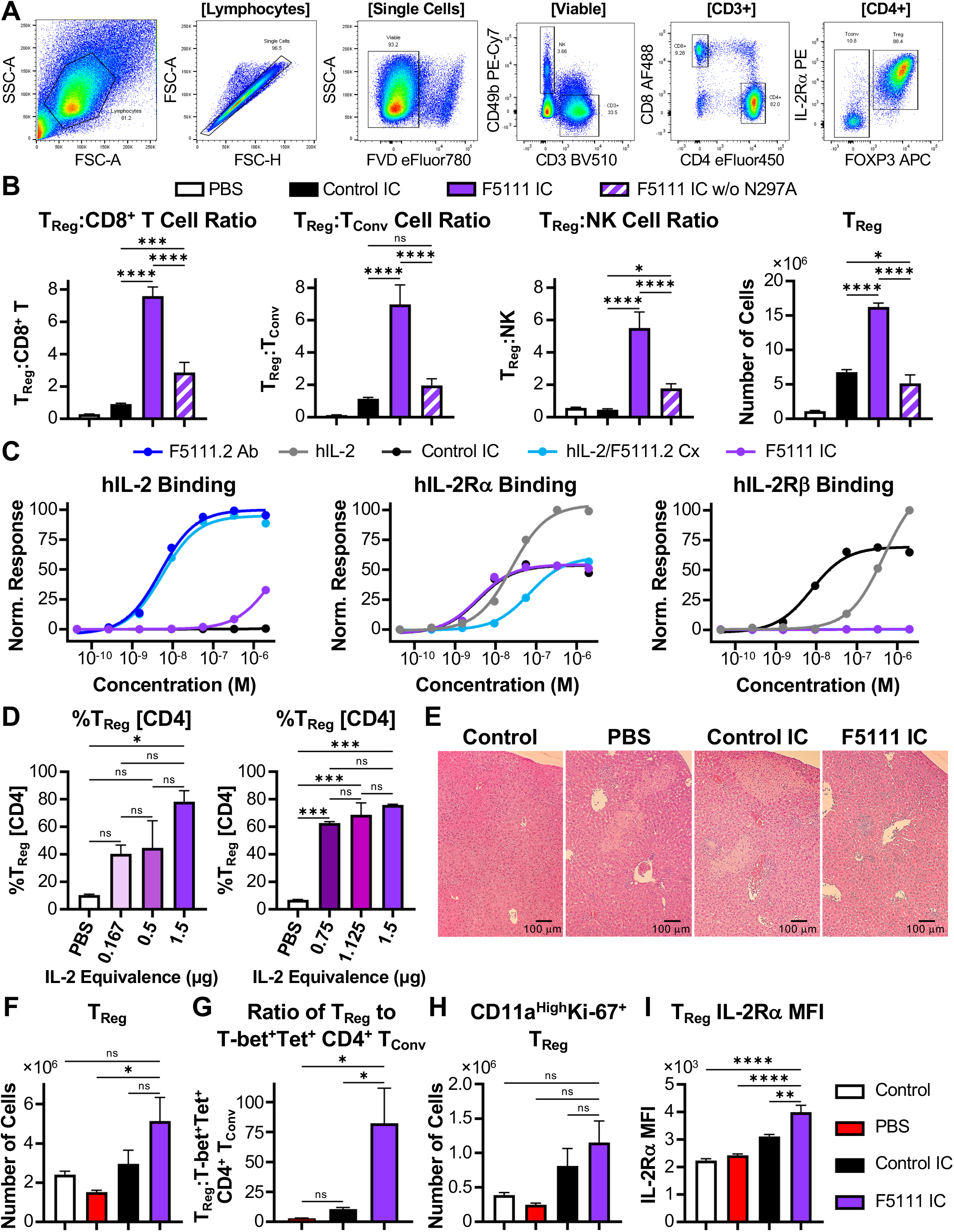
*In* vivo characterization of F5111 IC. **(A)** Representative flow cytometry plots illustrating the gating strategy used for NOD mouse immune cell subset expansion studies. **(B)** Ratios of T_Reg_ to CD8^+^ T cells (left), T_Reg_ to T_Conv_ cells (middle), and T_Reg_ to NK cells (right) in spleens harvested from NOD mice (n=4 per group) treated daily for four days (*i.p.*) with either PBS or 8.2 μg (1.5 μg IL-2 equivalence) Control IC, F5111 IC, or F5111 IC without the N297A mutation (w/o N297A, intact effector function). Data represent mean + SD. Statistical significance was determined by one-way ANOVA with a Tukey post hoc test. Significance is shown between Control IC and F5111 IC with and without the N297A mutation. **(C)** Equilibrium biolayer interferometry-based titrations of F5111.2 antibody (Ab), hIL-2, Control IC, hIL-2/F5111.2 complex (Cx, 1:1 molar ratio), and F5111 IC binding to immobilized hIL-2 (left), immobilized hIL-2Rα (middle), and immobilized hIL-2Rβ (right). **(D)** C57BL/6 mice (n=2 per group) were treated daily for 4 days (*i.p.*) with either PBS or varying dosages of F5111 IC: 0.91 μg (0.167 μg IL-2 equivalence); 2.7 μg (0.5 μg IL-2 equivalence); 4.1 μg (0.75 μg IL-2 equivalence); 6.2 μg (1.125 μg IL-2 equivalence); or 8.2 μg (1.5 μg IL-2 equivalence). Spleens were harvested 24 hours after the last dose. Percent of T_Reg_s within the CD4^+^ T cell population is shown. Data represent mean + SD. Statistical significance was determined by one-way ANOVA with a Tukey post hoc test. **(E-I)** C57BL/6 mice were administered (*i.p.*) 25 cysts of the ME-49 strain of *Toxoplasma gondii* (*T. gondii*) on day 0. Control group designates disease-free mice that were not given cysts. Starting on day 1, mice were treated daily for 5 days (*i.p.*) with either PBS (Control, n=5; PBS, n=4) or 8.2 μg (1.5 μg IL-2 equivalence) Control IC (n=5) or F5111 IC (n=5). Mice were sacrificed on day 10. **(E)** Representative H&E staining of harvested mouse livers. **(F)** Total number of T_Reg_s in harvested spleen. **(G)** Ratio of T_Reg_s to T-bet^+^Tetramer (Tet)^+^ CD4^+^ T_Conv_ cells in harvested mouse spleen. **(H)** Total number of CD11a^High^Ki-67^+^ T_Reg_s in harvested mouse spleen. **(I)** IL-2Rα MFI of T_Reg_s in harvested mouse spleen. Data in **(F-I)** represent mean + SEM. Statistical significance in **(F-I)** was determined by one-way ANOVA with a Tukey post hoc test. Significance compared to F5111 IC is shown. All statistical data are provided in **Table S6**. *P≤0.05, **P≤0.01, ***P≤0.001, ****P≤0.0001. See also **Table S2**.

**Figure S6.**
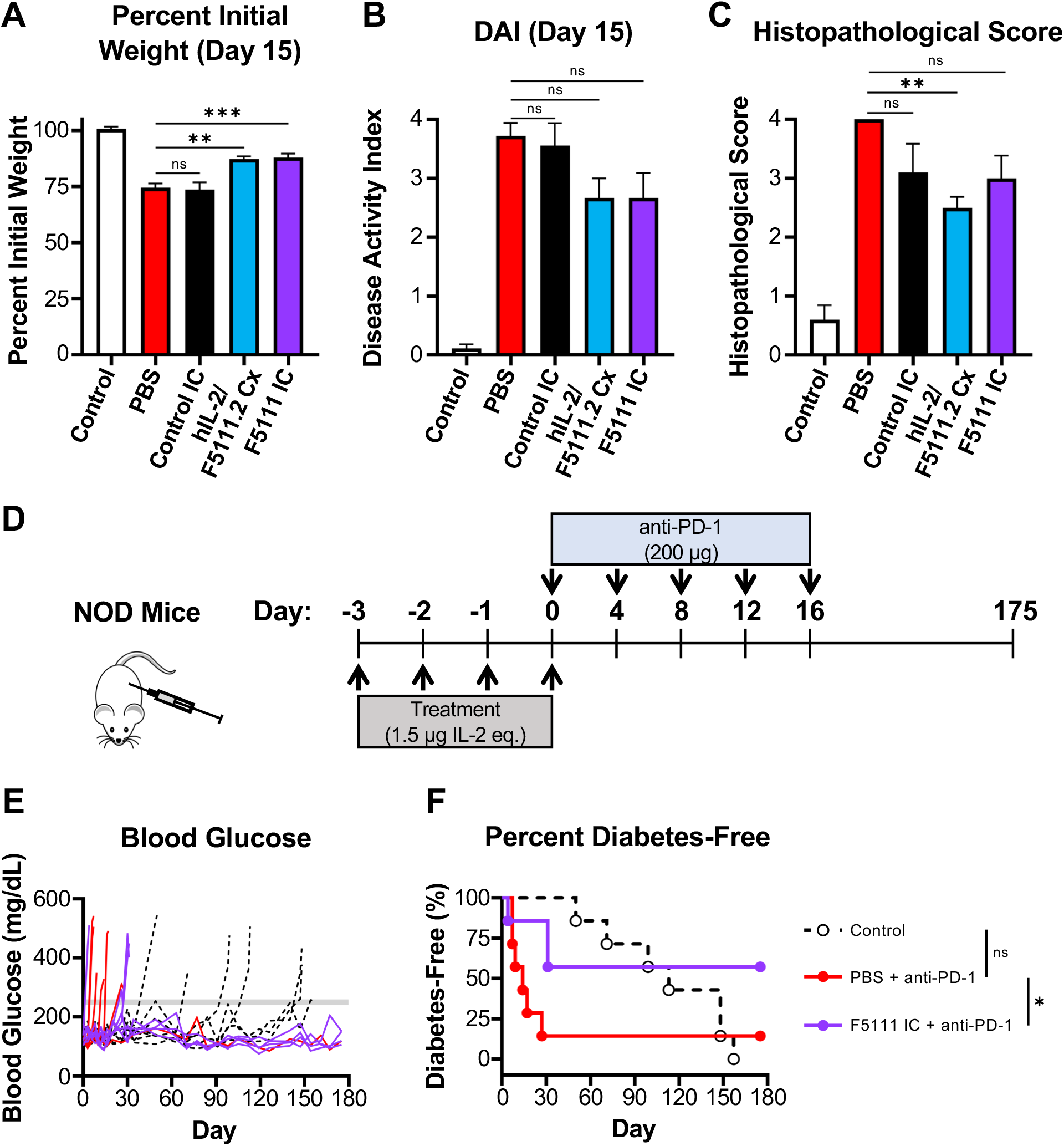
Evaluation of F5111 IC in mouse models of autoimmune disease. **(A-C)** BALB/c mice (n=6 per group) were treated daily for 7 days (*i.p.*) with either PBS (Control and PBS), 1.5 μg hIL-2 complexed with 6.6 μg F5111.2 antibody (1:2 molar ratio, hIL-2/F5111.2 Cx), or 8.2 μg (1.5 μg IL-2 equivalence) Control IC or F5111 IC. Beginning on day 7, all groups except for the disease-free cohort (Control) were administered 3% DSS in their drinking water. Mice were sacrificed on day 15. **(A)** Weight change on day 15. **(B)** Disease activity index (DAI) on day 15. **(C)** Histopathology scores for H&E stained colons (n=5 Control, Control IC; n=6 PBS, Cx, F5111 IC). Data represent mean ± SEM. Statistical significance was determined by one-way ANOVA with a Tukey post hoc test. All plots show significance of Control IC, Cx, and F5111 IC treated mice versus PBS treated mice. **(D-F)** 8-week-old NOD mice (n=7 per group) were treated daily for 4 days (*i.p.*, days -3, -2, -1, 0) with either PBS (Control and PBS) or 8.2 μg (1.5 μg IL-2 equivalence) F5111 IC. Starting on day 0 (4 hours after the last IC dose), mice were administered anti-PD-1 antibody (200 μg) every 4 days until day 16. Control group designates mice that did not receive anti-PD-1 antibody. **(E)** Blood glucose concentrations over the study. The threshold 250 mg/dL value is indicated by the gray line. **(F)** Percent diabetes-free mice. Statistical significance was determined by pairwise comparisons using the Log-rank (Mantel-Cox) test. Statistical significance compared to mice treated with PBS + anti-PD-1 is shown. All statistical data are provided in **Table S6**. *P≤0.05, **P≤0.01, ***P≤0.001, ****P≤0.0001.

**Table S1.**
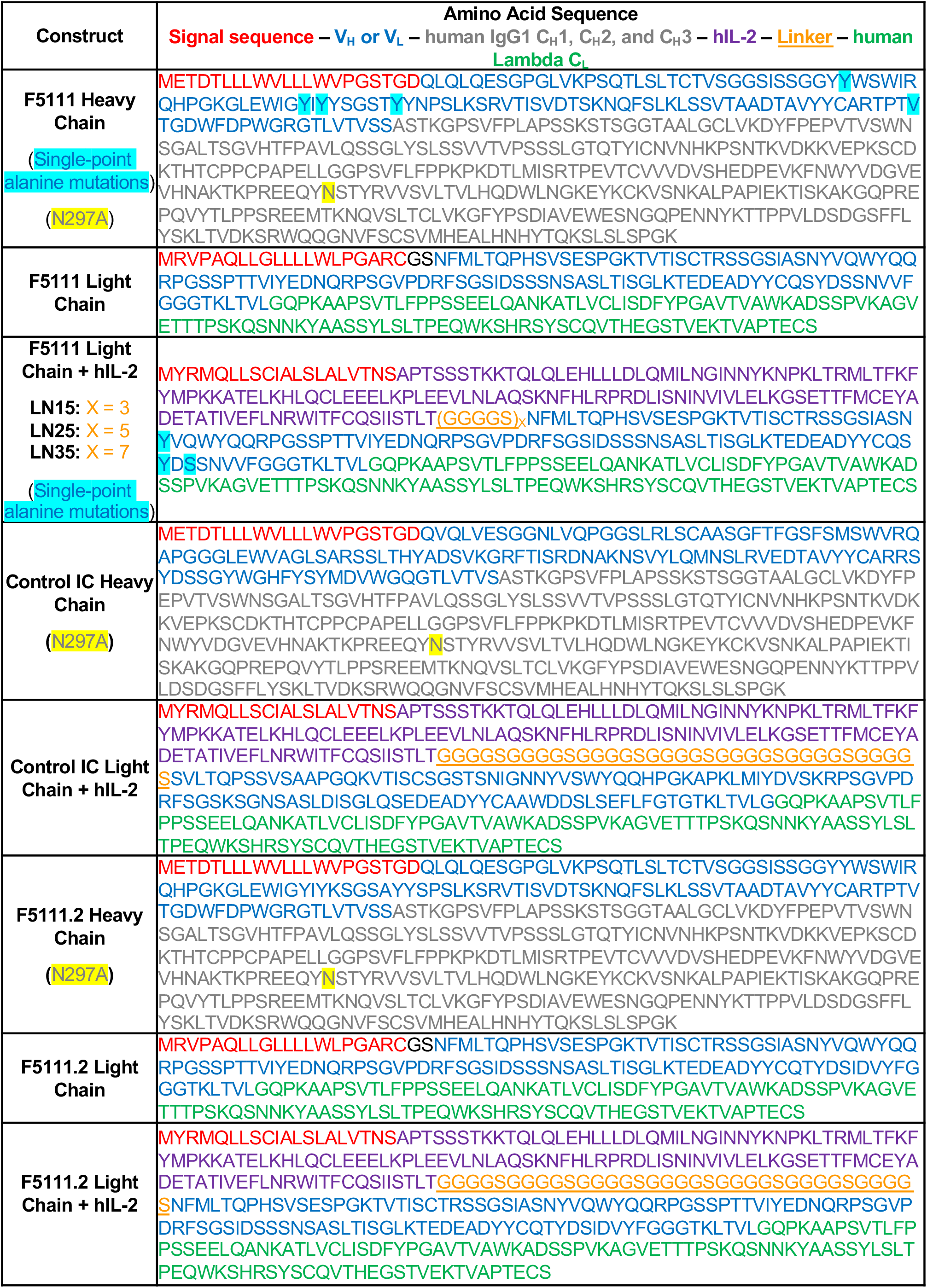
Antibody and IC sequences.

**Table S2.** Equilibrium K_D_ values from biolayer interferometry studies.

| Treatment | Equilibrium $K_D$ (nM) | | | Figure |
| --- | --- | --- | --- | --- |
| | hIL-2 | hIL-2Rα | hIL-2Rβ | |
| F5111 Ab | 2.7 | - | - | 2A, S1C, S1D |
| hIL-2 | - | 16 | 480 | 2A, S1C, S1D |
| Control IC | - | 3.4 | 4.4 | 2A |
| hIL-2/F5111 Complex | 4.8 | 18 | >2000 | 2A |
| F5111 IC LN15 | >2000 | 7.5 | 210 | S1C |
| F5111 IC LN25 | >2000 | 3.4 | >2000 | S1D |
| F5111 IC LN35 | >2000 | 3.0 | >2000 | 2A |
| hIL-2 | - | 16 | 310 | S1E |
| F5111 Ab | 2.1 | - | - | S1E |
| F5111 IC LN25 P1 | >2000 | 11 | >700 | S1E |
| F5111 IC LN25 P2 | >2000 | 8.1 | >1000 | S1E |
| F5111 IC LN25 P3 | >2000 | 4.8 | >2000 | S1E |
| hIL-2 | - | 25 | 470 | 3B, S3B, S5C |
| Control IC | - | 2.5 | 4.1 | 3B, S3B |
| F5111 IC | >2000 | 3.2 | >2000 | 3B, S3B, S4A |
| Y33A IC | >2000 | 2.6 | >2000 | 3B, S3B |
| Y94A IC | >2000 | 2.2 | >2000 | 3B, S3B |
| S96A IC | >2000 | 2.5 | >2000 | 3B, S3B |
| Y35A IC | >2000 | 2.5 | >2000 | 3B, S3B |
| Y52A IC | >2000 | 2.8 | 500 | 3B, S3B |
| Y54A IC | >2000 | 2.8 | >2000 | 3B, S3B |
| Y60A IC | >2000 | 2.7 | >2000 | 3B, S3B |
| V103A IC | >2000 | 2.4 | >2000 | 3B, S3B |
| F5111.2 IC | >2000 | 2.4 | >2000 | 3B, S3B |
| Control IC + N297A | - | 3.8 | 7.7 | S4A, S5C |
| F5111 IC + N297A | >2000 | 3.6 | >2000 | S4A, S5C |
| F5111.2 Ab + N297A | 4.9 | - | - | S5C |
| hIL-2/F5111.2 Complex | 5.6 | 68 | >2000 | S5C |

**Table S3.**
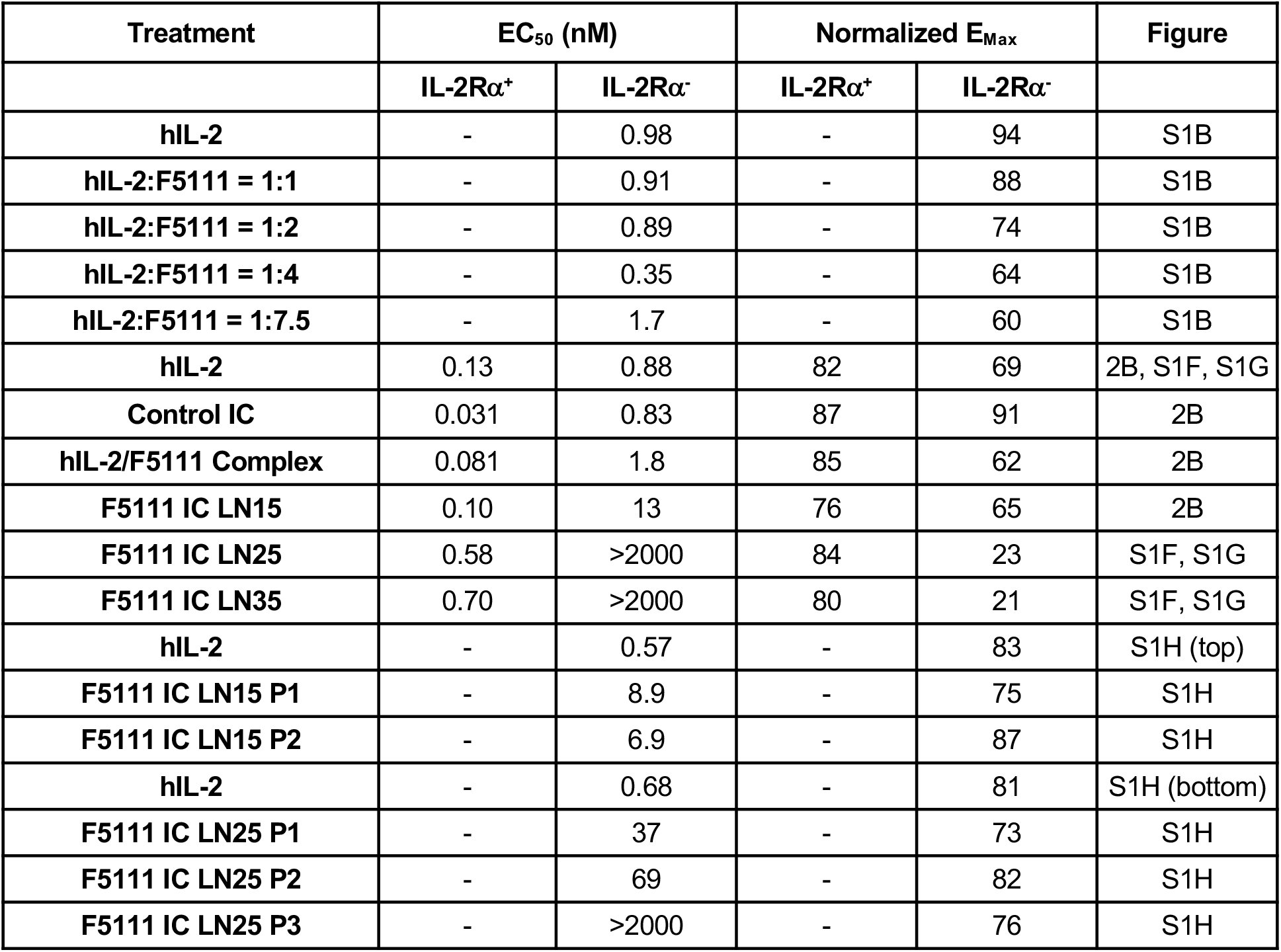
EC_50_ and E_Max_ values from YT-1 cell activation studies.

| Treatment | EC<sub>50</sub> (nM) |  | Normalized E<sub>Max</sub> |  | Figure |
| --- | --- | --- | --- | --- | --- |
| | IL-2R$\alpha^+$ | IL-2R$\alpha^-$ | IL-2R$\alpha^+$ | IL-2R$\alpha^-$ | |
| hIL-2 | - | 0.98 | - | 94 | S1B |
| hIL-2:F5111 = 1:1 | - | 0.91 | - | 88 | S1B |
| hIL-2:F5111 = 1:2 | - | 0.89 | - | 74 | S1B |
| hIL-2:F5111 = 1:4 | - | 0.35 | - | 64 | S1B |
| hIL-2:F5111 = 1:7.5 | - | 1.7 | - | 60 | S1B |
| hIL-2 | 0.13 | 0.88 | 82 | 69 | 2B, S1F, S1G |
| Control IC | 0.031 | 0.83 | 87 | 91 | 2B |
| hIL-2/F5111 Complex | 0.081 | 1.8 | 85 | 62 | 2B |
| F5111 IC LN15 | 0.10 | 13 | 76 | 65 | 2B |
| F5111 IC LN25 | 0.58 | >2000 | 84 | 23 | S1F, S1G |
| F5111 IC LN35 | 0.70 | >2000 | 80 | 21 | S1F, S1G |
| hIL-2 | - | 0.57 | - | 83 | S1H (top) |
| F5111 IC LN15 P1 | - | 8.9 | - | 75 | S1H |
| F5111 IC LN15 P2 | - | 6.9 | - | 87 | S1H |
| hIL-2 | - | 0.68 | - | 81 | S1H (bottom) |
| F5111 IC LN25 P1 | - | 37 | - | 73 | S1H |
| F5111 IC LN25 P2 | - | 69 | - | 82 | S1H |
| F5111 IC LN25 P3 | - | >2000 | - | 76 | S1H |

**Table S4.**
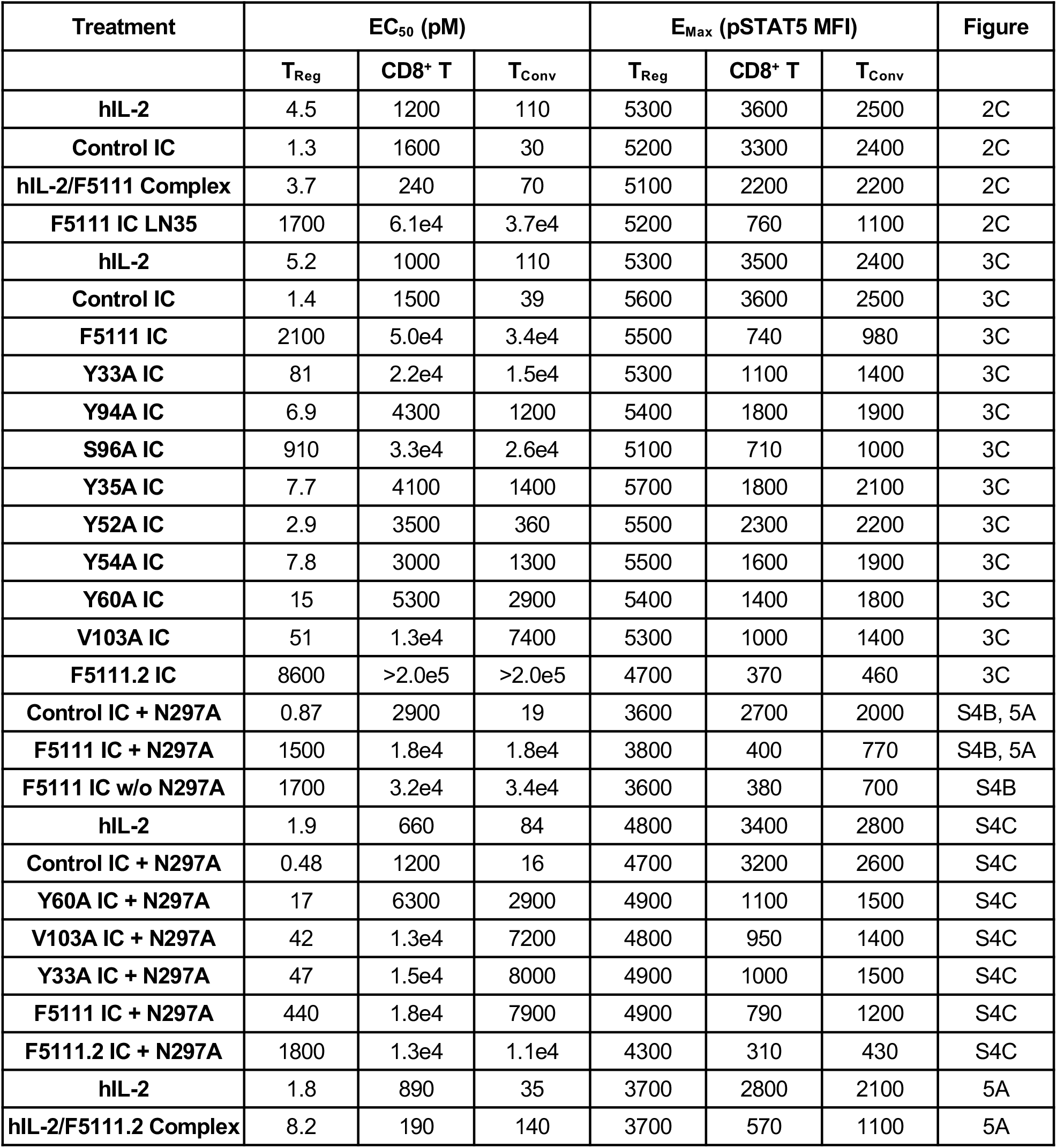
EC_50_ and E_Max_ values from human PBMC activation studies.

| Treatment | EC<sub>50</sub> (pM) |  |  | E<sub>Max</sub> (pSTAT5 MFI) |  |  | Figure |
| --- | --- | --- | --- | --- | --- | --- | --- |
|  | T<sub>Reg</sub> | CD8<sup>+</sup> T | T<sub>Conv</sub> | T<sub>Reg</sub> | CD8<sup>+</sup> T | T<sub>Conv</sub> |  |
| hIL-2 | 4.5 | 1200 | 110 | 5300 | 3600 | 2500 | 2C |
| Control IC | 1.3 | 1600 | 30 | 5200 | 3300 | 2400 | 2C |
| hIL-2/F5111 Complex | 3.7 | 240 | 70 | 5100 | 2200 | 2200 | 2C |
| F5111 IC LN35 | 1700 | 6.1e4 | 3.7e4 | 5200 | 760 | 1100 | 2C |
| hIL-2 | 5.2 | 1000 | 110 | 5300 | 3500 | 2400 | 3C |
| Control IC | 1.4 | 1500 | 39 | 5600 | 3600 | 2500 | 3C |
| F5111 IC | 2100 | 5.0e4 | 3.4e4 | 5500 | 740 | 980 | 3C |
| Y33A IC | 81 | 2.2e4 | 1.5e4 | 5300 | 1100 | 1400 | 3C |
| Y94A IC | 6.9 | 4300 | 1200 | 5400 | 1800 | 1900 | 3C |
| S96A IC | 910 | 3.3e4 | 2.6e4 | 5100 | 710 | 1000 | 3C |
| Y35A IC | 7.7 | 4100 | 1400 | 5700 | 1800 | 2100 | 3C |
| Y52A IC | 2.9 | 3500 | 360 | 5500 | 2300 | 2200 | 3C |
| Y54A IC | 7.8 | 3000 | 1300 | 5500 | 1600 | 1900 | 3C |
| Y60A IC | 15 | 5300 | 2900 | 5400 | 1400 | 1800 | 3C |
| V103A IC | 51 | 1.3e4 | 7400 | 5300 | 1000 | 1400 | 3C |
| F5111.2 IC | 8600 | >2.0e5 | >2.0e5 | 4700 | 370 | 460 | 3C |
| Control IC + N297A | 0.87 | 2900 | 19 | 3600 | 2700 | 2000 | S4B, 5A |
| F5111 IC + N297A | 1500 | 1.8e4 | 1.8e4 | 3800 | 400 | 770 | S4B, 5A |
| F5111 IC w/o N297A | 1700 | 3.2e4 | 3.4e4 | 3600 | 380 | 700 | S4B |
| hIL-2 | 1.9 | 660 | 84 | 4800 | 3400 | 2800 | S4C |
| Control IC + N297A | 0.48 | 1200 | 16 | 4700 | 3200 | 2600 | S4C |
| Y60A IC + N297A | 17 | 6300 | 2900 | 4900 | 1100 | 1500 | S4C |
| V103A IC + N297A | 42 | 1.3e4 | 7200 | 4800 | 950 | 1400 | S4C |
| Y33A IC + N297A | 47 | 1.5e4 | 8000 | 4900 | 1000 | 1500 | S4C |
| F5111 IC + N297A | 440 | 1.8e4 | 7900 | 4900 | 790 | 1200 | S4C |
| F5111.2 IC + N297A | 1800 | 1.3e4 | 1.1e4 | 4300 | 310 | 430 | S4C |
| hIL-2 | 1.8 | 890 | 35 | 3700 | 2800 | 2100 | 5A |
| hIL-2/F5111.2 Complex | 8.2 | 190 | 140 | 3700 | 570 | 1100 | 5A |

**Table S5.** EC_50_ and E_Max_ values from mouse splenocyte activation studies.

| Treatment | EC<sub>50</sub> (pM) |  |  | E<sub>Max</sub> (pSTAT5 MFI) |  |  | Figure |
| --- | --- | --- | --- | --- | --- | --- | --- |
|  | T<sub>Reg</sub> | CD8<sup>+</sup> T | T<sub>Conv</sub> | T<sub>Reg</sub> | CD8<sup>+</sup> T | T<sub>Conv</sub> |  |
| hIL-2 | 6.8 | 1.3e5 | - | 8600 | 5700 | 1000 | S4D |
| Control IC + N297A | 3.2 | 2.9e5 | - | 8600 | 4900 | 1100 | S4D |
| Y60A IC + N297A | 93 | >2e6 | - | 8400 | 280 | 650 | S4D |
| V103A IC + N297A | 260 | >2e6 | - | 8000 | 370 | 720 | S4D |
| Y33A IC + N297A | 270 | >2e6 | - | 8100 | 340 | 680 | S4D |
| F5111 IC + N297A | 2100 | >2e6 | - | 7700 | 320 | 720 | S4D |
| F5111.2 IC + N297A | 6300 | >2e6 | - | 5500 | 220 | 660 | S4D |

**Table S6.**
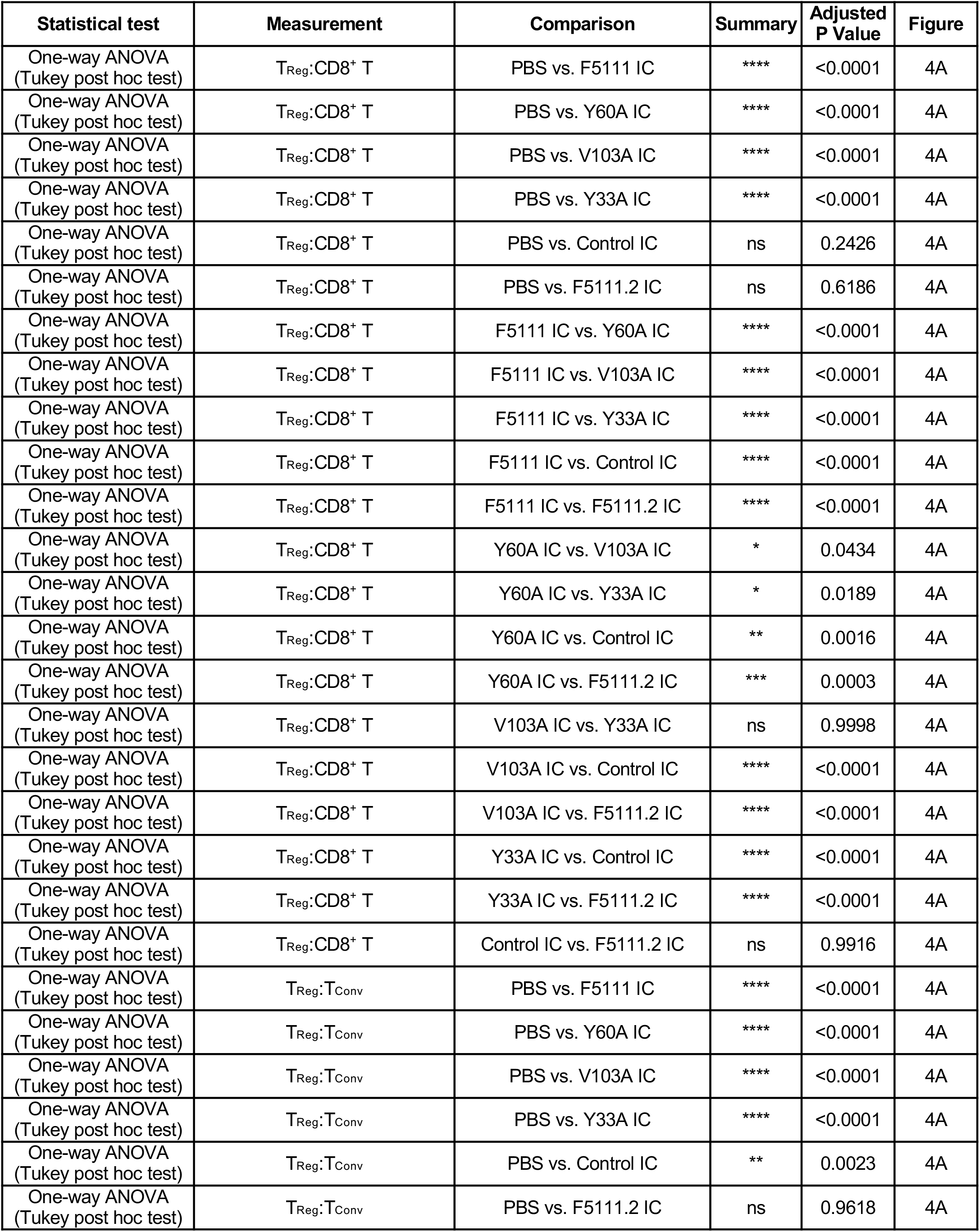
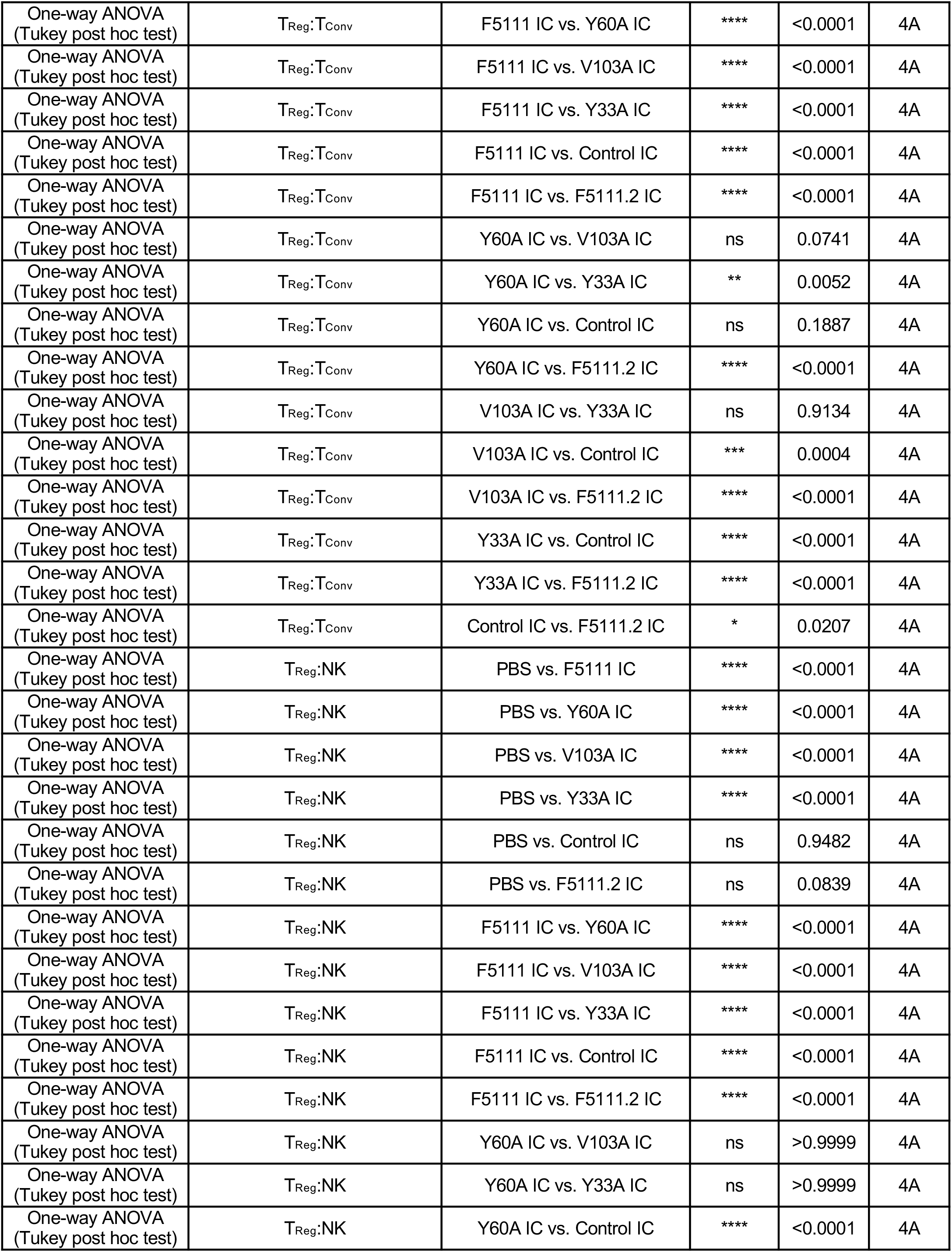
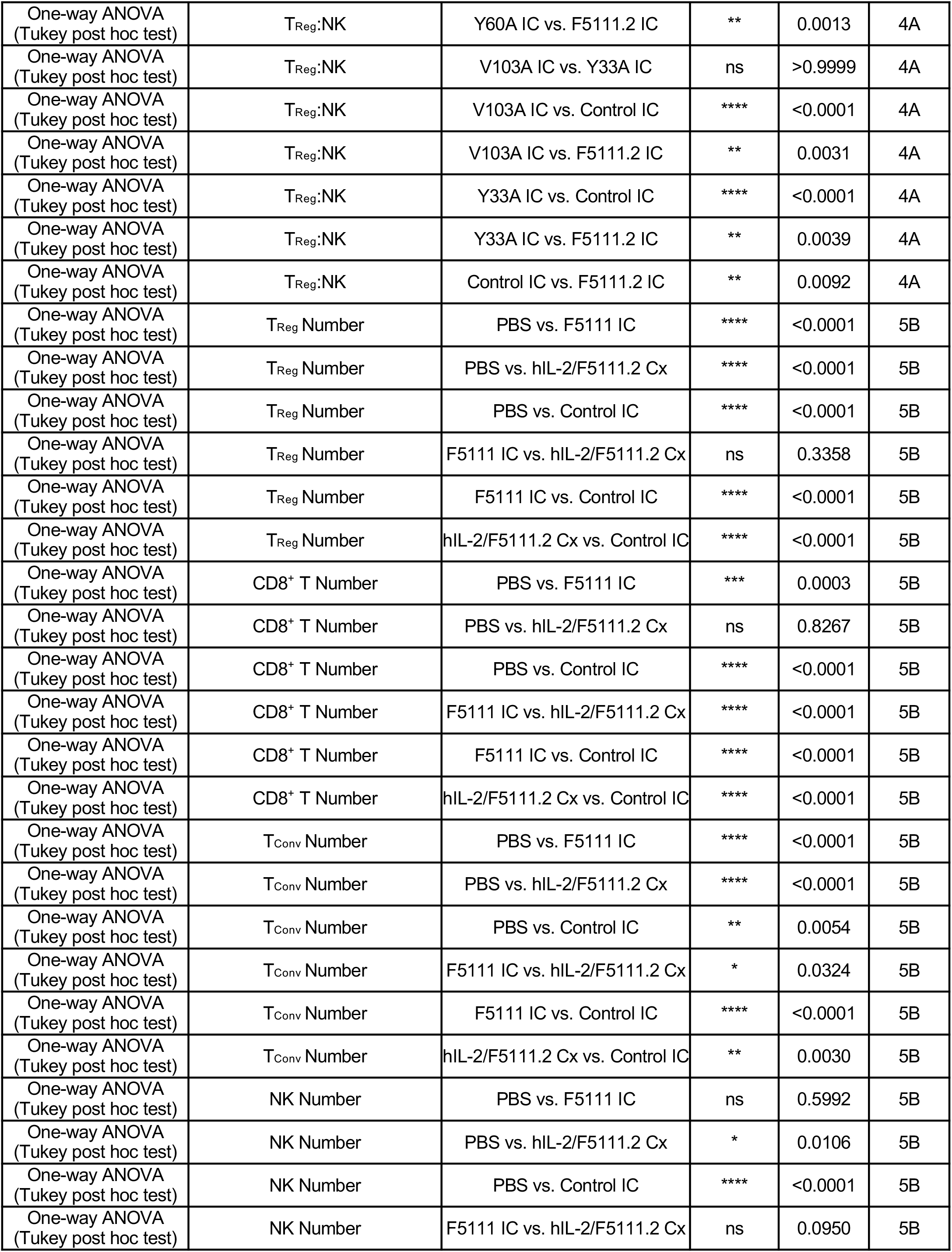
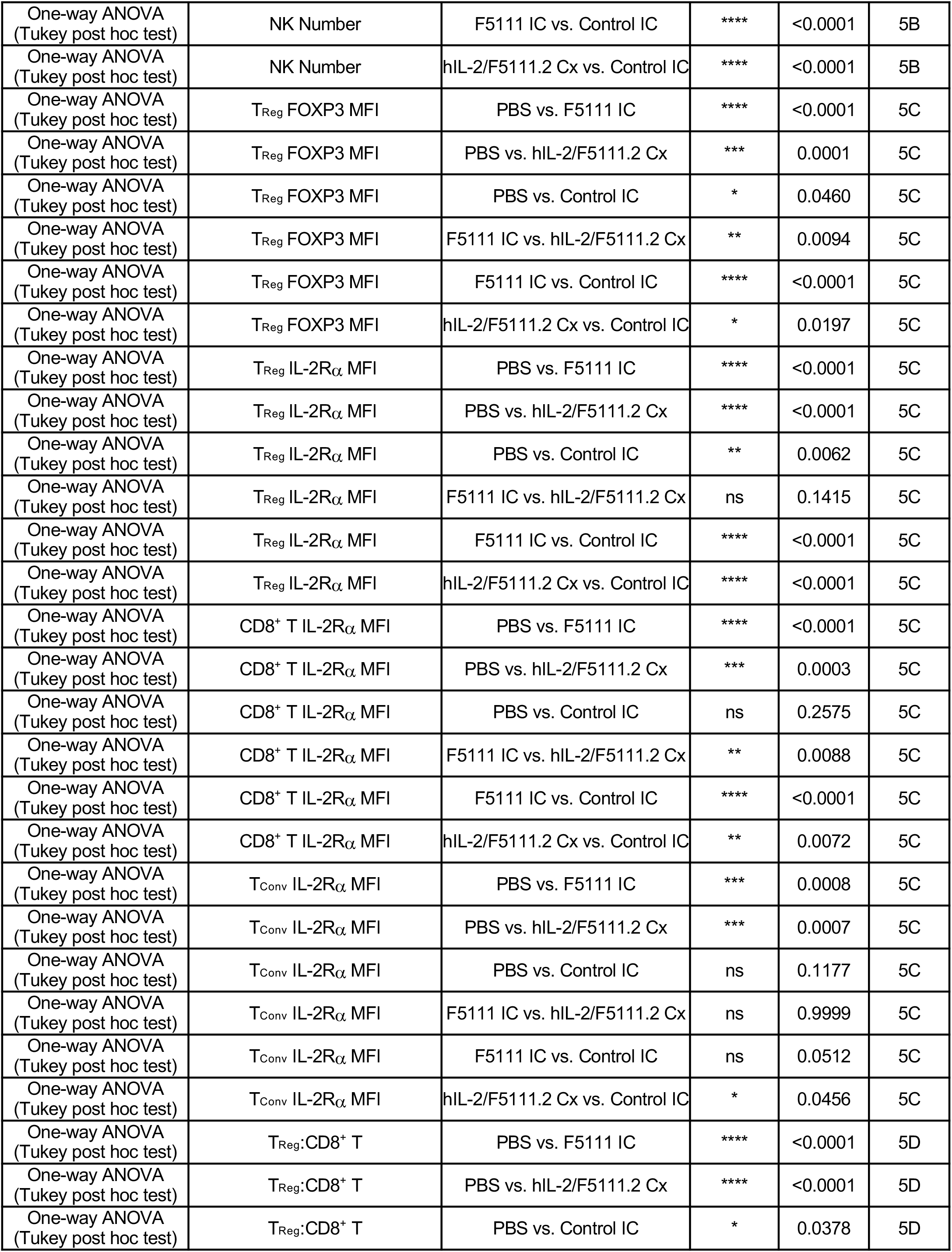
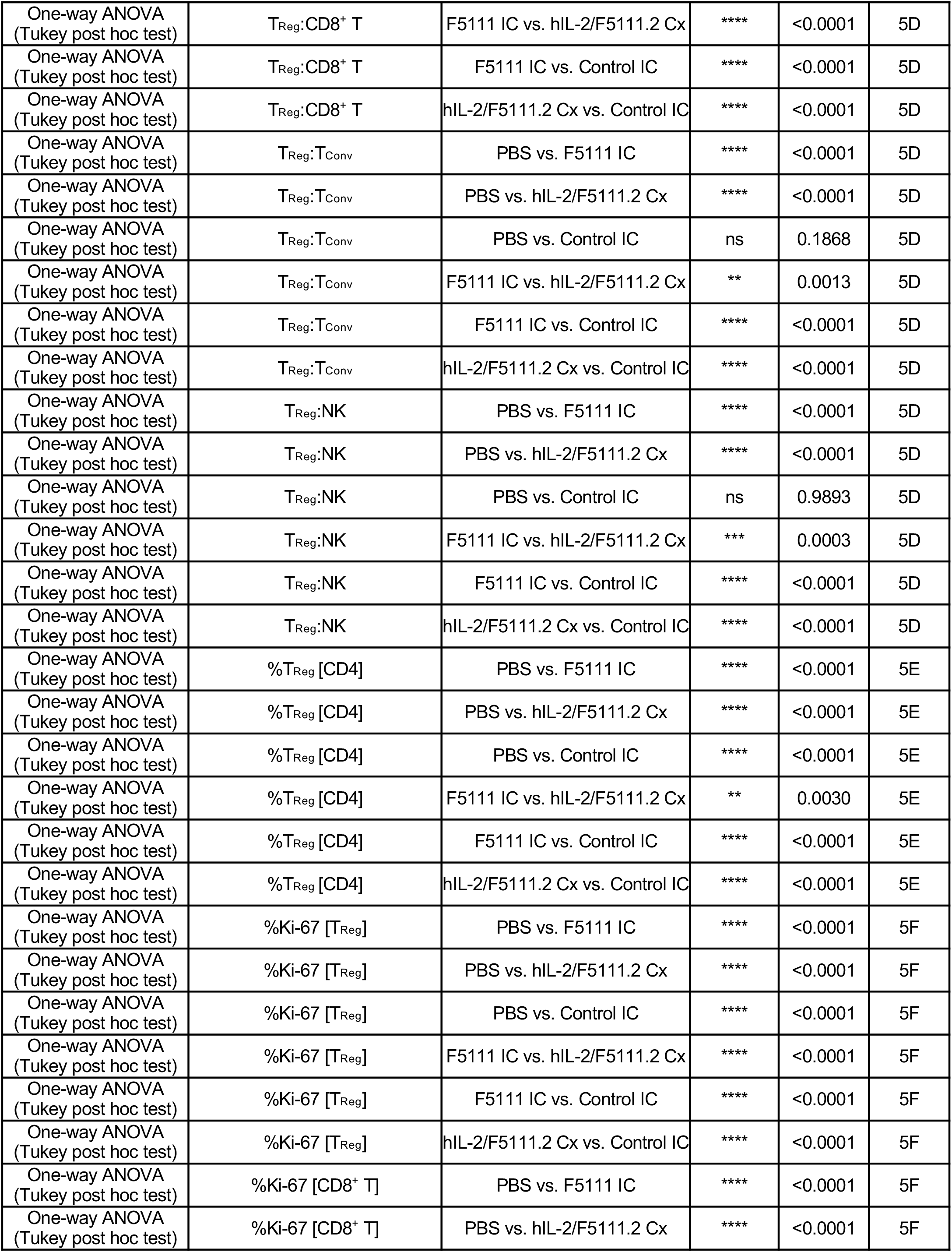
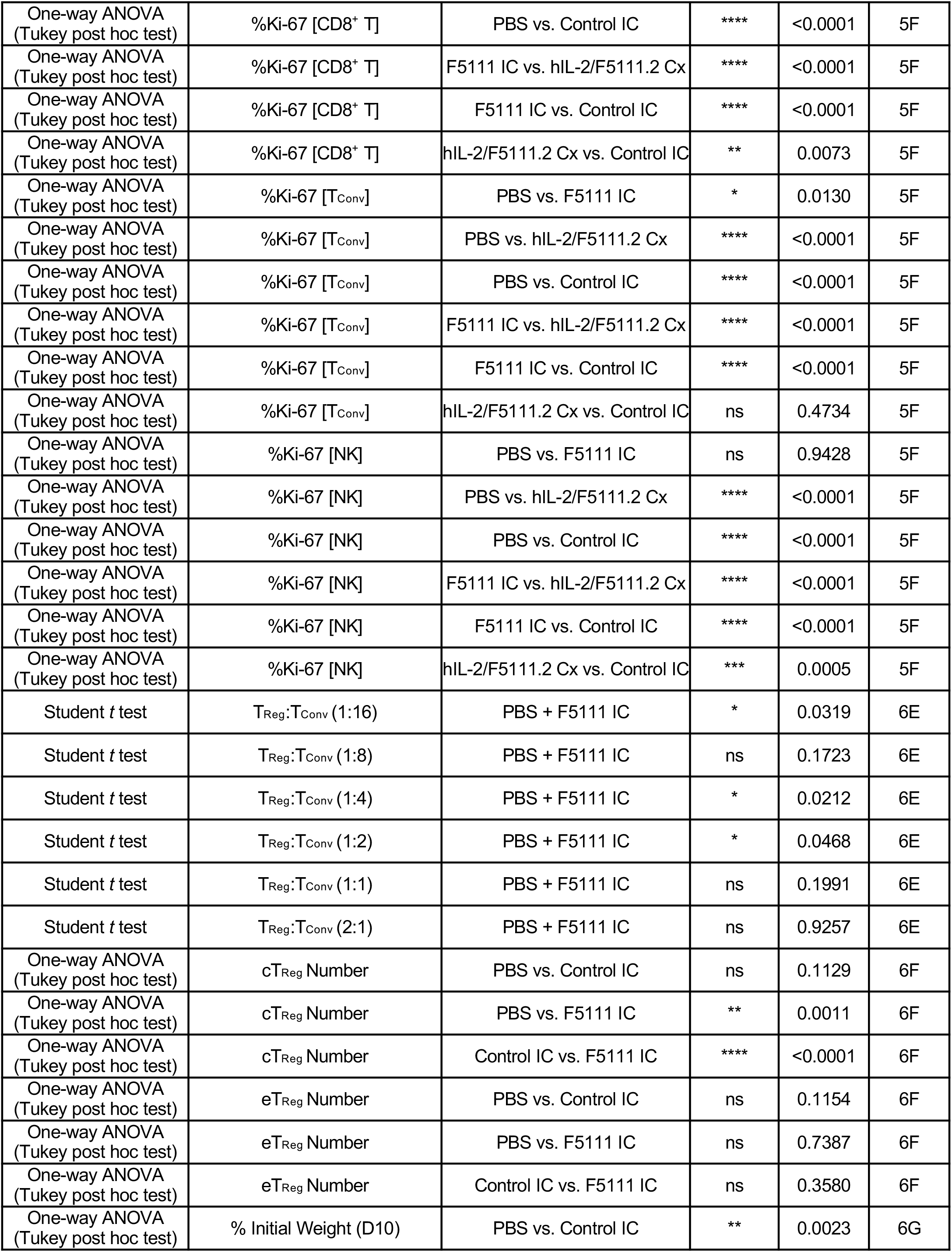
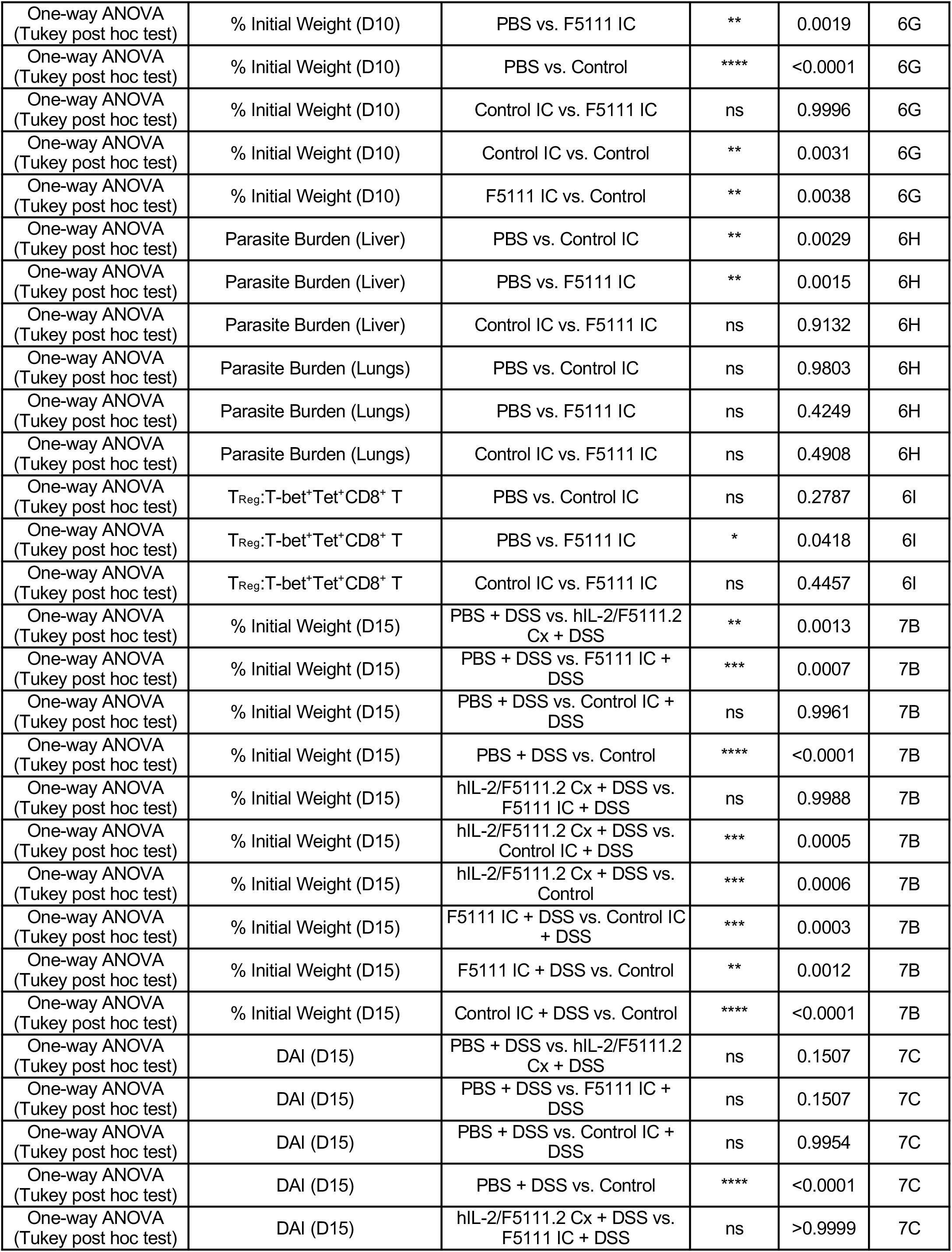
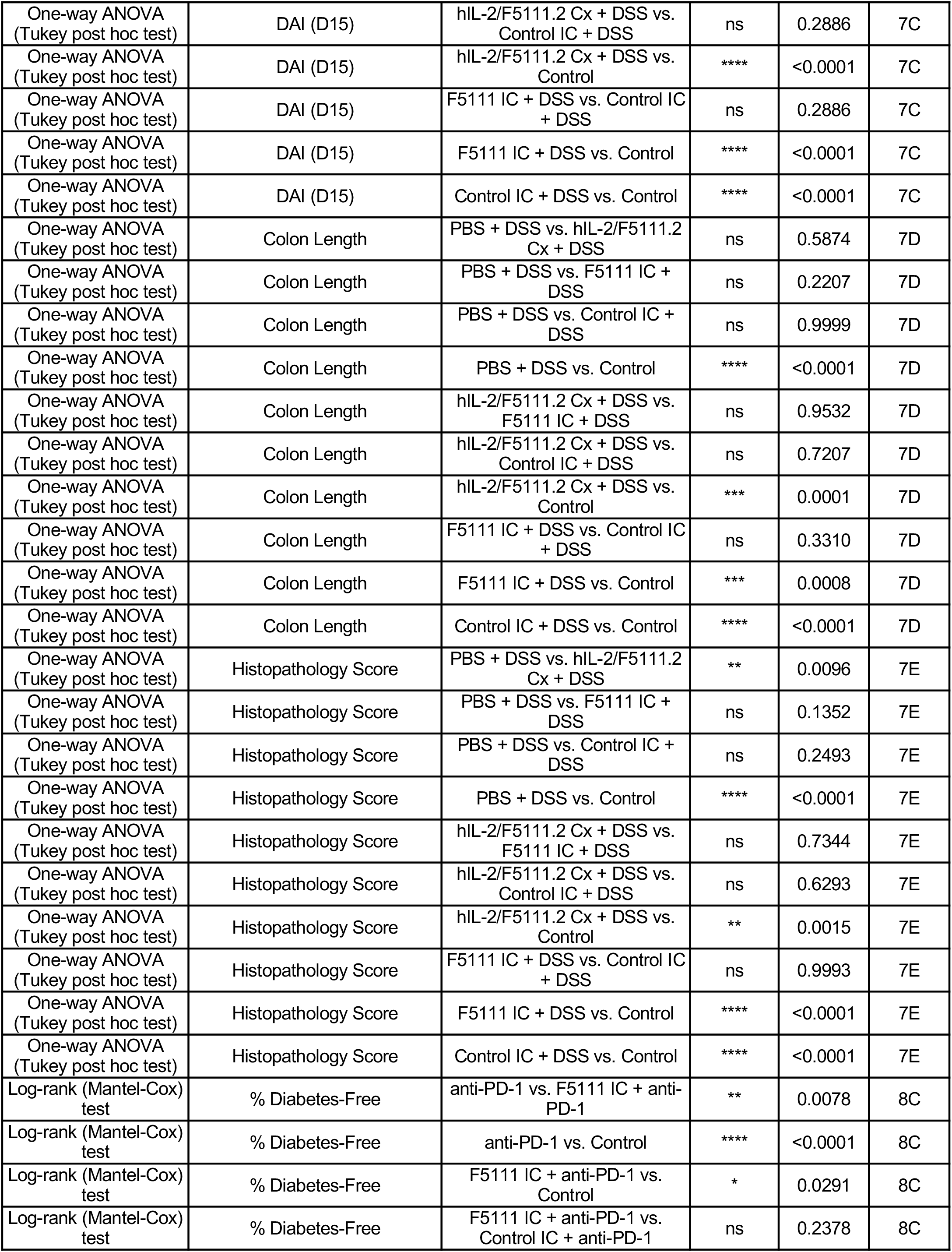
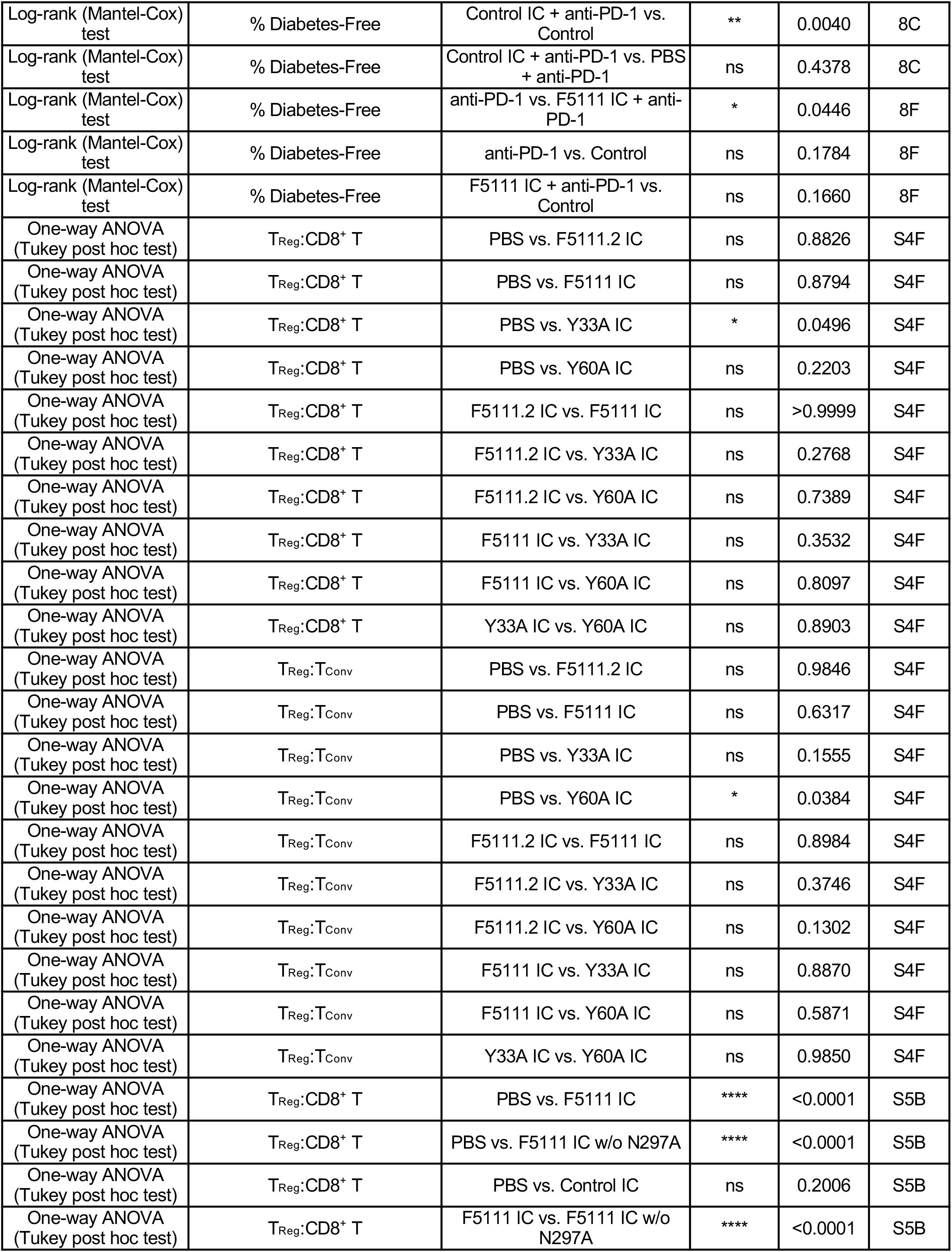
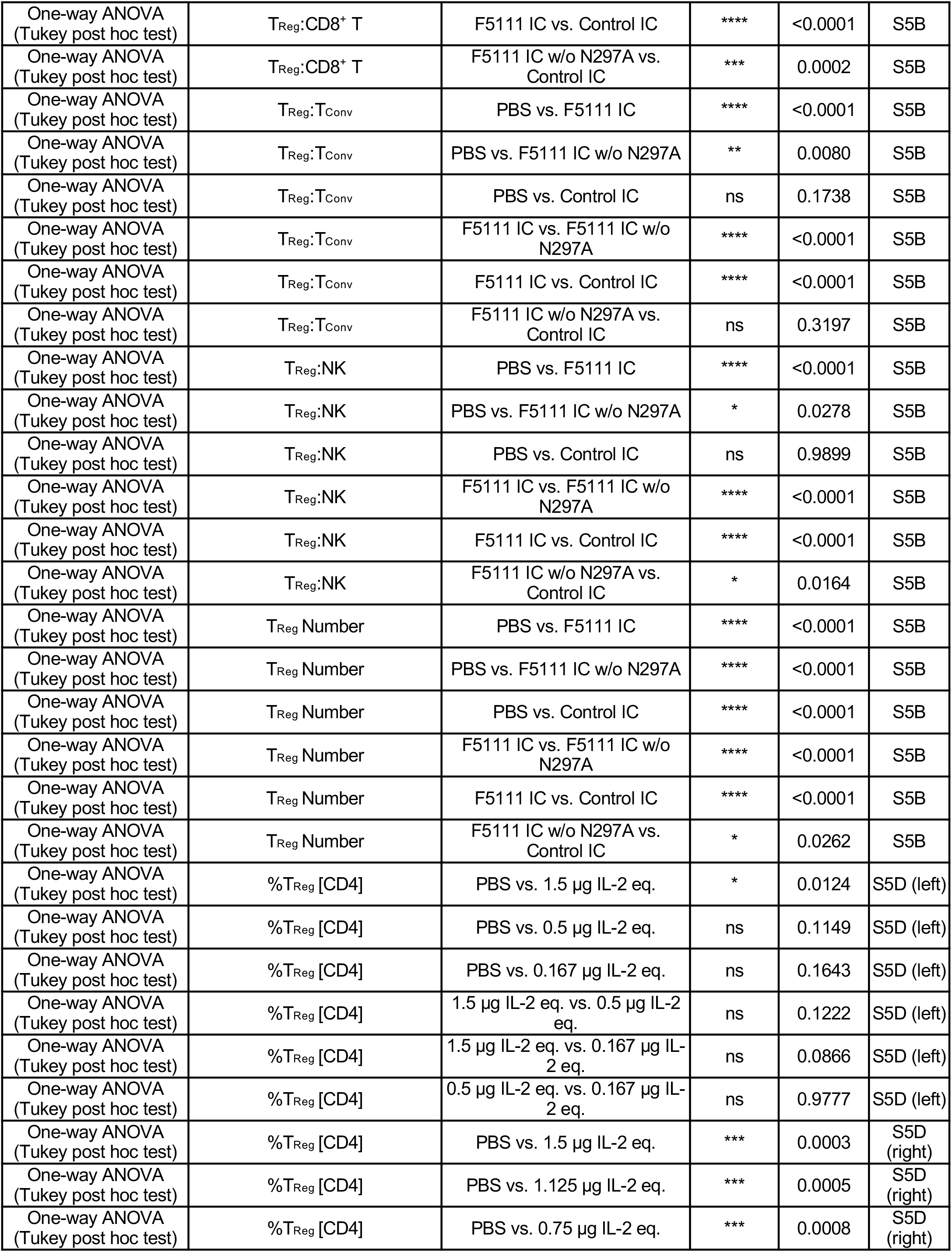
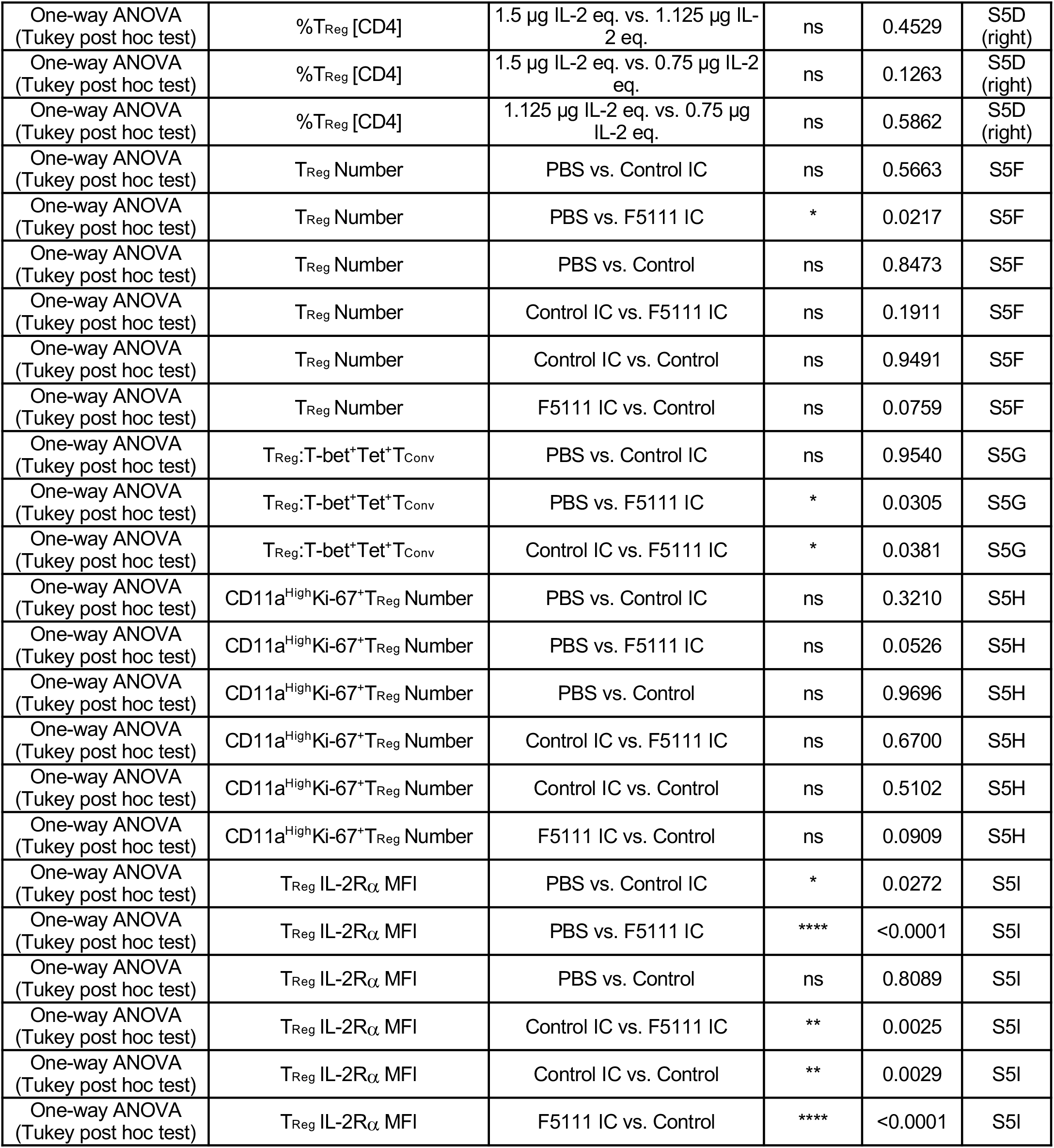
Statistical analysis.

| Statistical test | Measurement | Comparison | Summary | Adjusted P Value | Figure |
| --- | --- | --- | --- | --- | --- |
| One-way ANOVA<br>(Tukey post hoc test) | T<sub>Reg</sub>:CD8<sup>+</sup> T | PBS vs. F5111 IC | **** | <0.0001 | 4A |
| One-way ANOVA<br>(Tukey post hoc test) | T<sub>Reg</sub>:CD8<sup>+</sup> T | PBS vs. Y60A IC | **** | <0.0001 | 4A |
| One-way ANOVA<br>(Tukey post hoc test) | T<sub>Reg</sub>:CD8<sup>+</sup> T | PBS vs. V103A IC | **** | <0.0001 | 4A |
| One-way ANOVA<br>(Tukey post hoc test) | T<sub>Reg</sub>:CD8<sup>+</sup> T | PBS vs. Y33A IC | **** | <0.0001 | 4A |
| One-way ANOVA<br>(Tukey post hoc test) | T<sub>Reg</sub>:CD8<sup>+</sup> T | PBS vs. Control IC | ns | 0.2426 | 4A |
| One-way ANOVA<br>(Tukey post hoc test) | T<sub>Reg</sub>:CD8<sup>+</sup> T | PBS vs. F5111.2 IC | ns | 0.6186 | 4A |
| One-way ANOVA<br>(Tukey post hoc test) | T<sub>Reg</sub>:CD8<sup>+</sup> T | F5111 IC vs. Y60A IC | **** | <0.0001 | 4A |
| One-way ANOVA<br>(Tukey post hoc test) | T<sub>Reg</sub>:CD8<sup>+</sup> T | F5111 IC vs. V103A IC | **** | <0.0001 | 4A |
| One-way ANOVA<br>(Tukey post hoc test) | T<sub>Reg</sub>:CD8<sup>+</sup> T | F5111 IC vs. Y33A IC | **** | <0.0001 | 4A |
| One-way ANOVA<br>(Tukey post hoc test) | T<sub>Reg</sub>:CD8<sup>+</sup> T | F5111 IC vs. Control IC | **** | <0.0001 | 4A |
| One-way ANOVA<br>(Tukey post hoc test) | T<sub>Reg</sub>:CD8<sup>+</sup> T | F5111 IC vs. F5111.2 IC | **** | <0.0001 | 4A |
| One-way ANOVA<br>(Tukey post hoc test) | T<sub>Reg</sub>:CD8<sup>+</sup> T | Y60A IC vs. V103A IC | * | 0.0434 | 4A |
| One-way ANOVA<br>(Tukey post hoc test) | T<sub>Reg</sub>:CD8<sup>+</sup> T | Y60A IC vs. Y33A IC | * | 0.0189 | 4A |
| One-way ANOVA<br>(Tukey post hoc test) | T<sub>Reg</sub>:CD8<sup>+</sup> T | Y60A IC vs. Control IC | ** | 0.0016 | 4A |
| One-way ANOVA<br>(Tukey post hoc test) | T<sub>Reg</sub>:CD8<sup>+</sup> T | Y60A IC vs. F5111.2 IC | *** | 0.0003 | 4A |
| One-way ANOVA<br>(Tukey post hoc test) | T<sub>Reg</sub>:CD8<sup>+</sup> T | V103A IC vs. Y33A IC | ns | 0.9998 | 4A |
| One-way ANOVA<br>(Tukey post hoc test) | T<sub>Reg</sub>:CD8<sup>+</sup> T | V103A IC vs. Control IC | **** | <0.0001 | 4A |
| One-way ANOVA<br>(Tukey post hoc test) | T<sub>Reg</sub>:CD8<sup>+</sup> T | V103A IC vs. F5111.2 IC | **** | <0.0001 | 4A |
| One-way ANOVA<br>(Tukey post hoc test) | T<sub>Reg</sub>:CD8<sup>+</sup> T | Y33A IC vs. Control IC | **** | <0.0001 | 4A |
| One-way ANOVA<br>(Tukey post hoc test) | T<sub>Reg</sub>:CD8<sup>+</sup> T | Y33A IC vs. F5111.2 IC | **** | <0.0001 | 4A |
| One-way ANOVA<br>(Tukey post hoc test) | T<sub>Reg</sub>:CD8<sup>+</sup> T | Control IC vs. F5111.2 IC | ns | 0.9916 | 4A |
| One-way ANOVA<br>(Tukey post hoc test) | T<sub>Reg</sub>:T<sub>Conv</sub> | PBS vs. F5111 IC | **** | <0.0001 | 4A |
| One-way ANOVA<br>(Tukey post hoc test) | T<sub>Reg</sub>:T<sub>Conv</sub> | PBS vs. Y60A IC | **** | <0.0001 | 4A |
| One-way ANOVA<br>(Tukey post hoc test) | T<sub>Reg</sub>:T<sub>Conv</sub> | PBS vs. V103A IC | **** | <0.0001 | 4A |
| One-way ANOVA<br>(Tukey post hoc test) | T<sub>Reg</sub>:T<sub>Conv</sub> | PBS vs. Y33A IC | **** | <0.0001 | 4A |
| One-way ANOVA<br>(Tukey post hoc test) | T<sub>Reg</sub>:T<sub>Conv</sub> | PBS vs. Control IC | ** | 0.0023 | 4A |
| One-way ANOVA<br>(Tukey post hoc test) | T<sub>Reg</sub>:T<sub>Conv</sub> | PBS vs. F5111.2 IC | ns | 0.9618 | 4A |
| One-way ANOVA<br>(Tukey post hoc test) | $T_{Reg}:T_{Conv}$ | F5111 IC vs. Y60A IC | **** | <0.0001 | 4A |
| One-way ANOVA<br>(Tukey post hoc test) | $T_{Reg}:T_{Conv}$ | F5111 IC vs. V103A IC | **** | <0.0001 | 4A |
| One-way ANOVA<br>(Tukey post hoc test) | $T_{Reg}:T_{Conv}$ | F5111 IC vs. Y33A IC | **** | <0.0001 | 4A |
| One-way ANOVA<br>(Tukey post hoc test) | $T_{Reg}:T_{Conv}$ | F5111 IC vs. Control IC | **** | <0.0001 | 4A |
| One-way ANOVA<br>(Tukey post hoc test) | $T_{Reg}:T_{Conv}$ | F5111 IC vs. F5111.2 IC | **** | <0.0001 | 4A |
| One-way ANOVA<br>(Tukey post hoc test) | $T_{Reg}:T_{Conv}$ | Y60A IC vs. V103A IC | ns | 0.0741 | 4A |
| One-way ANOVA<br>(Tukey post hoc test) | $T_{Reg}:T_{Conv}$ | Y60A IC vs. Y33A IC | ** | 0.0052 | 4A |
| One-way ANOVA<br>(Tukey post hoc test) | $T_{Reg}:T_{Conv}$ | Y60A IC vs. Control IC | ns | 0.1887 | 4A |
| One-way ANOVA<br>(Tukey post hoc test) | $T_{Reg}:T_{Conv}$ | Y60A IC vs. F5111.2 IC | **** | <0.0001 | 4A |
| One-way ANOVA<br>(Tukey post hoc test) | $T_{Reg}:T_{Conv}$ | V103A IC vs. Y33A IC | ns | 0.9134 | 4A |
| One-way ANOVA<br>(Tukey post hoc test) | $T_{Reg}:T_{Conv}$ | V103A IC vs. Control IC | *** | 0.0004 | 4A |
| One-way ANOVA<br>(Tukey post hoc test) | $T_{Reg}:T_{Conv}$ | V103A IC vs. F5111.2 IC | **** | <0.0001 | 4A |
| One-way ANOVA<br>(Tukey post hoc test) | $T_{Reg}:T_{Conv}$ | Y33A IC vs. Control IC | **** | <0.0001 | 4A |
| One-way ANOVA<br>(Tukey post hoc test) | $T_{Reg}:T_{Conv}$ | Y33A IC vs. F5111.2 IC | **** | <0.0001 | 4A |
| One-way ANOVA<br>(Tukey post hoc test) | $T_{Reg}:T_{Conv}$ | Control IC vs. F5111.2 IC | * | 0.0207 | 4A |
| One-way ANOVA<br>(Tukey post hoc test) | $T_{Reg}:NK$ | PBS vs. F5111 IC | **** | <0.0001 | 4A |
| One-way ANOVA<br>(Tukey post hoc test) | $T_{Reg}:NK$ | PBS vs. Y60A IC | **** | <0.0001 | 4A |
| One-way ANOVA<br>(Tukey post hoc test) | $T_{Reg}:NK$ | PBS vs. V103A IC | **** | <0.0001 | 4A |
| One-way ANOVA<br>(Tukey post hoc test) | $T_{Reg}:NK$ | PBS vs. Y33A IC | **** | <0.0001 | 4A |
| One-way ANOVA<br>(Tukey post hoc test) | $T_{Reg}:NK$ | PBS vs. Control IC | ns | 0.9482 | 4A |
| One-way ANOVA<br>(Tukey post hoc test) | $T_{Reg}:NK$ | PBS vs. F5111.2 IC | ns | 0.0839 | 4A |
| One-way ANOVA<br>(Tukey post hoc test) | $T_{Reg}:NK$ | F5111 IC vs. Y60A IC | **** | <0.0001 | 4A |
| One-way ANOVA<br>(Tukey post hoc test) | $T_{Reg}:NK$ | F5111 IC vs. V103A IC | **** | <0.0001 | 4A |
| One-way ANOVA<br>(Tukey post hoc test) | $T_{Reg}:NK$ | F5111 IC vs. Y33A IC | **** | <0.0001 | 4A |
| One-way ANOVA<br>(Tukey post hoc test) | $T_{Reg}:NK$ | F5111 IC vs. Control IC | **** | <0.0001 | 4A |
| One-way ANOVA<br>(Tukey post hoc test) | $T_{Reg}:NK$ | F5111 IC vs. F5111.2 IC | **** | <0.0001 | 4A |
| One-way ANOVA<br>(Tukey post hoc test) | $T_{Reg}:NK$ | Y60A IC vs. V103A IC | ns | >0.9999 | 4A |
| One-way ANOVA<br>(Tukey post hoc test) | $T_{Reg}:NK$ | Y60A IC vs. Y33A IC | ns | >0.9999 | 4A |
| One-way ANOVA<br>(Tukey post hoc test) | $T_{Reg}:NK$ | Y60A IC vs. Control IC | **** | <0.0001 | 4A |
| One-way ANOVA<br>(Tukey post hoc test) | T<sub>Reg</sub>:NK | Y60A IC vs. F5111.2 IC | ** | 0.0013 | 4A |
| One-way ANOVA<br>(Tukey post hoc test) | T<sub>Reg</sub>:NK | V103A IC vs. Y33A IC | ns | >0.9999 | 4A |
| One-way ANOVA<br>(Tukey post hoc test) | T<sub>Reg</sub>:NK | V103A IC vs. Control IC | **** | <0.0001 | 4A |
| One-way ANOVA<br>(Tukey post hoc test) | T<sub>Reg</sub>:NK | V103A IC vs. F5111.2 IC | ** | 0.0031 | 4A |
| One-way ANOVA<br>(Tukey post hoc test) | T<sub>Reg</sub>:NK | Y33A IC vs. Control IC | **** | <0.0001 | 4A |
| One-way ANOVA<br>(Tukey post hoc test) | T<sub>Reg</sub>:NK | Y33A IC vs. F5111.2 IC | ** | 0.0039 | 4A |
| One-way ANOVA<br>(Tukey post hoc test) | T<sub>Reg</sub>:NK | Control IC vs. F5111.2 IC | ** | 0.0092 | 4A |
| One-way ANOVA<br>(Tukey post hoc test) | T<sub>Reg</sub> Number | PBS vs. F5111 IC | **** | <0.0001 | 5B |
| One-way ANOVA<br>(Tukey post hoc test) | T<sub>Reg</sub> Number | PBS vs. hIL-2/F5111.2 Cx | **** | <0.0001 | 5B |
| One-way ANOVA<br>(Tukey post hoc test) | T<sub>Reg</sub> Number | PBS vs. Control IC | **** | <0.0001 | 5B |
| One-way ANOVA<br>(Tukey post hoc test) | T<sub>Reg</sub> Number | F5111 IC vs. hIL-2/F5111.2 Cx | ns | 0.3358 | 5B |
| One-way ANOVA<br>(Tukey post hoc test) | T<sub>Reg</sub> Number | F5111 IC vs. Control IC | **** | <0.0001 | 5B |
| One-way ANOVA<br>(Tukey post hoc test) | T<sub>Reg</sub> Number | hIL-2/F5111.2 Cx vs. Control IC | **** | <0.0001 | 5B |
| One-way ANOVA<br>(Tukey post hoc test) | CD8<sup>+</sup> T Number | PBS vs. F5111 IC | *** | 0.0003 | 5B |
| One-way ANOVA<br>(Tukey post hoc test) | CD8<sup>+</sup> T Number | PBS vs. hIL-2/F5111.2 Cx | ns | 0.8267 | 5B |
| One-way ANOVA<br>(Tukey post hoc test) | CD8<sup>+</sup> T Number | PBS vs. Control IC | **** | <0.0001 | 5B |
| One-way ANOVA<br>(Tukey post hoc test) | CD8<sup>+</sup> T Number | F5111 IC vs. hIL-2/F5111.2 Cx | **** | <0.0001 | 5B |
| One-way ANOVA<br>(Tukey post hoc test) | CD8<sup>+</sup> T Number | F5111 IC vs. Control IC | **** | <0.0001 | 5B |
| One-way ANOVA<br>(Tukey post hoc test) | CD8<sup>+</sup> T Number | hIL-2/F5111.2 Cx vs. Control IC | **** | <0.0001 | 5B |
| One-way ANOVA<br>(Tukey post hoc test) | T<sub>Conv</sub> Number | PBS vs. F5111 IC | **** | <0.0001 | 5B |
| One-way ANOVA<br>(Tukey post hoc test) | T<sub>Conv</sub> Number | PBS vs. hIL-2/F5111.2 Cx | **** | <0.0001 | 5B |
| One-way ANOVA<br>(Tukey post hoc test) | T<sub>Conv</sub> Number | PBS vs. Control IC | ** | 0.0054 | 5B |
| One-way ANOVA<br>(Tukey post hoc test) | T<sub>Conv</sub> Number | F5111 IC vs. hIL-2/F5111.2 Cx | * | 0.0324 | 5B |
| One-way ANOVA<br>(Tukey post hoc test) | T<sub>Conv</sub> Number | F5111 IC vs. Control IC | **** | <0.0001 | 5B |
| One-way ANOVA<br>(Tukey post hoc test) | T<sub>Conv</sub> Number | hIL-2/F5111.2 Cx vs. Control IC | ** | 0.0030 | 5B |
| One-way ANOVA<br>(Tukey post hoc test) | NK Number | PBS vs. F5111 IC | ns | 0.5992 | 5B |
| One-way ANOVA<br>(Tukey post hoc test) | NK Number | PBS vs. hIL-2/F5111.2 Cx | * | 0.0106 | 5B |
| One-way ANOVA<br>(Tukey post hoc test) | NK Number | PBS vs. Control IC | **** | <0.0001 | 5B |
| One-way ANOVA<br>(Tukey post hoc test) | NK Number | F5111 IC vs. hIL-2/F5111.2 Cx | ns | 0.0950 | 5B |
| One-way ANOVA<br>(Tukey post hoc test) | NK Number | F5111 IC vs. Control IC | **** | <0.0001 | 5B |
| One-way ANOVA<br>(Tukey post hoc test) | NK Number | hIL-2/F5111.2 Cx vs. Control IC | **** | <0.0001 | 5B |
| One-way ANOVA<br>(Tukey post hoc test) | T<sub>Reg</sub> FOXP3 MFI | PBS vs. F5111 IC | **** | <0.0001 | 5C |
| One-way ANOVA<br>(Tukey post hoc test) | T<sub>Reg</sub> FOXP3 MFI | PBS vs. hIL-2/F5111.2 Cx | *** | 0.0001 | 5C |
| One-way ANOVA<br>(Tukey post hoc test) | T<sub>Reg</sub> FOXP3 MFI | PBS vs. Control IC | * | 0.0460 | 5C |
| One-way ANOVA<br>(Tukey post hoc test) | T<sub>Reg</sub> FOXP3 MFI | F5111 IC vs. hIL-2/F5111.2 Cx | ** | 0.0094 | 5C |
| One-way ANOVA<br>(Tukey post hoc test) | T<sub>Reg</sub> FOXP3 MFI | F5111 IC vs. Control IC | **** | <0.0001 | 5C |
| One-way ANOVA<br>(Tukey post hoc test) | T<sub>Reg</sub> FOXP3 MFI | hIL-2/F5111.2 Cx vs. Control IC | * | 0.0197 | 5C |
| One-way ANOVA<br>(Tukey post hoc test) | T<sub>Reg</sub> IL-2Rα MFI | PBS vs. F5111 IC | **** | <0.0001 | 5C |
| One-way ANOVA<br>(Tukey post hoc test) | T<sub>Reg</sub> IL-2Rα MFI | PBS vs. hIL-2/F5111.2 Cx | **** | <0.0001 | 5C |
| One-way ANOVA<br>(Tukey post hoc test) | T<sub>Reg</sub> IL-2Rα MFI | PBS vs. Control IC | ** | 0.0062 | 5C |
| One-way ANOVA<br>(Tukey post hoc test) | T<sub>Reg</sub> IL-2Rα MFI | F5111 IC vs. hIL-2/F5111.2 Cx | ns | 0.1415 | 5C |
| One-way ANOVA<br>(Tukey post hoc test) | T<sub>Reg</sub> IL-2Rα MFI | F5111 IC vs. Control IC | **** | <0.0001 | 5C |
| One-way ANOVA<br>(Tukey post hoc test) | T<sub>Reg</sub> IL-2Rα MFI | hIL-2/F5111.2 Cx vs. Control IC | **** | <0.0001 | 5C |
| One-way ANOVA<br>(Tukey post hoc test) | CD8<sup>+</sup> T IL-2Rα MFI | PBS vs. F5111 IC | **** | <0.0001 | 5C |
| One-way ANOVA<br>(Tukey post hoc test) | CD8<sup>+</sup> T IL-2Rα MFI | PBS vs. hIL-2/F5111.2 Cx | *** | 0.0003 | 5C |
| One-way ANOVA<br>(Tukey post hoc test) | CD8<sup>+</sup> T IL-2Rα MFI | PBS vs. Control IC | ns | 0.2575 | 5C |
| One-way ANOVA<br>(Tukey post hoc test) | CD8<sup>+</sup> T IL-2Rα MFI | F5111 IC vs. hIL-2/F5111.2 Cx | ** | 0.0088 | 5C |
| One-way ANOVA<br>(Tukey post hoc test) | CD8<sup>+</sup> T IL-2Rα MFI | F5111 IC vs. Control IC | **** | <0.0001 | 5C |
| One-way ANOVA<br>(Tukey post hoc test) | CD8<sup>+</sup> T IL-2Rα MFI | hIL-2/F5111.2 Cx vs. Control IC | ** | 0.0072 | 5C |
| One-way ANOVA<br>(Tukey post hoc test) | T<sub>Conv</sub> IL-2Rα MFI | PBS vs. F5111 IC | *** | 0.0008 | 5C |
| One-way ANOVA<br>(Tukey post hoc test) | T<sub>Conv</sub> IL-2Rα MFI | PBS vs. hIL-2/F5111.2 Cx | *** | 0.0007 | 5C |
| One-way ANOVA<br>(Tukey post hoc test) | T<sub>Conv</sub> IL-2Rα MFI | PBS vs. Control IC | ns | 0.1177 | 5C |
| One-way ANOVA<br>(Tukey post hoc test) | T<sub>Conv</sub> IL-2Rα MFI | F5111 IC vs. hIL-2/F5111.2 Cx | ns | 0.9999 | 5C |
| One-way ANOVA<br>(Tukey post hoc test) | T<sub>Conv</sub> IL-2Rα MFI | F5111 IC vs. Control IC | ns | 0.0512 | 5C |
| One-way ANOVA<br>(Tukey post hoc test) | T<sub>Conv</sub> IL-2Rα MFI | hIL-2/F5111.2 Cx vs. Control IC | * | 0.0456 | 5C |
| One-way ANOVA<br>(Tukey post hoc test) | T<sub>Reg</sub>:CD8<sup>+</sup> T | PBS vs. F5111 IC | **** | <0.0001 | 5D |
| One-way ANOVA<br>(Tukey post hoc test) | T<sub>Reg</sub>:CD8<sup>+</sup> T | PBS vs. hIL-2/F5111.2 Cx | **** | <0.0001 | 5D |
| One-way ANOVA<br>(Tukey post hoc test) | T<sub>Reg</sub>:CD8<sup>+</sup> T | PBS vs. Control IC | * | 0.0378 | 5D |
| One-way ANOVA<br>(Tukey post hoc test) | T<sub>Reg</sub>:CD8<sup>+</sup> T | F5111 IC vs. hIL-2/F5111.2 Cx | **** | <0.0001 | 5D |
| One-way ANOVA<br>(Tukey post hoc test) | T<sub>Reg</sub>:CD8<sup>+</sup> T | F5111 IC vs. Control IC | **** | <0.0001 | 5D |
| One-way ANOVA<br>(Tukey post hoc test) | T<sub>Reg</sub>:CD8<sup>+</sup> T | hIL-2/F5111.2 Cx vs. Control IC | **** | <0.0001 | 5D |
| One-way ANOVA<br>(Tukey post hoc test) | T<sub>Reg</sub>:T<sub>Conv</sub> | PBS vs. F5111 IC | **** | <0.0001 | 5D |
| One-way ANOVA<br>(Tukey post hoc test) | T<sub>Reg</sub>:T<sub>Conv</sub> | PBS vs. hIL-2/F5111.2 Cx | **** | <0.0001 | 5D |
| One-way ANOVA<br>(Tukey post hoc test) | T<sub>Reg</sub>:T<sub>Conv</sub> | PBS vs. Control IC | ns | 0.1868 | 5D |
| One-way ANOVA<br>(Tukey post hoc test) | T<sub>Reg</sub>:T<sub>Conv</sub> | F5111 IC vs. hIL-2/F5111.2 Cx | ** | 0.0013 | 5D |
| One-way ANOVA<br>(Tukey post hoc test) | T<sub>Reg</sub>:T<sub>Conv</sub> | F5111 IC vs. Control IC | **** | <0.0001 | 5D |
| One-way ANOVA<br>(Tukey post hoc test) | T<sub>Reg</sub>:T<sub>Conv</sub> | hIL-2/F5111.2 Cx vs. Control IC | **** | <0.0001 | 5D |
| One-way ANOVA<br>(Tukey post hoc test) | T<sub>Reg</sub>:NK | PBS vs. F5111 IC | **** | <0.0001 | 5D |
| One-way ANOVA<br>(Tukey post hoc test) | T<sub>Reg</sub>:NK | PBS vs. hIL-2/F5111.2 Cx | **** | <0.0001 | 5D |
| One-way ANOVA<br>(Tukey post hoc test) | T<sub>Reg</sub>:NK | PBS vs. Control IC | ns | 0.9893 | 5D |
| One-way ANOVA<br>(Tukey post hoc test) | T<sub>Reg</sub>:NK | F5111 IC vs. hIL-2/F5111.2 Cx | *** | 0.0003 | 5D |
| One-way ANOVA<br>(Tukey post hoc test) | T<sub>Reg</sub>:NK | F5111 IC vs. Control IC | **** | <0.0001 | 5D |
| One-way ANOVA<br>(Tukey post hoc test) | T<sub>Reg</sub>:NK | hIL-2/F5111.2 Cx vs. Control IC | **** | <0.0001 | 5D |
| One-way ANOVA<br>(Tukey post hoc test) | %T<sub>Reg</sub> [CD4] | PBS vs. F5111 IC | **** | <0.0001 | 5E |
| One-way ANOVA<br>(Tukey post hoc test) | %T<sub>Reg</sub> [CD4] | PBS vs. hIL-2/F5111.2 Cx | **** | <0.0001 | 5E |
| One-way ANOVA<br>(Tukey post hoc test) | %T<sub>Reg</sub> [CD4] | PBS vs. Control IC | **** | <0.0001 | 5E |
| One-way ANOVA<br>(Tukey post hoc test) | %T<sub>Reg</sub> [CD4] | F5111 IC vs. hIL-2/F5111.2 Cx | ** | 0.0030 | 5E |
| One-way ANOVA<br>(Tukey post hoc test) | %T<sub>Reg</sub> [CD4] | F5111 IC vs. Control IC | **** | <0.0001 | 5E |
| One-way ANOVA<br>(Tukey post hoc test) | %T<sub>Reg</sub> [CD4] | hIL-2/F5111.2 Cx vs. Control IC | **** | <0.0001 | 5E |
| One-way ANOVA<br>(Tukey post hoc test) | %Ki-67 [T<sub>Reg</sub>] | PBS vs. F5111 IC | **** | <0.0001 | 5F |
| One-way ANOVA<br>(Tukey post hoc test) | %Ki-67 [T<sub>Reg</sub>] | PBS vs. hIL-2/F5111.2 Cx | **** | <0.0001 | 5F |
| One-way ANOVA<br>(Tukey post hoc test) | %Ki-67 [T<sub>Reg</sub>] | PBS vs. Control IC | **** | <0.0001 | 5F |
| One-way ANOVA<br>(Tukey post hoc test) | %Ki-67 [T<sub>Reg</sub>] | F5111 IC vs. hIL-2/F5111.2 Cx | **** | <0.0001 | 5F |
| One-way ANOVA<br>(Tukey post hoc test) | %Ki-67 [T<sub>Reg</sub>] | F5111 IC vs. Control IC | **** | <0.0001 | 5F |
| One-way ANOVA<br>(Tukey post hoc test) | %Ki-67 [T<sub>Reg</sub>] | hIL-2/F5111.2 Cx vs. Control IC | **** | <0.0001 | 5F |
| One-way ANOVA<br>(Tukey post hoc test) | %Ki-67 [CD8<sup>+</sup> T] | PBS vs. F5111 IC | **** | <0.0001 | 5F |
| One-way ANOVA<br>(Tukey post hoc test) | %Ki-67 [CD8<sup>+</sup> T] | PBS vs. hIL-2/F5111.2 Cx | **** | <0.0001 | 5F |
| One-way ANOVA<br>(Tukey post hoc test) | %Ki-67 [CD8<sup>+</sup> T] | PBS vs. Control IC | **** | <0.0001 | 5F |
| One-way ANOVA<br>(Tukey post hoc test) | %Ki-67 [CD8<sup>+</sup> T] | F5111 IC vs. hIL-2/F5111.2 Cx | **** | <0.0001 | 5F |
| One-way ANOVA<br>(Tukey post hoc test) | %Ki-67 [CD8<sup>+</sup> T] | F5111 IC vs. Control IC | **** | <0.0001 | 5F |
| One-way ANOVA<br>(Tukey post hoc test) | %Ki-67 [CD8<sup>+</sup> T] | hIL-2/F5111.2 Cx vs. Control IC | ** | 0.0073 | 5F |
| One-way ANOVA<br>(Tukey post hoc test) | %Ki-67 [T<sub>Conv</sub>] | PBS vs. F5111 IC | * | 0.0130 | 5F |
| One-way ANOVA<br>(Tukey post hoc test) | %Ki-67 [T<sub>Conv</sub>] | PBS vs. hIL-2/F5111.2 Cx | **** | <0.0001 | 5F |
| One-way ANOVA<br>(Tukey post hoc test) | %Ki-67 [T<sub>Conv</sub>] | PBS vs. Control IC | **** | <0.0001 | 5F |
| One-way ANOVA<br>(Tukey post hoc test) | %Ki-67 [T<sub>Conv</sub>] | F5111 IC vs. hIL-2/F5111.2 Cx | **** | <0.0001 | 5F |
| One-way ANOVA<br>(Tukey post hoc test) | %Ki-67 [T<sub>Conv</sub>] | F5111 IC vs. Control IC | **** | <0.0001 | 5F |
| One-way ANOVA<br>(Tukey post hoc test) | %Ki-67 [T<sub>Conv</sub>] | hIL-2/F5111.2 Cx vs. Control IC | ns | 0.4734 | 5F |
| One-way ANOVA<br>(Tukey post hoc test) | %Ki-67 [NK] | PBS vs. F5111 IC | ns | 0.9428 | 5F |
| One-way ANOVA<br>(Tukey post hoc test) | %Ki-67 [NK] | PBS vs. hIL-2/F5111.2 Cx | **** | <0.0001 | 5F |
| One-way ANOVA<br>(Tukey post hoc test) | %Ki-67 [NK] | PBS vs. Control IC | **** | <0.0001 | 5F |
| One-way ANOVA<br>(Tukey post hoc test) | %Ki-67 [NK] | F5111 IC vs. hIL-2/F5111.2 Cx | **** | <0.0001 | 5F |
| One-way ANOVA<br>(Tukey post hoc test) | %Ki-67 [NK] | F5111 IC vs. Control IC | **** | <0.0001 | 5F |
| One-way ANOVA<br>(Tukey post hoc test) | %Ki-67 [NK] | hIL-2/F5111.2 Cx vs. Control IC | *** | 0.0005 | 5F |
| Student *t* test | T<sub>Reg</sub>:T<sub>Conv</sub> (1:16) | PBS + F5111 IC | * | 0.0319 | 6E |
| Student *t* test | T<sub>Reg</sub>:T<sub>Conv</sub> (1:8) | PBS + F5111 IC | ns | 0.1723 | 6E |
| Student *t* test | T<sub>Reg</sub>:T<sub>Conv</sub> (1:4) | PBS + F5111 IC | * | 0.0212 | 6E |
| Student *t* test | T<sub>Reg</sub>:T<sub>Conv</sub> (1:2) | PBS + F5111 IC | * | 0.0468 | 6E |
| Student *t* test | T<sub>Reg</sub>:T<sub>Conv</sub> (1:1) | PBS + F5111 IC | ns | 0.1991 | 6E |
| Student *t* test | T<sub>Reg</sub>:T<sub>Conv</sub> (2:1) | PBS + F5111 IC | ns | 0.9257 | 6E |
| One-way ANOVA<br>(Tukey post hoc test) | cT<sub>Reg</sub> Number | PBS vs. Control IC | ns | 0.1129 | 6F |
| One-way ANOVA<br>(Tukey post hoc test) | cT<sub>Reg</sub> Number | PBS vs. F5111 IC | ** | 0.0011 | 6F |
| One-way ANOVA<br>(Tukey post hoc test) | cT<sub>Reg</sub> Number | Control IC vs. F5111 IC | **** | <0.0001 | 6F |
| One-way ANOVA<br>(Tukey post hoc test) | eT<sub>Reg</sub> Number | PBS vs. Control IC | ns | 0.1154 | 6F |
| One-way ANOVA<br>(Tukey post hoc test) | eT<sub>Reg</sub> Number | PBS vs. F5111 IC | ns | 0.7387 | 6F |
| One-way ANOVA<br>(Tukey post hoc test) | eT<sub>Reg</sub> Number | Control IC vs. F5111 IC | ns | 0.3580 | 6F |
| One-way ANOVA<br>(Tukey post hoc test) | % Initial Weight (D10) | PBS vs. Control IC | ** | 0.0023 | 6G |
| One-way ANOVA<br>(Tukey post hoc test) | % Initial Weight (D10) | PBS vs. F5111 IC | ** | 0.0019 | 6G |
| One-way ANOVA<br>(Tukey post hoc test) | % Initial Weight (D10) | PBS vs. Control | **** | <0.0001 | 6G |
| One-way ANOVA<br>(Tukey post hoc test) | % Initial Weight (D10) | Control IC vs. F5111 IC | ns | 0.9996 | 6G |
| One-way ANOVA<br>(Tukey post hoc test) | % Initial Weight (D10) | Control IC vs. Control | ** | 0.0031 | 6G |
| One-way ANOVA<br>(Tukey post hoc test) | % Initial Weight (D10) | F5111 IC vs. Control | ** | 0.0038 | 6G |
| One-way ANOVA<br>(Tukey post hoc test) | Parasite Burden (Liver) | PBS vs. Control IC | ** | 0.0029 | 6H |
| One-way ANOVA<br>(Tukey post hoc test) | Parasite Burden (Liver) | PBS vs. F5111 IC | ** | 0.0015 | 6H |
| One-way ANOVA<br>(Tukey post hoc test) | Parasite Burden (Liver) | Control IC vs. F5111 IC | ns | 0.9132 | 6H |
| One-way ANOVA<br>(Tukey post hoc test) | Parasite Burden (Lungs) | PBS vs. Control IC | ns | 0.9803 | 6H |
| One-way ANOVA<br>(Tukey post hoc test) | Parasite Burden (Lungs) | PBS vs. F5111 IC | ns | 0.4249 | 6H |
| One-way ANOVA<br>(Tukey post hoc test) | Parasite Burden (Lungs) | Control IC vs. F5111 IC | ns | 0.4908 | 6H |
| One-way ANOVA<br>(Tukey post hoc test) | T<sub>Reg</sub>:T-bet<sup>+</sup>Tet<sup>+</sup>CD8<sup>+</sup> T | PBS vs. Control IC | ns | 0.2787 | 6I |
| One-way ANOVA<br>(Tukey post hoc test) | T<sub>Reg</sub>:T-bet<sup>+</sup>Tet<sup>+</sup>CD8<sup>+</sup> T | PBS vs. F5111 IC | * | 0.0418 | 6I |
| One-way ANOVA<br>(Tukey post hoc test) | T<sub>Reg</sub>:T-bet<sup>+</sup>Tet<sup>+</sup>CD8<sup>+</sup> T | Control IC vs. F5111 IC | ns | 0.4457 | 6I |
| One-way ANOVA<br>(Tukey post hoc test) | % Initial Weight (D15) | PBS + DSS vs. hIL-2/F5111.2<br>Cx + DSS | ** | 0.0013 | 7B |
| One-way ANOVA<br>(Tukey post hoc test) | % Initial Weight (D15) | PBS + DSS vs. F5111 IC +<br>DSS | *** | 0.0007 | 7B |
| One-way ANOVA<br>(Tukey post hoc test) | % Initial Weight (D15) | PBS + DSS vs. Control IC +<br>DSS | ns | 0.9961 | 7B |
| One-way ANOVA<br>(Tukey post hoc test) | % Initial Weight (D15) | PBS + DSS vs. Control | **** | <0.0001 | 7B |
| One-way ANOVA<br>(Tukey post hoc test) | % Initial Weight (D15) | hIL-2/F5111.2 Cx + DSS vs.<br>F5111 IC + DSS | ns | 0.9988 | 7B |
| One-way ANOVA<br>(Tukey post hoc test) | % Initial Weight (D15) | hIL-2/F5111.2 Cx + DSS vs.<br>Control IC + DSS | *** | 0.0005 | 7B |
| One-way ANOVA<br>(Tukey post hoc test) | % Initial Weight (D15) | hIL-2/F5111.2 Cx + DSS vs.<br>Control | *** | 0.0006 | 7B |
| One-way ANOVA<br>(Tukey post hoc test) | % Initial Weight (D15) | F5111 IC + DSS vs. Control IC<br>+ DSS | *** | 0.0003 | 7B |
| One-way ANOVA<br>(Tukey post hoc test) | % Initial Weight (D15) | F5111 IC + DSS vs. Control | ** | 0.0012 | 7B |
| One-way ANOVA<br>(Tukey post hoc test) | % Initial Weight (D15) | Control IC + DSS vs. Control | **** | <0.0001 | 7B |
| One-way ANOVA<br>(Tukey post hoc test) | DAI (D15) | PBS + DSS vs. hIL-2/F5111.2<br>Cx + DSS | ns | 0.1507 | 7C |
| One-way ANOVA<br>(Tukey post hoc test) | DAI (D15) | PBS + DSS vs. F5111 IC +<br>DSS | ns | 0.1507 | 7C |
| One-way ANOVA<br>(Tukey post hoc test) | DAI (D15) | PBS + DSS vs. Control IC +<br>DSS | ns | 0.9954 | 7C |
| One-way ANOVA<br>(Tukey post hoc test) | DAI (D15) | PBS + DSS vs. Control | **** | <0.0001 | 7C |
| One-way ANOVA<br>(Tukey post hoc test) | DAI (D15) | hIL-2/F5111.2 Cx + DSS vs.<br>F5111 IC + DSS | ns | >0.9999 | 7C |
| One-way ANOVA<br>(Tukey post hoc test) | DAI (D15) | hIL-2/F5111.2 Cx + DSS vs.<br>Control IC + DSS | ns | 0.2886 | 7C |
| One-way ANOVA<br>(Tukey post hoc test) | DAI (D15) | hIL-2/F5111.2 Cx + DSS vs.<br>Control | **** | <0.0001 | 7C |
| One-way ANOVA<br>(Tukey post hoc test) | DAI (D15) | F5111 IC + DSS vs. Control IC<br>+ DSS | ns | 0.2886 | 7C |
| One-way ANOVA<br>(Tukey post hoc test) | DAI (D15) | F5111 IC + DSS vs. Control | **** | <0.0001 | 7C |
| One-way ANOVA<br>(Tukey post hoc test) | DAI (D15) | Control IC + DSS vs. Control | **** | <0.0001 | 7C |
| One-way ANOVA<br>(Tukey post hoc test) | Colon Length | PBS + DSS vs. hIL-2/F5111.2<br>Cx + DSS | ns | 0.5874 | 7D |
| One-way ANOVA<br>(Tukey post hoc test) | Colon Length | PBS + DSS vs. F5111 IC +<br>DSS | ns | 0.2207 | 7D |
| One-way ANOVA<br>(Tukey post hoc test) | Colon Length | PBS + DSS vs. Control IC +<br>DSS | ns | 0.9999 | 7D |
| One-way ANOVA<br>(Tukey post hoc test) | Colon Length | PBS + DSS vs. Control | **** | <0.0001 | 7D |
| One-way ANOVA<br>(Tukey post hoc test) | Colon Length | hIL-2/F5111.2 Cx + DSS vs.<br>F5111 IC + DSS | ns | 0.9532 | 7D |
| One-way ANOVA<br>(Tukey post hoc test) | Colon Length | hIL-2/F5111.2 Cx + DSS vs.<br>Control IC + DSS | ns | 0.7207 | 7D |
| One-way ANOVA<br>(Tukey post hoc test) | Colon Length | hIL-2/F5111.2 Cx + DSS vs.<br>Control | *** | 0.0001 | 7D |
| One-way ANOVA<br>(Tukey post hoc test) | Colon Length | F5111 IC + DSS vs. Control IC<br>+ DSS | ns | 0.3310 | 7D |
| One-way ANOVA<br>(Tukey post hoc test) | Colon Length | F5111 IC + DSS vs. Control | *** | 0.0008 | 7D |
| One-way ANOVA<br>(Tukey post hoc test) | Colon Length | Control IC + DSS vs. Control | **** | <0.0001 | 7D |
| One-way ANOVA<br>(Tukey post hoc test) | Histopathology Score | PBS + DSS vs. hIL-2/F5111.2<br>Cx + DSS | ** | 0.0096 | 7E |
| One-way ANOVA<br>(Tukey post hoc test) | Histopathology Score | PBS + DSS vs. F5111 IC +<br>DSS | ns | 0.1352 | 7E |
| One-way ANOVA<br>(Tukey post hoc test) | Histopathology Score | PBS + DSS vs. Control IC +<br>DSS | ns | 0.2493 | 7E |
| One-way ANOVA<br>(Tukey post hoc test) | Histopathology Score | PBS + DSS vs. Control | **** | <0.0001 | 7E |
| One-way ANOVA<br>(Tukey post hoc test) | Histopathology Score | hIL-2/F5111.2 Cx + DSS vs.<br>F5111 IC + DSS | ns | 0.7344 | 7E |
| One-way ANOVA<br>(Tukey post hoc test) | Histopathology Score | hIL-2/F5111.2 Cx + DSS vs.<br>Control IC + DSS | ns | 0.6293 | 7E |
| One-way ANOVA<br>(Tukey post hoc test) | Histopathology Score | hIL-2/F5111.2 Cx + DSS vs.<br>Control | ** | 0.0015 | 7E |
| One-way ANOVA<br>(Tukey post hoc test) | Histopathology Score | F5111 IC + DSS vs. Control IC<br>+ DSS | ns | 0.9993 | 7E |
| One-way ANOVA<br>(Tukey post hoc test) | Histopathology Score | F5111 IC + DSS vs. Control | **** | <0.0001 | 7E |
| One-way ANOVA<br>(Tukey post hoc test) | Histopathology Score | Control IC + DSS vs. Control | **** | <0.0001 | 7E |
| Log-rank (Mantel-Cox)<br>test | % Diabetes-Free | anti-PD-1 vs. F5111 IC + anti-PD-1 | ** | 0.0078 | 8C |
| Log-rank (Mantel-Cox)<br>test | % Diabetes-Free | anti-PD-1 vs. Control | **** | <0.0001 | 8C |
| Log-rank (Mantel-Cox)<br>test | % Diabetes-Free | F5111 IC + anti-PD-1 vs.<br>Control | * | 0.0291 | 8C |
| Log-rank (Mantel-Cox)<br>test | % Diabetes-Free | F5111 IC + anti-PD-1 vs.<br>Control IC + anti-PD-1 | ns | 0.2378 | 8C |
| Log-rank (Mantel-Cox) test | % Diabetes-Free | Control IC + anti-PD-1 vs. Control | ** | 0.0040 | 8C |
| Log-rank (Mantel-Cox) test | % Diabetes-Free | Control IC + anti-PD-1 vs. PBS + anti-PD-1 | ns | 0.4378 | 8C |
| Log-rank (Mantel-Cox) test | % Diabetes-Free | anti-PD-1 vs. F5111 IC + anti-PD-1 | * | 0.0446 | 8F |
| Log-rank (Mantel-Cox) test | % Diabetes-Free | anti-PD-1 vs. Control | ns | 0.1784 | 8F |
| Log-rank (Mantel-Cox) test | % Diabetes-Free | F5111 IC + anti-PD-1 vs. Control | ns | 0.1660 | 8F |
| One-way ANOVA (Tukey post hoc test) | T<sub>Reg</sub>:CD8<sup>+</sup> T | PBS vs. F5111.2 IC | ns | 0.8826 | S4F |
| One-way ANOVA (Tukey post hoc test) | T<sub>Reg</sub>:CD8<sup>+</sup> T | PBS vs. F5111 IC | ns | 0.8794 | S4F |
| One-way ANOVA (Tukey post hoc test) | T<sub>Reg</sub>:CD8<sup>+</sup> T | PBS vs. Y33A IC | * | 0.0496 | S4F |
| One-way ANOVA (Tukey post hoc test) | T<sub>Reg</sub>:CD8<sup>+</sup> T | PBS vs. Y60A IC | ns | 0.2203 | S4F |
| One-way ANOVA (Tukey post hoc test) | T<sub>Reg</sub>:CD8<sup>+</sup> T | F5111.2 IC vs. F5111 IC | ns | >0.9999 | S4F |
| One-way ANOVA (Tukey post hoc test) | T<sub>Reg</sub>:CD8<sup>+</sup> T | F5111.2 IC vs. Y33A IC | ns | 0.2768 | S4F |
| One-way ANOVA (Tukey post hoc test) | T<sub>Reg</sub>:CD8<sup>+</sup> T | F5111.2 IC vs. Y60A IC | ns | 0.7389 | S4F |
| One-way ANOVA (Tukey post hoc test) | T<sub>Reg</sub>:CD8<sup>+</sup> T | F5111 IC vs. Y33A IC | ns | 0.3532 | S4F |
| One-way ANOVA (Tukey post hoc test) | T<sub>Reg</sub>:CD8<sup>+</sup> T | F5111 IC vs. Y60A IC | ns | 0.8097 | S4F |
| One-way ANOVA (Tukey post hoc test) | T<sub>Reg</sub>:CD8<sup>+</sup> T | Y33A IC vs. Y60A IC | ns | 0.8903 | S4F |
| One-way ANOVA (Tukey post hoc test) | T<sub>Reg</sub>:T<sub>Conv</sub> | PBS vs. F5111.2 IC | ns | 0.9846 | S4F |
| One-way ANOVA (Tukey post hoc test) | T<sub>Reg</sub>:T<sub>Conv</sub> | PBS vs. F5111 IC | ns | 0.6317 | S4F |
| One-way ANOVA (Tukey post hoc test) | T<sub>Reg</sub>:T<sub>Conv</sub> | PBS vs. Y33A IC | ns | 0.1555 | S4F |
| One-way ANOVA (Tukey post hoc test) | T<sub>Reg</sub>:T<sub>Conv</sub> | PBS vs. Y60A IC | * | 0.0384 | S4F |
| One-way ANOVA (Tukey post hoc test) | T<sub>Reg</sub>:T<sub>Conv</sub> | F5111.2 IC vs. F5111 IC | ns | 0.8984 | S4F |
| One-way ANOVA (Tukey post hoc test) | T<sub>Reg</sub>:T<sub>Conv</sub> | F5111.2 IC vs. Y33A IC | ns | 0.3746 | S4F |
| One-way ANOVA (Tukey post hoc test) | T<sub>Reg</sub>:T<sub>Conv</sub> | F5111.2 IC vs. Y60A IC | ns | 0.1302 | S4F |
| One-way ANOVA (Tukey post hoc test) | T<sub>Reg</sub>:T<sub>Conv</sub> | F5111 IC vs. Y33A IC | ns | 0.8870 | S4F |
| One-way ANOVA (Tukey post hoc test) | T<sub>Reg</sub>:T<sub>Conv</sub> | F5111 IC vs. Y60A IC | ns | 0.5871 | S4F |
| One-way ANOVA (Tukey post hoc test) | T<sub>Reg</sub>:T<sub>Conv</sub> | Y33A IC vs. Y60A IC | ns | 0.9850 | S4F |
| One-way ANOVA (Tukey post hoc test) | T<sub>Reg</sub>:CD8<sup>+</sup> T | PBS vs. F5111 IC | **** | <0.0001 | S5B |
| One-way ANOVA (Tukey post hoc test) | T<sub>Reg</sub>:CD8<sup>+</sup> T | PBS vs. F5111 IC w/o N297A | **** | <0.0001 | S5B |
| One-way ANOVA (Tukey post hoc test) | T<sub>Reg</sub>:CD8<sup>+</sup> T | PBS vs. Control IC | ns | 0.2006 | S5B |
| One-way ANOVA (Tukey post hoc test) | T<sub>Reg</sub>:CD8<sup>+</sup> T | F5111 IC vs. F5111 IC w/o N297A | **** | <0.0001 | S5B |
| One-way ANOVA<br>(Tukey post hoc test) | T<sub>Reg</sub>:CD8<sup>+</sup> T | F5111 IC vs. Control IC | **** | <0.0001 | S5B |
| One-way ANOVA<br>(Tukey post hoc test) | T<sub>Reg</sub>:CD8<sup>+</sup> T | F5111 IC w/o N297A vs.<br>Control IC | *** | 0.0002 | S5B |
| One-way ANOVA<br>(Tukey post hoc test) | T<sub>Reg</sub>:T<sub>Conv</sub> | PBS vs. F5111 IC | **** | <0.0001 | S5B |
| One-way ANOVA<br>(Tukey post hoc test) | T<sub>Reg</sub>:T<sub>Conv</sub> | PBS vs. F5111 IC w/o N297A | ** | 0.0080 | S5B |
| One-way ANOVA<br>(Tukey post hoc test) | T<sub>Reg</sub>:T<sub>Conv</sub> | PBS vs. Control IC | ns | 0.1738 | S5B |
| One-way ANOVA<br>(Tukey post hoc test) | T<sub>Reg</sub>:T<sub>Conv</sub> | F5111 IC vs. F5111 IC w/o<br>N297A | **** | <0.0001 | S5B |
| One-way ANOVA<br>(Tukey post hoc test) | T<sub>Reg</sub>:T<sub>Conv</sub> | F5111 IC vs. Control IC | **** | <0.0001 | S5B |
| One-way ANOVA<br>(Tukey post hoc test) | T<sub>Reg</sub>:T<sub>Conv</sub> | F5111 IC w/o N297A vs.<br>Control IC | ns | 0.3197 | S5B |
| One-way ANOVA<br>(Tukey post hoc test) | T<sub>Reg</sub>:NK | PBS vs. F5111 IC | **** | <0.0001 | S5B |
| One-way ANOVA<br>(Tukey post hoc test) | T<sub>Reg</sub>:NK | PBS vs. F5111 IC w/o N297A | * | 0.0278 | S5B |
| One-way ANOVA<br>(Tukey post hoc test) | T<sub>Reg</sub>:NK | PBS vs. Control IC | ns | 0.9899 | S5B |
| One-way ANOVA<br>(Tukey post hoc test) | T<sub>Reg</sub>:NK | F5111 IC vs. F5111 IC w/o<br>N297A | **** | <0.0001 | S5B |
| One-way ANOVA<br>(Tukey post hoc test) | T<sub>Reg</sub>:NK | F5111 IC vs. Control IC | **** | <0.0001 | S5B |
| One-way ANOVA<br>(Tukey post hoc test) | T<sub>Reg</sub>:NK | F5111 IC w/o N297A vs.<br>Control IC | * | 0.0164 | S5B |
| One-way ANOVA<br>(Tukey post hoc test) | T<sub>Reg</sub> Number | PBS vs. F5111 IC | **** | <0.0001 | S5B |
| One-way ANOVA<br>(Tukey post hoc test) | T<sub>Reg</sub> Number | PBS vs. F5111 IC w/o N297A | **** | <0.0001 | S5B |
| One-way ANOVA<br>(Tukey post hoc test) | T<sub>Reg</sub> Number | PBS vs. Control IC | **** | <0.0001 | S5B |
| One-way ANOVA<br>(Tukey post hoc test) | T<sub>Reg</sub> Number | F5111 IC vs. F5111 IC w/o<br>N297A | **** | <0.0001 | S5B |
| One-way ANOVA<br>(Tukey post hoc test) | T<sub>Reg</sub> Number | F5111 IC vs. Control IC | **** | <0.0001 | S5B |
| One-way ANOVA<br>(Tukey post hoc test) | T<sub>Reg</sub> Number | F5111 IC w/o N297A vs.<br>Control IC | * | 0.0262 | S5B |
| One-way ANOVA<br>(Tukey post hoc test) | %T<sub>Reg</sub> [CD4] | PBS vs. 1.5 µg IL-2 eq. | * | 0.0124 | S5D (left) |
| One-way ANOVA<br>(Tukey post hoc test) | %T<sub>Reg</sub> [CD4] | PBS vs. 0.5 µg IL-2 eq. | ns | 0.1149 | S5D (left) |
| One-way ANOVA<br>(Tukey post hoc test) | %T<sub>Reg</sub> [CD4] | PBS vs. 0.167 µg IL-2 eq. | ns | 0.1643 | S5D (left) |
| One-way ANOVA<br>(Tukey post hoc test) | %T<sub>Reg</sub> [CD4] | 1.5 µg IL-2 eq. vs. 0.5 µg IL-2<br>eq. | ns | 0.1222 | S5D (left) |
| One-way ANOVA<br>(Tukey post hoc test) | %T<sub>Reg</sub> [CD4] | 1.5 µg IL-2 eq. vs. 0.167 µg IL-2 eq. | ns | 0.0866 | S5D (left) |
| One-way ANOVA<br>(Tukey post hoc test) | %T<sub>Reg</sub> [CD4] | 0.5 µg IL-2 eq. vs. 0.167 µg IL-2 eq. | ns | 0.9777 | S5D (left) |
| One-way ANOVA<br>(Tukey post hoc test) | %T<sub>Reg</sub> [CD4] | PBS vs. 1.5 µg IL-2 eq. | *** | 0.0003 | S5D<br>(right) |
| One-way ANOVA<br>(Tukey post hoc test) | %T<sub>Reg</sub> [CD4] | PBS vs. 1.125 µg IL-2 eq. | *** | 0.0005 | S5D<br>(right) |
| One-way ANOVA<br>(Tukey post hoc test) | %T<sub>Reg</sub> [CD4] | PBS vs. 0.75 µg IL-2 eq. | *** | 0.0008 | S5D<br>(right) |
| One-way ANOVA<br>(Tukey post hoc test) | %T<sub>Reg</sub> [CD4] | 1.5 µg IL-2 eq. vs. 1.125 µg IL-2 eq. | ns | 0.4529 | S5D<br>(right) |
| One-way ANOVA<br>(Tukey post hoc test) | %T<sub>Reg</sub> [CD4] | 1.5 µg IL-2 eq. vs. 0.75 µg IL-2 eq. | ns | 0.1263 | S5D<br>(right) |
| One-way ANOVA<br>(Tukey post hoc test) | %T<sub>Reg</sub> [CD4] | 1.125 µg IL-2 eq. vs. 0.75 µg IL-2 eq. | ns | 0.5862 | S5D<br>(right) |
| One-way ANOVA<br>(Tukey post hoc test) | T<sub>Reg</sub> Number | PBS vs. Control IC | ns | 0.5663 | S5F |
| One-way ANOVA<br>(Tukey post hoc test) | T<sub>Reg</sub> Number | PBS vs. F5111 IC | * | 0.0217 | S5F |
| One-way ANOVA<br>(Tukey post hoc test) | T<sub>Reg</sub> Number | PBS vs. Control | ns | 0.8473 | S5F |
| One-way ANOVA<br>(Tukey post hoc test) | T<sub>Reg</sub> Number | Control IC vs. F5111 IC | ns | 0.1911 | S5F |
| One-way ANOVA<br>(Tukey post hoc test) | T<sub>Reg</sub> Number | Control IC vs. Control | ns | 0.9491 | S5F |
| One-way ANOVA<br>(Tukey post hoc test) | T<sub>Reg</sub> Number | F5111 IC vs. Control | ns | 0.0759 | S5F |
| One-way ANOVA<br>(Tukey post hoc test) | T<sub>Reg</sub>:T-bet<sup>+</sup>Tet<sup>+</sup>T<sub>Conv</sub> | PBS vs. Control IC | ns | 0.9540 | S5G |
| One-way ANOVA<br>(Tukey post hoc test) | T<sub>Reg</sub>:T-bet<sup>+</sup>Tet<sup>+</sup>T<sub>Conv</sub> | PBS vs. F5111 IC | * | 0.0305 | S5G |
| One-way ANOVA<br>(Tukey post hoc test) | T<sub>Reg</sub>:T-bet<sup>+</sup>Tet<sup>+</sup>T<sub>Conv</sub> | Control IC vs. F5111 IC | * | 0.0381 | S5G |
| One-way ANOVA<br>(Tukey post hoc test) | CD11a<sup>High</sup>Ki-67<sup>+</sup>T<sub>Reg</sub> Number | PBS vs. Control IC | ns | 0.3210 | S5H |
| One-way ANOVA<br>(Tukey post hoc test) | CD11a<sup>High</sup>Ki-67<sup>+</sup>T<sub>Reg</sub> Number | PBS vs. F5111 IC | ns | 0.0526 | S5H |
| One-way ANOVA<br>(Tukey post hoc test) | CD11a<sup>High</sup>Ki-67<sup>+</sup>T<sub>Reg</sub> Number | PBS vs. Control | ns | 0.9696 | S5H |
| One-way ANOVA<br>(Tukey post hoc test) | CD11a<sup>High</sup>Ki-67<sup>+</sup>T<sub>Reg</sub> Number | Control IC vs. F5111 IC | ns | 0.6700 | S5H |
| One-way ANOVA<br>(Tukey post hoc test) | CD11a<sup>High</sup>Ki-67<sup>+</sup>T<sub>Reg</sub> Number | Control IC vs. Control | ns | 0.5102 | S5H |
| One-way ANOVA<br>(Tukey post hoc test) | CD11a<sup>High</sup>Ki-67<sup>+</sup>T<sub>Reg</sub> Number | F5111 IC vs. Control | ns | 0.0909 | S5H |
| One-way ANOVA<br>(Tukey post hoc test) | T<sub>Reg</sub> IL-2R<sub>α</sub> MFI | PBS vs. Control IC | * | 0.0272 | S5I |
| One-way ANOVA<br>(Tukey post hoc test) | T<sub>Reg</sub> IL-2R<sub>α</sub> MFI | PBS vs. F5111 IC | **** | <0.0001 | S5I |
| One-way ANOVA<br>(Tukey post hoc test) | T<sub>Reg</sub> IL-2R<sub>α</sub> MFI | PBS vs. Control | ns | 0.8089 | S5I |
| One-way ANOVA<br>(Tukey post hoc test) | T<sub>Reg</sub> IL-2R<sub>α</sub> MFI | Control IC vs. F5111 IC | ** | 0.0025 | S5I |
| One-way ANOVA<br>(Tukey post hoc test) | T<sub>Reg</sub> IL-2R<sub>α</sub> MFI | Control IC vs. Control | ** | 0.0029 | S5I |
| One-way ANOVA<br>(Tukey post hoc test) | T<sub>Reg</sub> IL-2R<sub>α</sub> MFI | F5111 IC vs. Control | **** | <0.0001 | S5I |

**Table S7.** Flow cytometry antibodies and fluorescent dyes.

| Antigen | Fluorophore | Clone | Manufacturer | Catalog Number |
| --- | --- | --- | --- | --- |
| human IgG Fc | APC | HP6017 | BioLegend | 409306 |
| mouse/human phospho-STAT5 | AlexaFluor 647 | pY694 | BD Biosciences | 562076 |
| human CD3 | APC-eFluor780 | UCHT1 | ThermoFisher Scientific | 47-0038-42 |
| human CD4 | PerCP-Cy5.5 | SK3 | BD Biosciences | 341654 |
| human CD8 | BV605 | SK1 | BioLegend | 344742 |
| human IL-2Rα | BV421 | M-A251 | BD Biosciences | 562442 |
| human FOXP3 | PE | 236A/E7 | BD Biosciences | 560852 |
| human CD127 | AlexaFluor 488 | eBioRDR5 | ThermoFisher Scientific | 53-1278-42 |
| mouse CD3 | BV510 | 145-2C11 | BioLegend | 100353 |
| mouse CD4 | APC-eF780 | RM4-5 | ThermoFisher Scientific | 47-0042-82 |
| mouse CD8a | AlexaFluor 488 | 53-6.7 | BD Biosciences | 557668 |
| mouse IL-2Rα | PE | PC61.5 | ThermoFisher Scientific | 12-0251-83 |
| mouse FOXP3 | eFluor450 | FJK-16s | ThermoFisher Scientific | 48-5773-82 |
| LIVE/DEAD™ Fixable Blue | - | - | ThermoFisher Scientific | L34961 |
| human CD4 | PerCP-Cy5.5 | SK3 | BD Biosciences | 566316 |
| Fixable Viability Dye | eFluor750 | - | ThermoFisher Scientific | 65-0865-18 |
| mouse CD4 | eFluor450 | RM4-5 | ThermoFisher Scientific | 48-0042-82 |
| mouse CD49b | PE-Cy7 | DX5 | BioLegend | 108992 |
| mouse CD16/32 | - | 2.4G2 | BD Biosciences | 553142 |
| mouse FOXP3 | APC | FJK-16s | ThermoFisher Scientific | 17-5733-82 |
| mouse CD8a | BV570 | 53-6.7 | BioLegend | 100739 |
| mouse Ki-67 | AlexaFluor 488 | 16A8 | BioLegend | 652417 |
| mouse CD44 | PerCP-Cy5.5 | IM7 | ThermoFisher Scientific | 45-0441-82 |
| mouse/human Helios | PE/Dazzle 594 | 22F6 | BioLegend | 137231 |
| Violet Proliferation Dye 450 | - | - | BD Biosciences | 562158 |
| Zombie NIR Fixable Viability Kit | - | - | BioLegend | 423105 |
| mouse CD45 | APC-Cy7 | QA17A26 | BioLegend | 157617 |
| mouse TER-119 | APC-Cy7 | TER-119 | BioLegend | 116223 |
| human CD3 | BV605 | OKT3 | BioLegend | 317321 |
| human CD56 | PE | TULY56 | ThermoFisher Scientific | 12-0566-41 |
| human CD4 | PE-eFluor610 | RPA-T4 | ThermoFisher Scientific | 61-0049-42 |
| human CD8 | BV711 | SK1 | BioLegend | 344733 |
| human IL-2Rα | PE-Cy7 | M-A251 | BD Biosciences | 557741 |
| human FOXP3 | AlexaFluor 647 | 259D | BioLegend | 320214 |
| human Ki-67 | FITC | 20Raj1 | ThermoFisher Scientific | 11-5699-42 |
| mouse CD45 | PerCP-Cy5.5 | 30-F11 | BioLegend | 103132 |
| mouse CD3 | BV605 | 17A2 | BioLegend | 100237 |
| mouse CD4 | APC | GK1.5 | BioLegend | 100411 |
| mouse CD8a | BV786 | 53-6.7 | BD Biosciences | 563332 |
| mouse IL-2Rα | BV421 | 7D4 | BD Biosciences | 564571 |
| mouse FOXP3 | FITC | FJK-16s | ThermoFisher Scientific | 11-5773-82 |
| mouse NK-1.1 | PerCP-Cy5.5 | PK136 | BioLegend | 108727 |
| CellTrace™ Violet Cell Proliferation Kit | - | - | ThermoFisher Scientific | C34571 |
| mouse CD45.1 | APC-Cy7 | A20 | BioLegend | 110716 |
| mouse CD45.2 | PerCP-Cy5.5 | 104 | BioLegend | 109828 |
| mouse CD62L | PE | MEL-14 | BioLegend | 104408 |
| mouse CD44 | AlexaFluor 700 | IM7 | BioLegend | 103026 |
| Ghost Dye™ Violet 510 | - | - | Tonbo Biosciences | 13-0870-T100 |
| mouse CD16/32 | - | 2.4G2 | Bio X Cell | BP0307 |
| Rat IgG Isotype Control | - | - | ThermoFisher Scientific | 10700 |
| mouse CD4 | BUV563 | GK1.5 | BD Biosciences | 612923 |
| mouse CD8a | BUV615 | 53-6.7 | BD Biosciences | 613004 |
| mouse CD11a | BUV805 | 2D7 | BD Biosciences | 741919 |
| mouse IL-2Rα | BV785 | PC61 | BioLegend | 102051 |
| mouse CD27 | BV650 | LG.3A10 | BioLegend | 124233 |
| mouse CD44 | BV570 | 1M7 | BioLegend | 103037 |
| mouse CD69 | BUV737 | H1.2F3 | BD Biosciences | 612793 |
| mouse IL-2Rβ | BUV661 | TM-β1 | BD Biosciences | 741493 |
| mouse KLRG1 | BUV395 | 2F1 | BD Biosciences | 740279 |
| mouse ICOS | APC/Fire 750 | C398.4A | BioLegend | 313536 |
| mouse PD-1 | BV421 | 29F.1A12 | BioLegend | 135218 |
| mouse BCL-2 | AlexaFluor 647 | BCL/10C4 | BioLegend | 633510 |
| mouse CD3 | BV570 | 17A2 | BioLegend | 100249 |
| mouse CTLA-4 | APC-R700 | UC10-4F10-11 | BD Biosciences | 565778 |
| mouse FOXP3 | PE-Cy5.5 | FJK-16s | ThermoFisher Scientific | 35-5773-82 |
| mouse Ki-67 | AlexaFluor 488 | B56 | BD Biosciences | 558616 |
| mouse T-bet | PE-Cy5 | 4B10 | ThermoFisher Scientific | 15-5825-82 |
| mouse CXCR3 | BV650 | LG.3A10 | BioLegend | 124233 |
| mouse NK1.1 | BV711 | PK136 | BioLegend | 108245 |
| mouse NKp46 | BV605 | 29A1.4 | BioLegend | 137619 |
| Tetramer H2k(b) Tgd057 | PE | - | NIH | SVLAFRRL |
| Tetramer I-A(b) AS15 | PE | - | NIH | AVEIHRPVPGTAPPS |

